# Mapping Structural Aging of Human Tissue reveals tissue-specific trajectories and coordinated deterioration

**DOI:** 10.1101/2025.09.29.679316

**Authors:** Anamika Yadav, Kyle Alvarez, Kevin Y Yip, Eytan Ruppin, Jacqueline C Yano Maher, Veronica Lobo-Gomez, Caroline Kumsta, Sanju Sinha

**Affiliations:** Center for Data Science, Sanford Burnham Prebys Medical Discovery Institute, La Jolla, CA, USA; Cancer Data Science Lab, National Cancer Institute, National Institutes of Health, Bethesda, Maryland, 20894, USA; Pediatric and Adolescent Gynecology, Eunice Kennedy Shriver National Institute of Child Health and Human Development, Bethesda, Maryland, USA; Center for Cardiovascular and Muscular Diseases, Sanford Burnham Prebys Medical Discovery Institute, La Jolla, CA, USA

## Abstract

Tissue structure, the organization of cells, vasculature and extracellular matrix, underpins organ function. Yet how it deteriorates with age remains largely uncharacterized. Current aging research focuses primarily on molecular changes, missing this structural dimension. Here we present PathStAR (Pathology-based Structural Aging Rate), a computational framework that quantifies tissue structural aging from histopathology images without training on chronological age. Applying PathStAR to 25,306 postmortem biopsies spanning 40 tissue types from 970 individuals aged 21–70, we show that structural aging unfolds through non-linear phases rather than gradual decline. We identify three distinct temporal programs: early-aging tissues (vascular system, peaking in the 30s), late-aging tissues (uterus and vagina, peaking around menopause) and biphasic-aging tissues (digestive and male reproductive organs, with two acceleration periods). During accelerated phases, tissues exhibit common molecular signature of increased inflammation, decreased energy production, regeneration and quality control alongside tissue-specific pathway disruptions related to its function: Artery-specific decline of peroxisomal function, responsible for fatty acid breakdown and testis-specific decline of spermatogenesis. Cross-organ analysis reveals coordinated deterioration within individuals, not only within expected organ systems but also unexpectedly between digestive and reproductive tissues, traced to shared sex hormone signaling. This reveals a role for sex hormones in maintaining structural integrity of non-reproductive organs during aging. Genome-wide association analysis identifies 123 germline variants associated with organ-specific accelerated structural aging, including variants in the longevity regulator SIRT6 linked specifically to vascular structural decline. As proof of concept, PathStAR captures the established non-linear functional decline of the ovary, fertility loss in the 30s and menopause in the 50s, that bulk transcriptomic and methylation profiles from matched samples fail to detect. PathStAR provides a systematic map of structural aging across the human body: when tissues structure decline, what molecular programs define each phase, and how organs deteriorate in coordination.

## 2. Introduction

Aging is the biggest risk factor for most chronic diseases, including cardiovascular disease, Type 2 diabetes, Alzheimer’s disease, and many cancers [1]. Current aging research has advanced our understanding of molecular changes during aging, both cell intrinsic like telomeres shortening, cell senescence, and autophagy, as well as systemic pathways like the insulin/IGF-1 axis [1, 2, 3, 4]. Aging clocks, either from methylation or other omics layers, enabled improved prediction of risk of mortality and diseases [5, 6]. Complementing these clocks, recent studies showed that molecular changes during aging unfold through discrete, non-linear transitions rather than gradual decline, in blood multi-omics [7] as well as in tissues proteomics [8].

While we have advanced our knowledge of molecular, cellular and systemic changes during aging, a critical missing gap remains in understanding how the physical structure of tissues, the spatial organization of cells, vasculature and extracellular matrix, and the basis of their function, deteriorates with aging (*Structural Aging*). Our current understanding of tissue structural aging comes from manual inspection of histology images revealing both global deterioration patterns like fiber fragmentation [9,10], atrophy [11], and fibrosis [12], and, tissue-specific patterns like ovarian follicle loss [13], bone porosity [14], brain cortical thinning and demyelination, muscle atrophy [15], arterial stiffening [16], heart fibrosis [17,18], thymic involution [19], and lens clouding [20]. These structural changes precede both decline of organ function with age and diseases, e.g. ovarian follicle loss reducing fertility and cardiac fibrosis increasing heart failure risk. However, despite these specific examples, currently there is no systematic study to capture generalizable tissue structure changes during aging. This gap has persisted due to lack of large-scale high-resolution tissue images across the body and computational tools to quantify these tissue sub-structures.

Recent studies have applied deep-learning models to histopathology images[21,22] to predict chronological age, effectively constructing morphology-based aging clocks, analogous to methylation or transcriptomic clocks. While these approaches demonstrate that tissue architecture contains age-predictive information, they are optimized for age estimation rather than for characterizing when and how structural changes unfold.

These limitations have given rise to several fundamental open questions about aging: <u>When and how do different tissue physical structures deteriorate during aging? What are their molecular drivers? Are there tissue-specific periods of accelerated structural aging, and what molecular dysregulation drives this acceleration? Which organs are early-agers versus late-agers? Do organs age independently or in co-ordination? How do clinical factors accelerate or protect against structural aging? And finally, how do germline variants affect organ-specific structural aging?</u>

We present the first framework that tries to answer these questions: PathStAR (Pathology-based Structural Aging Rate), a computational framework that quantifies tissue structural aging from histopathology images without training on chronological age, and applied it to 25,306 post-mortem biopsies across 40 tissue types from 970 individuals (ages 21–70) in the Genotype-Tissue Expression (GTEx) cohort [23]. As proof-of-concept, PathStAR captures the established non-linear ovarian functional decline: fertility loss in the 30s and menopause in the 50s, that bulk transcriptomic and methylation profiles from matched samples fail to detect. Across the body, we identify three distinct temporal aging programs, coordinated cross-organ structural deterioration, molecular drivers of accelerated structural aging periods, and germline variants linked to organ-specific aging trajectories. Our framework establishes structural aging as a quantifiable dimension of the aging process, complementing molecular approaches. We define “structural aging” as age-associated shifts in tissue architecture comprising degenerative processes, adaptive remodeling, or neutral architectural variation. Our objective is to quantify how this unfolds systematically at a population level.

## 3. Result

### 3.1 Overview of PathStAR Pipeline

Based on the notion that histology images capture architecture features critical for tissue function, we developed a computational framework called *PathStAR* (Figure 1A-C), that quantify how and when this tissue structure changes during aging from high-resolution H&E images of large-scale post-mortem biopsies.

**Fig 1:**
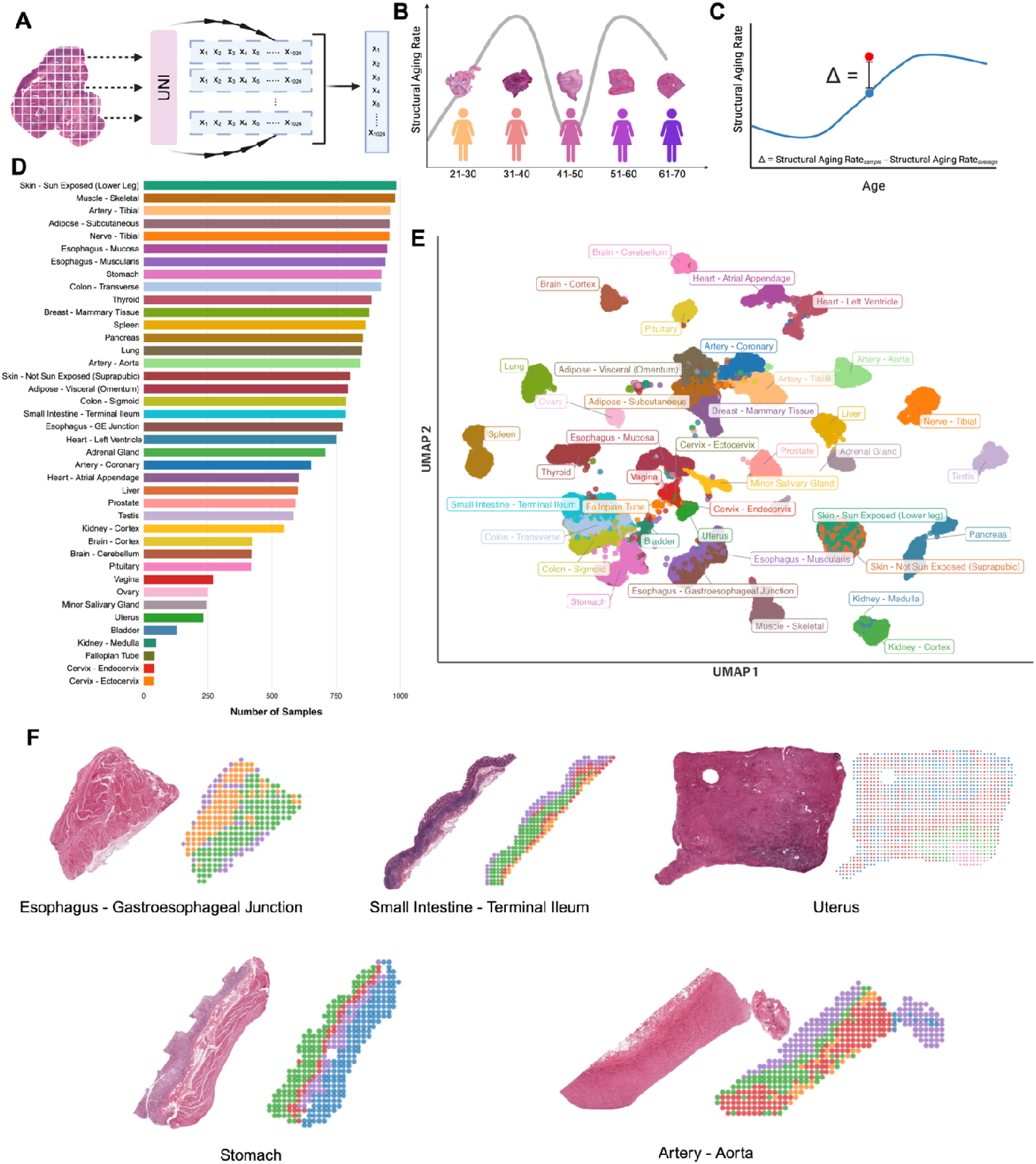
Overview of PathStAR pipeline Steps and GTEx cohort used here: **(A-C)** PathStAR start (Step 1: Feature extraction) by extracting morphology feature of 256*256 pixels patch-level using a pre-trained pathology foundation model (UNI) which are aggregated (mean pooling) to generate slide-level representation. It then computes structural aging rate by quantifying the rate of morphological change per year using sliding 10-year windows, generating age-specific trajectories that reveal periods of Accelerated Structural Aging periods. Finally, it computes individual delta-structural aging scores (red) measuring each sample’s deviation from the population aging trajectory (blue). **(D)** Number of samples available for this study per tissue type. **(E)** UMAP based on whole-slide level representation cluster different tissue types, demonstrating its ability to capture tissue type-specific morphological properties. **(F)** Morphological patch level features from UNI clustered using spatial information and K-means, where different colors are different clusters.

PathStAR was applied on (GTEx cohort) post-mortem biopsies from 970 non-diseased individuals (age range = 21-70 years, 66.4% males, **Figure S1**) spanning 40 tissue types (**Figure 1D**). GTEx provided a total of 25,306 biopsy whole-slide scans (×20 magnification), from which we extracted 30.3 million patch images (512 × 512 pixels). Pathology annotations from GTEx for these images confirmed canonical aging-related increase in atrophy, fibrosis, cysts, atherosclerosis, and decrease in ovum, spermatogenesis, and of protective layers including mucosa, endo-and myometrium (**Figure S2**), demonstrating quality to develop a systematic method. Tissue-specific quality control analysis is presented in **Notes S1**.

Briefly, PathStAR consists of **three steps** (Figure 1A–C) where it first extracts structural features from the image, builds a structural aging trajectory based on it, and then finally, compute individual-level deviation from the trajectory. This is provided in detail in Methods 5.3, and we here briefly describe the three steps.

1. ***<u>Step 1: Feature extraction from whole-slide images</u>*:** Each whole-slide image is segmented into patches after background removal, and each patch is encoded into a 1024-dimensional embedding using the UNI vision encoder [24], a vision transformer pretrained on 100,000 whole-slide images (Methods 5.3). Patch embeddings are aggregated via mean pooling to produce a single slide-level representation per sample (Figure 1A). These representations capture tissue-specific morphological patterns, readily separating distinct tissues and substructures in low-dimensional space (Figure 1E–F). We tested a few others diversely trained and broadly adopted feature extractors to ensure our framework results are not artifacts of one encoder (**Notes S2**).
2. ***<u>Step 2: Construction of structural aging trajectories</u>*:** We next asked: when does tissue structure change most rapidly during aging? For each tissue, we quantify the rate of structural change per year: the Structural Aging Rate, by comparing morphological representations between consecutive 10-year age windows using an effect-size-based measure. Sliding this window forward in one-year increments produces a continuous, population-level structural aging trajectory spanning approximately ages 30–60. This trajectory reveals periods where tissue structure changes rapidly versus remains stable; we define intervals with pronounced acceleration as **A**ccelerated **S**tructural **A**ging (**ASA**) periods.
3. **<u>Step 3: Individual-level deviation from population trajectories</u>**: With population-level trajectories established, we next ask: which individuals are aging faster or slower than expected for their tissue and age? For each sample, we compute an individual structural aging score: a measure of how much a given tissue in a given individual deviates from its expected aging path (Methods 5.3, delta-structural aging score, Figure 1C). These scores enable identification of individuals with accelerated or protected structural aging, and downstream analysis of the clinical, lifestyle, and genetic factors that drive these deviations.

### 3.2 PathStAR captures ovary’s functional decline and its molecular correlates without chronological age training

#### 3.2.1 Morphology Captures Ovary’s Non-Linear Functional Decline without supervision

Ovary exhibits a unique functional decline: fertility decreases gradually from the early 30s, marked by rapid follicular depletion that accelerates through the late 30s, and culminates in menopause at approximately 50–55 years of age [25, 26]. Current molecular clocks, trained on chronological age, assume linear aging, and thus cannot capture this non-linear ovary remodeling (**Figure S3).** We thus tested whether PathStAR can capture this established non-linear functional decline.

We first observed that patch-level UNI-features can spatially segregate distinct ovary substructures (**Figure 2A-B, Figure S4**). We then find that these morphology features readily distinguished young, middle-aged and old samples: the first two principal components correlated with age (r² = 0.43), indicating substantial age-related structural change (**Figure 2C**). We then used PathStAR to compute the structural aging rate trajectory using 250 ovary biopsies (age range: 21-70 years, **Figure 2F**). This yielded a striking bimodal pattern with two peaks of accelerated structural aging aligning with established reproductive milestones (**Figure 2F**, Methods 5.3). The first **A**ccelerated **S**tructural **A**ging (*ASA*) period peaks at ages 35-40, coinciding with fertility decline, while the second ASA period peaks at ages 55-60, corresponding to menopause. Importantly, ischemic time and other post-mortem artifacts showed minimal correlation with this trajectory (**Notes S3, Figure S5**).

**Figure 2:**
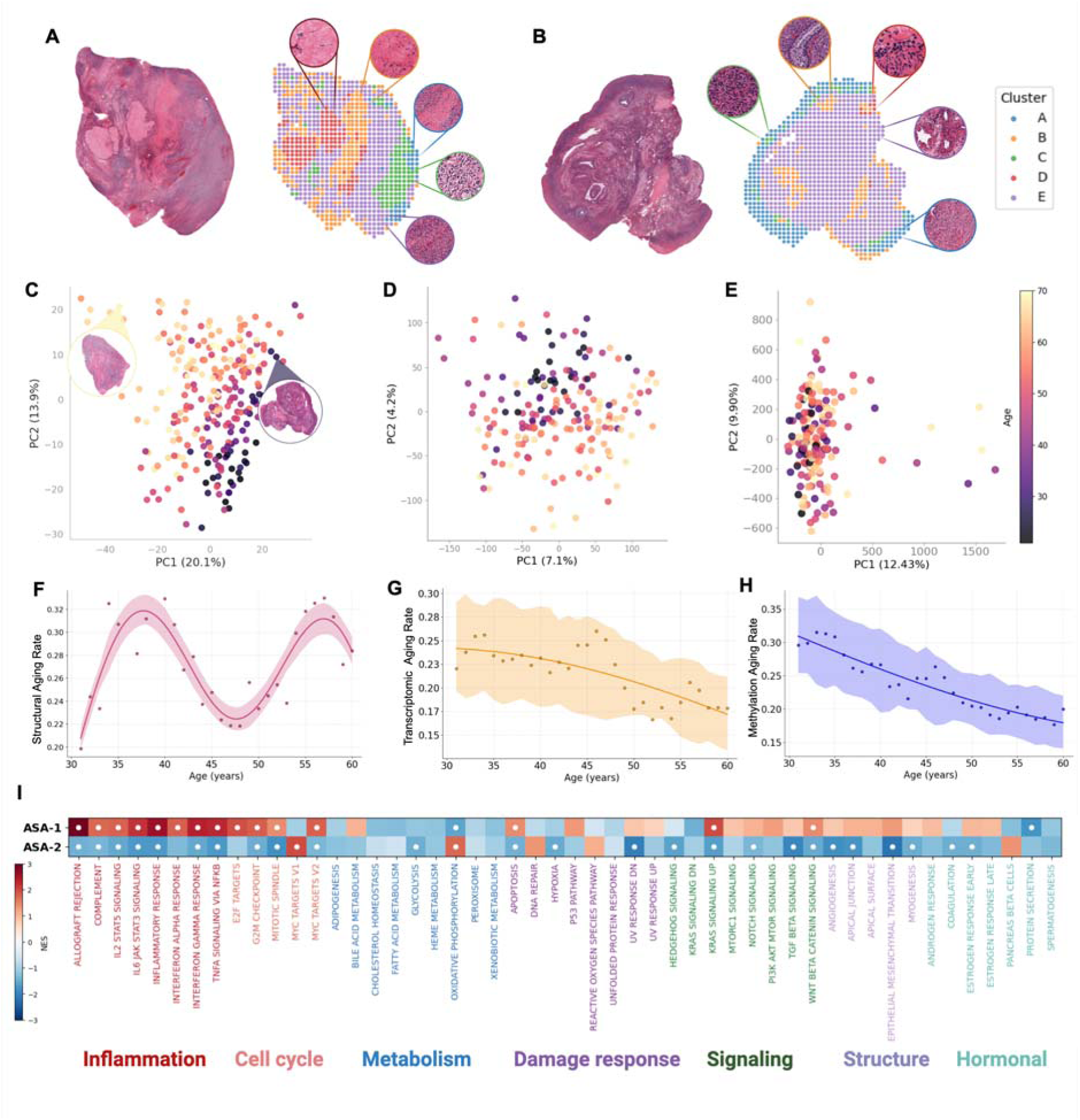
Morphology features capture non-linear ovary structural aging during aging that bulk-molecular profiles cannot. **(A-B)** The morphological patch level features from 2 ovary samples from both older (left, age 60-69) and younger (right, age 20-29). The cluster shows that UNI can very well capture the structural property of the tissue. **(C-E)** Principal component analysis of all morphology features extracted by UNI in 250 ovary samples from whole slide images separate young vs. old samples, without prior training on chronological age; expression and methylation profiles from the matched samples respectively. **(F)** PathStAR applied to UNI features from 250 ovarian samples, age 21-70 years, captures non-linear structural aging. Structural aging rate was estimated using sliding 10-year age windows (e.g., 21–30 vs. 31–40), advanced in 1-year increments across the age span. Each data point represents the structural difference between two adjacent windows and is positioned at the upper bound of the younger window. Windows containing insufficient donor representation were excluded, yielding 28 retained trajectory points derived from the full cohort. Smoothing was applied on points with > 5% of UNI features with significant Structural Aging Rate (p < 0.05) yielding 27 datapoints. **(G)** Expression profiles can segregate young vs. old samples but fail to capture the bimodal pattern in structural aging trajectory of ovary. **(H)** Methylation profiles fail to capture young vs. old samples as well as doesn’t reflect the bimodal pattern captured by structural aging trajectory. **(I)** Heatmap showing pathway enrichment in ASA 1 and ASA 2 compared to their controls. white dots denote statistically significant enrichment (P < 0.05).

In comparison, when this exact <u>unsupervised trajectory analysis is applied to bulk-expression and methylation profiles (**Figure 2D-E**) from the same samples, neither expression nor methylation trajectory recapitulated the bimodal pattern of structural trajectory</u> (**Figure 2 G, H, Methods 5.4**). Finally, we interpret the structural changes during the two ASA periods, using a pathology vision-language model (PLIP [27], QC testing in **Notes S3, Figure S6**), revealing increased fibrosis and atrophy during both ASA periods, with pronounced decline in ovum abundance during ASA-2 (**Figure S7**), consistent with the known structural changes of follicular depletion and post-menopausal involution. Specific morphological feature groups showing distinct age-related patterns (increasing, decreasing, or stable) are described in **Notes S4** (**Figure S8-9, Methods 5.5**).

In summary, this non-linear trajectory emerged from histology without any training on chronological age, a capability that neither unsupervised molecular analysis nor linear-by-design molecular clocks can provide. We next identified the molecular changes accompanying each structural transition.

#### 3.2.2 Molecular Changes During Accelerated Structural Aging Periods of Ovary

Having established the two ASA periods from histology, we next asked what molecular changes accompany each structural transition. We compared bulk-expression and methylation profiles from each ASA period to its preceding structurally stable phase (**Figure 2I**): ASA-1 (ages 35–40) vs. baseline (ages 25–30), and ASA-2 (ages 55–60) vs. baseline (ages 45–50). For each contrast, we performed differential expression analysis between age groups, ranked genes by signed effect size and statistical significance, and conducted pathway enrichment analysis on the resulting ranked lists to identify coordinated biological programs associated with each structural transition.

ASA-1 is marked by upregulation of multiple inflammatory pathways (TNF-alpha via NFKB, IL2-STAT5, and others; **Figure 2I**, **Figure S10**), alongside hypermethylation-mediated silencing of Notch signaling, a master regulator of granulosa cell proliferation [28, 29], and ERK1/2, which governs both FSH and LH signaling [30, 31] (**Figure S10**). Together, these changes suggest a pro-inflammatory microenvironment coinciding with rapid loss of ovarian reserve [32, 33, 34, 35, 36]. In contrast, ASA-2 is uniquely characterized by downregulation of growth factor signaling (TGF-Beta), EMT, and cell cycle pathways, consistent with loss of follicular activity and reduced steroidogenesis in the post-menopausal period [37, 38, 39] (**Figure 2I**). These period-specific molecular programs were identifiable only because PathStAR defined the structural windows from histology, establishing the foundation for whole-body analysis described next.

### 3.3 Structural Aging dynamics in different tissues across the whole human body

#### 3.3.1 Morphological Features Segregate Young vs. Old Tissues for Many Tissues Across Body

We next applied PathStAR across 40 human tissues with available H&E images (N = 970 individuals; N = 25,306 biopsies; **Methods 5.1**). Morphology-based PCA readily separated young from old samples in many tissues without any training on age (**Figure 3**), indicating that aging, not individual variation or batch effects, is the dominant driver of structural change. To quantify this separation, we measured both the age-related variance captured by PC1–2 (cross-validated OLS, **Table S1**) and the ability to distinguish young (21–30) from old (60–69) samples (cross-validated LDA, **Table S2**). We chose fifteen tissues that where age is the dominant driver of structural change (r² > 0.15 and AUC > 0.8), at least 200 total samples with samples distributed across whole age span, for calculation of structural aging trajectory.

**Figure 3:**
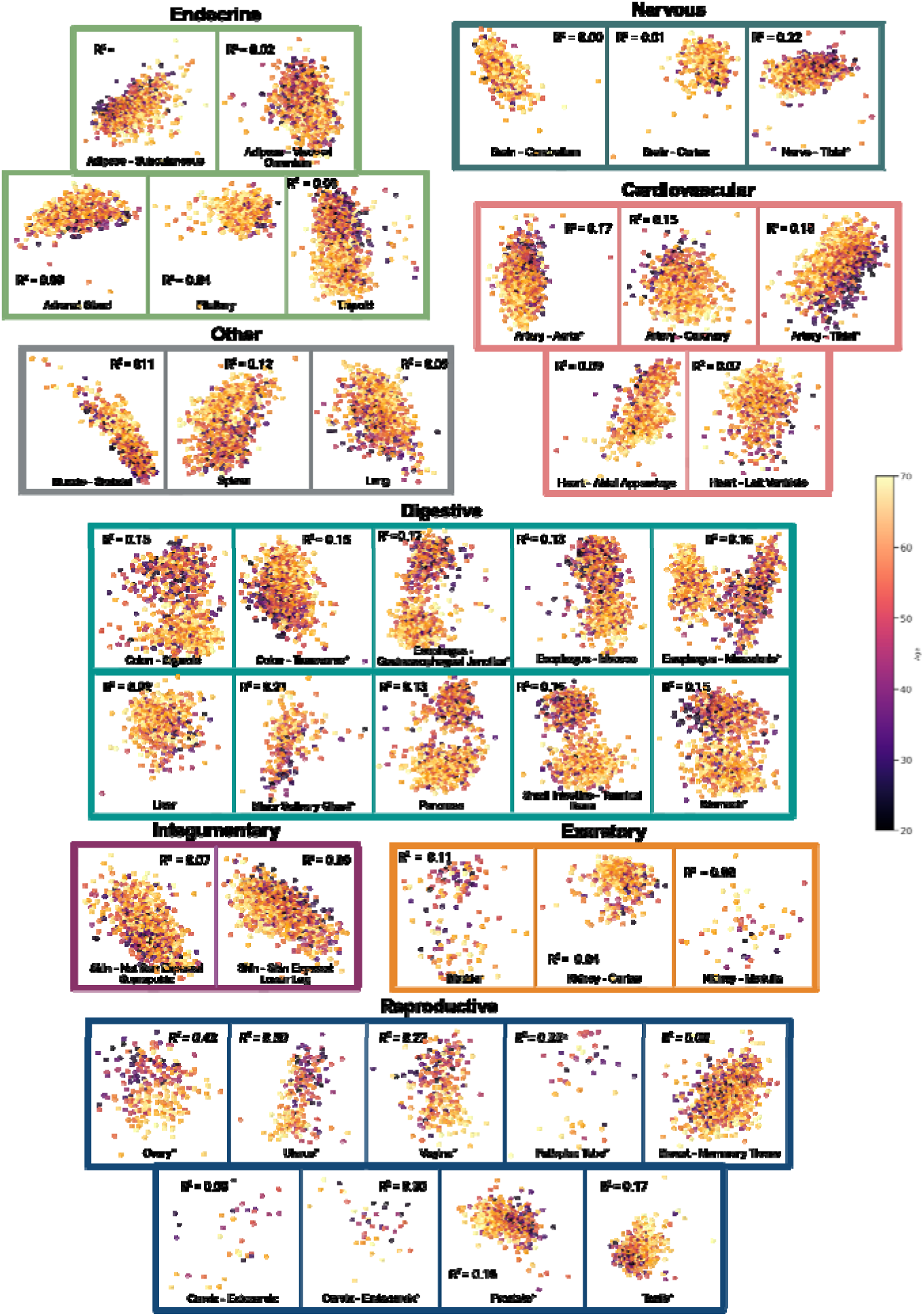
Tissue morphology naturally segregates young vs. old tissues across multiple tissue in human body: Morphology features-based PCA for 40 tissue types, where darker color represents younger tissue and light colors represents older tissue(**Table S1**). Tissues are grouped into their respective organ system outlined in an organ-system specific color. Stars in the name of the tissue represent significant tissues that separate old, middle and young samples without pre-training.

#### 3.3.2 Human Tissues Age Through at-least Three Distinct Temporal Programs

We generated structural aging rate trajectories for each tissue, selecting only tissues that showed clear morphological differences between young and old samples, indicating detectable structural aging above individual variation. We next filtered and focused on tissue types with high-confidence trajectory (**Methods 5.2, Notes S6**). This yielded a total of 15 tissues comprising vascular, digestive, and reproductive tissues (including ovaries), where we generated remodeling trajectories (**Figure 4A-C**). For transparency, we provide all Raw Data Plots (**Figures R1-R9**) showing individual data points, sample distributions, and sex-stratified analyses.

**Figure 4:**
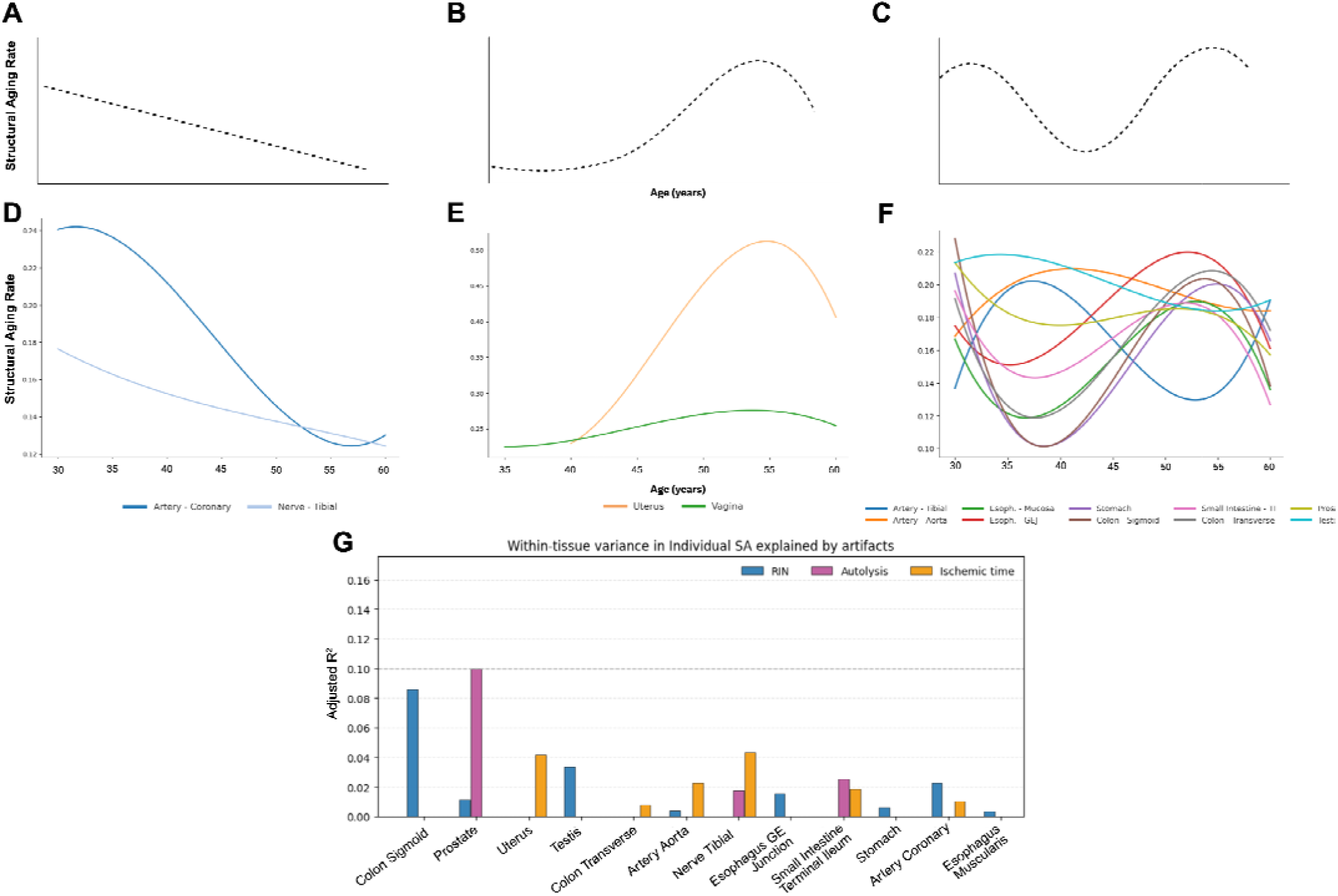
Overview of Structural Aging Trajectories across 14 tissues. **(A-C)** Three distinct aging patterns identified: Early Agers peak in the 30s then decline, Late Agers remain stable until major changes in the 50s-60s, and Biphasic Agers (the predominant pattern in 9/14 tissues) exhibit two acceleration periods around the early 30s and 50s. **(D-F)** Tissue-specific examples showing vascular and neural tissues as Early Agers, female reproductive tissues as Late Agers, and digestive tract plus male reproductive organs as Biphasic Agers, revealing coordinated aging within functionally related systems. **(G)** Within-tissue variance in individual structural age (SA) explained by technical artifacts (RIN, autolysis, ischemic time); except prostate, other tissues show minimal artifact-driven variance.

Structural aging trajectories across these 15 tissues revealed tissue-specific patterns that could be clustered into three groups (**Figure 4A-C, Methods 5.2**). <u>Early-Aging</u> <u>Tissues</u> exhibit high structural aging rates in the 30s where this rate declines later, experiencing high structural changes during young adulthood (**Figure 4A**). <u>Late-Aging</u> <u>Tissues</u> remain structurally stable through early adulthood, then undergo major structural changes in later decades (**Figure 4B**). <u>Biphasic-Aging Tissues</u> display two distinct periods of accelerated structural aging separated by stable phases, like the ovary pattern (**Figure 4C**). Notably, some tissues exhibited markedly higher structural aging rates than others, indicating greater cumulative remodeling over time.

The vascular system, tibial and coronary arteries, showed Early Structural Aging, with structural aging highest in the 30s (**Figure 4D, F**; *Raw points, # of samples/age group & confidence intervals in* **Figure S12**, *Sex-stratified plot in* **R1**). This trajectory is consistent with proteomic studies that also report early cardiovascular aging [40]. The trajectory remains consistent even after filtering out individuals with systemic diseases (**Figure R2**). Pathology notes support these findings: 1) Individuals with higher delta-structural aging score is more likely to have atherosclerosis (**Figure S13**); 2) Atherosclerotic changes, early plaque formation in arteries, showed their steepest increase during the 30s vs. 20s, then plateaued in later decades (**Figure S2**, top-left panel). Interestingly, tibial nerve also displays Early Structural Aging, an unexpected finding warranting further investigation. Also, an overall higher structural aging rates of coronary artery vs. tibial nerve indicate higher cumulative remodeling during aging.

In contrast to the above, uterus and vagina exhibited Late Structural Aging pattern, with their highest structural aging rates occurring in early and middle 50s **Figure 4E**), aligning with menopause and post-menopause. This likely reflects major changes in uterus including substantial atrophy, endometrial thinning [41], and myometrial structural changes following estrogen withdrawal [42] during menopause and post-menopause [43]. Accelerated structural aging periods of vagina and uterus aligns with ASA-2 of ovary, suggesting coordinated reproductive aging, potentially driven by shared hormonal regulation. We tested this coordination aging at individual-level in next section.

The predominant pattern observed in our dataset was Biphasic Structural Aging, as earlier described for the ovary (**Figure 2F**), characterized by two distinct peaks of structural aging rates, indicating two periods of accelerated structural aging (**Figure 4C**). While the timing and structural aging rates of two peaks differed among tissue types, this pattern was exhibited by 9 of the 14 tissues examined (aside of ovary), including digestive tract (esophagus, stomach, colon, small intestine, salivary gland), male reproductive organs (prostate, testis), and artery-tibial (**Figure 4F**). Overall, the two major periods of accelerated structural aging for these tissues were in the 30s and around the 50s. Beyond sharing similar population-level trajectory, we explored whether digestive tract and male reproductive organs show coordinated structural aging at individual level in next section. Testing contribution of post-mortem artifacts towards these patterns, we find the total variance explained of structural aging rates by all post-mortem artifacts to be minimal in almost all tissues (<10%, **Figure 4G, Table S5-6)**.

Finally, the above trajectories <u>do not</u> emerge from molecular age clocks trained on chronological age, which assume linearity by design, nor when applying the same trajectory analysis to bulk expression or methylation profiles (**Figure S14A-B, S15, R4-5**). <u>Overall, this shows that structural aging of human tissues occurs in nonlinear</u> <u>phases, with distinct patterns across organ systems: vascular tissues age early (30s),</u> <u>female reproductive tissues coordinate with ovary’s late peak (50s), and digestive and</u> <u>male reproductive organs follow a biphasic pattern.</u> Trajectories of 20 additional tissues, with at least 200 cases, which were excluded from the main analysis due to not meeting all inclusion criteria, are shown in **Figure R3** and their molecular trajectory are shown in **Figure R4-5**. Repeating the analysis using alternative pathology foundation models (UNI v2, CONCH v1.5[44], and Virchow2[45]) reproduced the overall similar structural patterns across most tissues (**Figure R6–R8; Supplementary Note 2**), indicating that the observed nonlinear remodeling dynamics are robust to encoder choice.

### 3.4 Molecular Programs During Accelerated Structural Aging

We next identified molecular changes associated with tissue-specific Accelerated Structural Aging (ASA) periods. ASA intervals were first defined from structural aging trajectories as peak inflection periods unique to each tissue. For each ASA interval (heuristically defined as a 5-year window based on peak width), we compared bulk transcriptomic profiles to those from the immediately preceding 5-year structurally stable window within the same tissue. Differential expression and pathway enrichment analyses were then performed for each ASA–stable transition across tissues (complete pathway results in **Table S8; Fig. S10**).

#### 3.4.1 A global inflammatory-metabolic program with organ-specific points of failure

Across tissues, accelerated structural aging periods were defined by a common molecular signature (**Figure 5A**): inflammatory pathways (TNF-alpha signaling, interferon alpha and gamma response, and complement) were broadly upregulated, while energy production (oxidative phosphorylation), proliferative capacity (E2F targets, G2M checkpoint, MYC targets), and cellular quality control (unfolded protein response, DNA repair, mTORC1 signaling) were simultaneously suppressed. This convergent program: inflammatory activation coinciding with the decline of regenerative, metabolic, and quality control, was the dominant molecular feature of structural aging acceleration across the body. Finding pathways that simply linearly increase or decrease with age cannot fully capture this combination of activation/inactivation (**Figure S16**).

**Figure 5.**
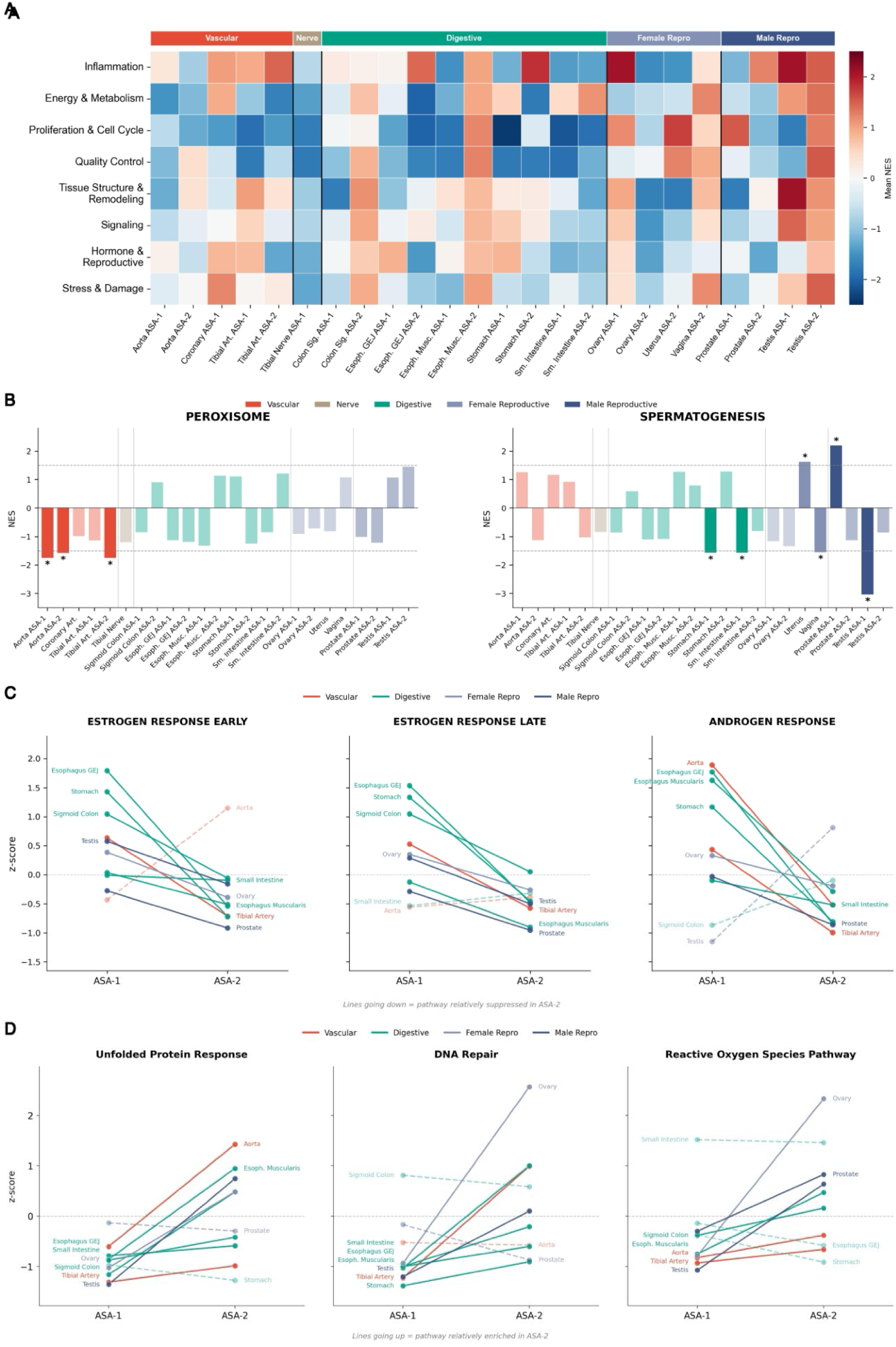
Molecular programs underlying accelerated structural aging. **(A)** Mean NES of 50 Hallmark pathways grouped into eight categories across 24 tissue-ASA combinations, organized by organ system. **(B)** Tissue-level NES for Peroxisome (significant exclusively in arteries) and Spermatogenesis (strongest in testis, NES = − 3.04). **(C)** Z-scored NES from ASA-1 to ASA-2 for three hormone pathways, each declining in 8-9 of 10 tissues. **(D)** Opposing pattern for three damage-response pathways, each increasing in 7-8 of 10 tissues. Solid lines indicate dominant direction; dashed lines indicate exceptions.

Within this shared program, individual organ systems harbored distinct molecular vulnerabilities (**Figure 5B, Figure S17**). In vascular tissues, peroxisome pathway activity, responsible for fatty acid oxidation, plasmalogen synthesis, and reactive oxygen species detoxification, declined exclusively in the aorta and tibial artery and in no other tissue (**Figure 5B**). This artery-specific loss of peroxisomal function may compound the early cardiovascular aging observed in their structural trajectories (**Figure 4A**), as peroxisomal failure drives lipid accumulation and oxidative damage in the vessel wall. At the other extreme, testis exhibited the most dramatic molecular response of any tissue: upregulated inflammatory and immune pathways, while spermatogenesis was simultaneously suppressed to the strongest degree in any tissue (**Figure 5B**, **Figure S18**). This pattern is consistent with immune-mediated germ cell destruction following blood-testis barrier breakdown. This suggests an inflammatory catastrophe rather than a hormonal deficiency. We note that finding pathways that linearly increase/decrease with chronological age cannot find above-noted pathways (**Figure S16**).

#### 3.4.2 A molecular phase transition between ASAs: from hormone signaling to damage response

What distinguishes the first period of acceleration from the second? To answer this, we compared the relative pathway enrichment profiles between ASA-1 and ASA-2. This comparison revealed a consistent temporal transition across two classes of pathways. First, all three hormone-responsive pathways, estrogen response early, estrogen response late, and androgen response, declined between ASA-1 and ASA-2 in nearly every tissue examined (9/10, 8/10, and 8/10 tissues, respectively; **Figure 5C**), indicating a broad loss of hormonal signaling capacity as tissues enter their later period of structural aging. Notably, this decline was not driven by reproductive organs. The steepest drops occurred in gastrointestinal tissues, particularly the esophagus gastroesophageal junction and stomach, consistent with the known expression of estrogen receptors throughout the digestive epithelium and their role in mucosal barrier maintenance. Second, and in the opposite direction, the damage-response pathways increased between ASA-1 and ASA-2 in the majority of tissues: the unfolded protein response (8/10 tissues), reflecting rising proteotoxic stress from misfolded protein accumulation; DNA repair (7/10), reflecting increasing genotoxic burden; and the reactive oxygen species pathway (7/10), reflecting escalating oxidative damage (**Figure 5D, Figure S19**). Together, these indicate that the second period of acceleration is defined by high damage response across tissues.

Overall, we above provide molecular program, either driving or characterizing, critical structural aging periods of organs, which will inform anti-aging therapies to protect function in organ-defined and time-resolved (when and how long to treat) manner.

### 3.5 Cross-organ coordination of structural aging within individuals

We next asked whether individuals showing accelerated structural aging in one tissue also exhibit accelerated aging in related tissues: Coordinated deterioration within the same person. PathStAR computes individual level delta-Structural Aging scores (ΔSA, **Table S7**) denoting individual’s deviation from its population-level trajectory (**Method 5.3**). To identify coordinated aging, we computed correlations among ΔSA scores of the 15 tissues with high-confidence trajectories (**Figure 6, Method 5.7**). We first observed that tissues within the same organ system showed significant positive correlations, indicating that individuals who age faster in one tissue tend to age faster in related tissues: gastrointestinal (colon, esophagus, stomach; r = 0.20–0.43, **Figure 6A**, *Box I*), female reproductive (uterus-vagina; r = 0.48, *Box II*), and vascular (tibial, coronary, aorta; r = 0.14–0.16, *Box III*), with additional brain (cerebellum-cortex: r = 0.55 males, 0.40 females) and cardiac coordination across all 40 tissues (**Figure S20**).

**Figure 6:**
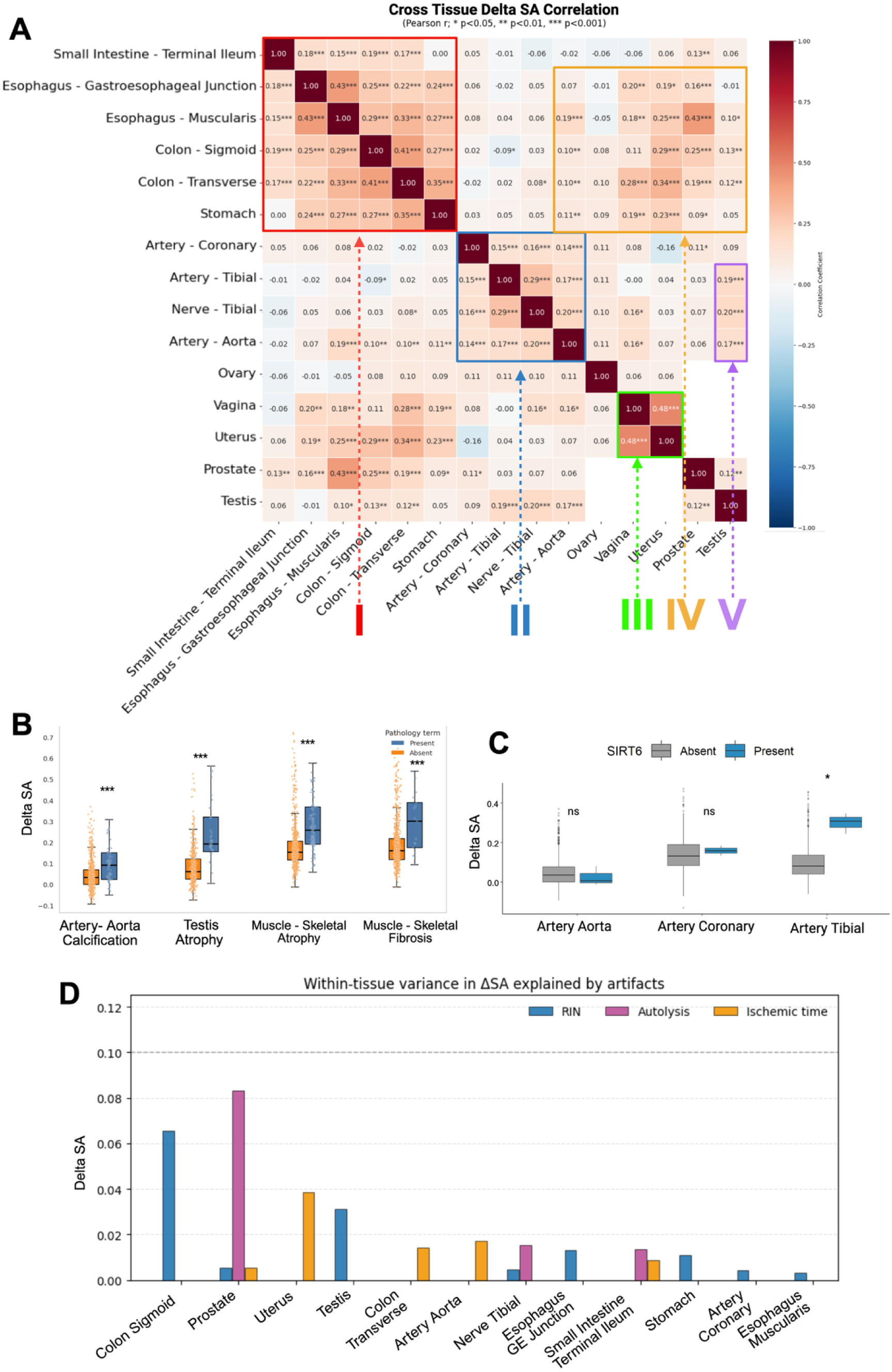
Cross-tissue Delta-Structural Aging score correlations reveal coordinated structural aging within individuals. **(A)**. Heatmap showing Pearson correlation coefficients between individual Delta-Structural Aging (SA) scores across 15 tissue types from the same individuals (n=970). Delta-SA scores quantify each individual’s deviation from expected structural aging trajectories for each tissue. Color scale represents correlation strength: dark red indicates strong positive correlations (r≥0.4), light red/orange indicates moderate positive correlations (r=0.2-0.4), white indicates weak correlations (r<0.2), and blue indicates negative correlations. Strong positive correlations reveal coordinated aging between tissue pairs, where individuals with accelerated structural aging in one tissue consistently exhibit accelerated aging in the partner tissue, suggesting synchronized deterioration driven by shared mechanisms. The p-values reported are before FDR-correction. (**B)** Box plot showing the distribution of delta SA score in calcification present and absent in artery aorta, atrophy present and absent in testis and muscle, fibrosis in muscle (p value for *** < 0.001) **(C)** Distribution of Delta SA score in Artery- Aorta, coronary and tibial with SIRT6 present and absent. p value for * < 0.05) **(D)** Within-tissue variance in ΔSA across age groups attributable to artifacts. The variance explained is uniformly low, suggesting remodeling dynamics are not driven by technical confounders.

Beyond within-system coordination, we observed an unexpected cross-system coordinated deterioration axis between digestive and reproductive tissues. Individuals who showed accelerated structural aging in colon and esophagus also showed accelerated aging in prostate (r = 0.14–0.43, **Figure 6A**, *Box IV*). Aligning with this, at the population level, these same tissues share biphasic structural aging trajectories (**Figure 4**). Further corroborating this, at the molecular level, gastrointestinal tissues showed the highest enrichment of hormone-responsive pathways (estrogen and androgen response) during their first period of acceleration, even higher than reproductive tissues (**Figure 5A, C**). Overall, this suggests that structural aging of digestive and reproductive tissues may be linked through a hormonal signaling axis whose decline drives synchronized structural deterioration across organ system boundaries. Additional cross-system axes (**Figure S20**) emerged that lacked an obvious shared molecular program but were nonetheless robust: neurovascular coordination (tibial nerve with multiple arteries, r = 0.23–0.28) and sex-specific neuroendocrine coordination (pituitary-brain correlations stronger in males than females). The mechanisms driving these axes remain to be determined.

### 3.6 Clinical and Genetic Determinants of Organ-Specific Structural Aging

To ask whether accelerated structural aging has identifiable clinical and genetic drivers, we tested associations between individual level ΔSA scores and clinical health records, lifestyle factors, and germline variants. Clinically, accelerated structural aging showed strong positive associations with canonical aging pathologies (**Figure 6B**, **Figure S21**), including atrophy (broad associations across tissues, particularly gastrointestinal and reproductive systems), fibrosis, vascular calcification, and hyalinization. Systemic diseases including autoimmune conditions (Lupus) and neurological diseases (Dementia) showed positive associations across multiple tissues (**Figure S21**), confirming that individuals with these conditions exhibit globally accelerated structural aging. Lifestyle variables had insufficient sample sizes to reach significance (**Figure S22**).

We next performed a genome-wide association analysis between germline variants of 970 individuals and ΔSA scores, focusing on 127K functional variants filtered from 998K starting variants (Methods 5.9). A total of 123 genes showed significant associations (FDR p < 0.1**, Figure S23, S19, Table S9**). The pathway enrichment of these hits directly paralleled the molecular programs identified in Section 3.4: detrimental variants that accelerate structural aging were enriched in oxidative phosphorylation (**Figure S24**), aligning with the same energy production programs that declined universally during ASA periods (Figure 5A). Conversely, protective variants were enriched in immune signaling pathways, particularly IL17 and interferon signaling, suggesting that genetic capacity to regulate the inflammatory response, the dominant upregulated program during structural aging acceleration, is protective against it.

The most striking individual association: SIRT6 variants were specifically associated with accelerated vascular structural aging (**Figure 6C**). SIRT6 encodes a NAD+-dependent deacetylase and master regulator of DNA repair, telomere maintenance, and metabolic homeostasis, and its knockout in mice produces early atherosclerosis. This vascular specificity mirrors the molecular finding that arteries, and only arteries, showed exclusive decline in peroxisome pathway activity during ASA periods (**Figure 5B**), a vulnerability that depends on the same NAD+-dependent metabolic processes that SIRT6 regulates. This represents the first evidence linking genetic variation in a known longevity regulator to organ-specific structural aging, demonstrating that aging pathways have tissue-selective rather than uniform whole-body effects.

## 4. Discussion

Our results establish that the physical structure of human tissues deteriorates through organ-specific trajectory, with many cases of discrete and non-linear phases, aligned with their known functional decline, rather than gradual decline. This pattern is invisible to molecular aging clocks trained on chronological age. Three distinct temporal programs, early vascular aging in the 30s, late reproductive aging around menopause, and biphasic aging in digestive and male reproductive tissues, suggest that different organ systems follow fundamentally different aging schedules. This is consistent with recent proteomic evidence of non-linear molecular transitions [7, 8], but adds a critical layer: because tissue structure directly underlies organ function, these structural trajectories may more directly map onto the functional decline that individauls experience than molecular markers alone. The ovary illustrates this: PathStAR captures both fertility decline and menopause from histology images, while matched transcriptomic and methylation profiles from the same samples do not. Using this framework, we provide a tissue-resolved map of what changes when during structural aging: which pathways are active during the first acceleration (hormone signaling) versus the second (damage response), and which disruptions are organ-specific (peroxisomal metabolism in arteries, spermatogenesis in testis). This map identifies both the molecular targets and their timing across organs, a prerequisite for designing tissue-appropriate geroprotective strategies.

We also find that structural aging is coordinated across organs within individuals, especially within expected organ systems (vascular, digestive). Importantly, this revealed an unexpected axis linking digestive and male reproductive tissues. Convergent molecular enrichment of hormone pathways in gastrointestinal tissues suggests shared hormonal regulation of both organ systems, providing a therapeutic opportunity.

Several methodological limitations warrant consideration in interpreting these biological insights spanning feature extraction, temporal resolution, data heterogeneity, and interpretability. Our approach generates slide-level embeddings through mean pooling across all patches, representing a relatively blunt aggregation method that could be considerably improved through more sophisticated attention-based or hierarchical pooling strategies. The temporal resolution of our analysis is constrained by available data points in each age group, necessitating a sliding window approach with 10-year increments that may obscure finer structural transitions occurring at narrower time intervals. The GTEx dataset’s inherent heterogeneity, comprising tissues from donors with diverse clinical backgrounds including ventilator cases and varying postmortem intervals, introduces potential confounding variables that could influence structural assessments beyond chronological aging effects. Finally, our reliance on deep-learning morphological features leaves multiple structural patterns during aging with limited biological interpretability and developing more interpretable feature extraction methods that directly map to known histological patterns would significantly advance the framework’s clinical utility and biological understanding. Not all age-associated structural changes necessarily imply functional decline. Some alterations may reflect adaptive remodeling or neutral architectural variation.

In conclusion, our structural aging framework provides a new window into aging biology. As the physical structure of tissue is the basis of its function, structural aging offers an interpretable connection to functional decline during aging than molecular approaches alone. We believe this will lay the basis for assessing anti-aging intervention efficacy by quantifying whether treatments preserve tissue architecture and connecting tissue structural patterns to blood-based biomarkers for non-invasive aging monitoring. Additionally, it may enable to assess the structural effects of different drugs currently in use on different tissues, and the association of drug toxicities and adverse effects with rapid structural alterations in different tissues. In closing, by establishing structural aging as a quantifiable and predictable process, PathStAR opens a new, structural dimension for aging research.

## 5. Methods

### 5.1 Data Acquisition

Whole-slide images were obtained from the GTEx public portal via v10 release (https://www.GTExportal.org/home/). All slides were digitized at 20X magnification (∼0.5 microns per pixel) using an Aperio scanner (SVS format). Protected data, including participant demographic information (chronological age), lifestyle factors, clinical information, and gene expression data, were accessed through dbGAP (Request accession ID: 39103) (20).

### 5.2 Tissue Selection and Filtering

Although GTEx includes samples from 54 tissue types collected from nearly 1,000 adult post-mortem donors, we restricted our analysis to tissues selected using a systematic filtering strategy to ensure adequate statistical power, detectable age-related morphological variation, and robust estimation of structural aging trajectories.

First, we retained only tissues with available H&E-stained whole-slide images and a minimum of 200 samples per tissue, providing sufficient representation across age windows. Second, we required evidence of age-associated morphological change, assessed by correlation between UNI-extracted morphological features and chronological age, as well as age-dependent separation in principal component analyses of the feature space. We built a simple linear regression model and tissues which PC 1-2 showed variance *r*^2^ higher than 0.15 explained by age were picked. We also trained a LDA to separate young vs old and the tissues with AUC higher than .8 were picked.

Together, these criteria ensured that all selected tissues exhibited sufficient sample sizes, clear age-related structural remodeling, and robust aging signals, enabling reliable construction of tissue-specific structural aging trajectories.

### 5.3 PathStAR Pipeline

*Using above filtrated GTEx data, we developed PathStAR pipeline comprising the following three steps:*

#### i. Preprocessing of WSI and Feature Extraction

We implement the vision encoder UNI for WSI feature extraction. UNI gives patch level embeddings that we then aggregate to create whole slide representations. UNI is a large vision transformer (ViT), pre-trained using the state-of-the-art self-supervised learning algorithm DINOv2[X] - a student-teacher knowledge distillation for ViT architecture. UNI was pretrained on the Mass-100K dataset, a dataset containing more than 100 million tissue patches across 20 major tissue types. To preprocess whole slide images for UNI, we implemented Sobel edge detection to first identify regions of the slide containing tissue. Each 20x magnification WSI was then segmented into 512 x 512-pixel patches. Patches were excluded if more than 50% of their pixels exhibited low weighted gradient magnitude, as determined by Sobel. RGB color normalization was applied to remaining tiles using Macenko’s. Following image preprocessing, these patches are passed to UNI to extract 1024-dimension patch level features. We create a whole slide representation by aggregating each 1,024-patch level feature together by taking mean across the patches for each feature, giving us a final output of 1024 vectors. Considering our goal is to capture the overall structure of the tissue, we opted for mean-pooling. Further, a recent study [46] showed that mean pooling performs comparably to more complex aggregation strategies for slide-level representations of whole-slide images.

#### ii. Quantifying Structural Aging Rate

To identify age-related Structural Aging Rate in each tissue, we developed a sliding window analysis framework that quantifies effect sizes while stratifying by biological sex. For each tissue type with ≥200 samples, we implemented a comparative analysis between age-adjacent windows. For a given window size *w* (primarily using *w*=10 years) and target age *t*, we compared the 1024 slide level features from UNI, in the lower window [*t*-*w*, *t*-1] with those in the upper window [*t*, *t*+*w*-1]. This analysis was restricted to valid age ranges (*t* ∈ [min_age+*w*, max_age-*w*]) to ensure complete windows.

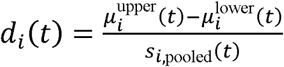

For each feature dimension *i* (1≤*i*≤1024), we calculated total change in the whole-slide morphology vector and the overall effect size as:

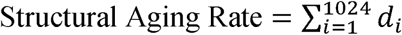

Statistical significance was assessed using Welch’s t-test with significance threshold α=0.05. We defined significant age transitions as those where ≥5% of features showed statistically significant changes after testing. For visualization and analysis, we computed the mean absolute effect size across all features for each target age.

##### Filtering Structural Aging Rate Data points with low statistical Confidence

We only considered age points with sufficient samples (n≥10) in both lower and upper windows, and only these were included in the analysis. We enforced a statistical robustness criterion, retaining only tissues in which at least 80% of estimated structural aging rates were statistically significant, defined as cases where more than 5% of morphological features (UNI derived) exhibited significant effect sizes during rate estimation (Methods 5.3). Later only the points where more than 5% UNI features showed a significant structural aging rate (p <0.05) were included in the smoothening process.

All analyses were conducted together and then separately for male and female samples to identify sex-specific structural aging patterns. This approach enabled identification of critical age thresholds where substantial morphological structural aging occurs across different tissue types and biological sexes.

##### Generating Structural Aging Trajectory

Smoothing Structural Aging rates across all age points, we developed Structural Aging trajectory in each tissue type by smoothening the datapoints using Spline and Gaussian Smoothening. For spline smoothing, we used Univariate Spline with a smoothing parameter calculated as s = (window size / n) × 0.5 × n, where window size = 10. The Gaussian process utilized RBF kernels with length scales proportional to window size for comparable smoothing behavior. Then one with best fit was chosen for the tissue type.

#### iii. Individual Level Tissue Score

A sliding window approach was implemented to quantify morphological deviations within tissue-specific contexts. For each target sample *i* with age ai ≥ 30 years, a reference group Ri was defined using the morphological Structural Aging R[*ai* - 10, *ai* - 1][*ai* - 10, *ai* - 1]

An effect size metric was defined to quantify the deviation in morphological Structural Aging Rate magnitude j ￼, the effect size was computed as the absolute mean difference:

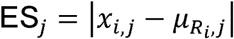

Where *x_i_,_j_* represents the *j*-th feature dimension of the target sample *i*, and 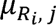 is the mean of the j-th feature across the reference group *R_i_*. This effect size metric captures the absolute magnitude of morphological deviation without normalization by reference group variability, providing a direct measure of the morphological distance between target samples and their younger counterparts. The final effect size for each target sample was computed as the mean across all p feature dimensions:

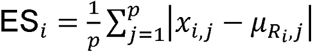

The effect size provides an interpretable measure of morphological changes, with larger values indicating greater deviation from the typical morphology of individuals within the same tissue.

#### iv. Trajectory for Molecular data

To enable direct comparison with structural Aging, we applied the same sliding-window framework to molecular datasets. For tissues with ≥150 samples, we selected the top 20,000 and top 3,000 highly variable genes (or CpGs for methylation) based on variance across samples. Using a 10-year window, we computed mean absolute effect sizes between adjacent age bins, assessed statistical significance with Welch’s t-test (FDR-adjusted), and defined significant transitions as those with ≥5% of points significantly (p <0.05) changing.

To reduce dimensional noise and capture dominant aging structure, we additionally computed trajectories using the top 50 principal components derived from the 20,000-feature set. Across tissues (**Figures R9–R10**), PCA-based trajectories exhibited smoother and more consistent age-dependent patterns, indicating that low-dimensional representations better capture underlying molecular aging dynamics compared to raw feature-level analyses.

To reduce noise and capture dominant aging structure, we additionally computed trajectories using the top 50 principal components derived from the 20k feature set. Across tissues (**Figures R9–R10**), PCA-based trajectories exhibited smoother, more consistent age-dependent patterns, indicating that low-dimensional representations better capture the underlying molecular aging trajectory compared to raw feature-level analyses.

To further reduce high-frequency noise and emphasize dominant age-associated transitions, trajectories were smoothed using a uniform spline-based approach. This approach prioritizes detection of broad inflection periods over fine-grained year-to-year fluctuations. While localized peaks may reflect genuine biological variation, the current cohort sizes limit statistical power to robustly resolve such short-interval effects. Larger cohorts will be required to determine whether higher-frequency trajectory features are reproducible.

### 5.4 Identifying pathways up/down regulated during Accelerated Structural Aging periods

We developed a comparison framework to identify the key pathways up/down regulated during major ASA periods. The key to this was utilizing paired transcriptomics and methylation data from the same biopsies available at GTEx Portal. We identified pathways up/downregulated during ASA1 by comparing the expression profile of samples from ASA 1 with the 5 age periods where accelerated Structural Aging graph shows inflection point. Gene expression profiles were filtered to include only protein-coding genes based on established gene symbols. Expression values for each gene were extracted for both sample groups, and genes with missing data in either group were excluded from analysis. Statistical significance was assessed using Welch’s two-sample t-test, with log2 fold changes calculated as log2(mean_group2 + 1e-5) −log2(mean_group1 + 1e-5) to avoid division by zero. All genes from the differential expression analysis were ranked by log2 fold change in descending order and subjected to Gene Set Enrichment Analysis (GSEA) using GSEApy against the [pathway database] with 10,000 permutations. Pathways with false discovery rate (FDR) q-value < 0.1 and absolute normalized enrichment score (|NES|) > 1.5 were considered significantly enriched.

Above analysis was repeated for methylation, also downloaded from GTEx portal, separately for hypermethylated and hypomethylated states._Gene Set Enrichment Analysis (GSEA) was performed using the GSEApy package with pre-ranked gene lists derived from differential expression results. Pathway enrichment was assessed against multiple curated gene set collections from the Molecular Signatures Database (MSigDB), including KEGG pathways, Gene Ontology (GO) terms, and the C3 collection containing transcription factor targets (TFT).

### 5.5 Cluster the ovary feature changes with age to find modules

To identify age-informative feature modules in ovary tissue, we employed a two-stage approach combining mutual information (MI) feature selection with K-means clustering. Mutual information between each feature and continuous chronological age was calculated using k-nearest neighbor density estimation, which directly handles continuous age values without discretization. Features with higher MI scores indicate stronger association with age-related changes. The top 200 features with highest MI scores were selected and subsequently clustered using K-means clustering after z-score normalization to ensure equal weighting across features with different expression ranges. Clustering was performed across multiple K values (4, 5, 6, 9, 12, 15, and 20) with K-means initialization and fixed random seed (47) for reproducibility. Each resulting cluster represents co-regulated features exhibiting similar trajectories.

### 5.6 Mechanistic Interpretation of Morphological Features using vision language models

To interpret morphological features extracted by UNI, we employed two state-of-the-art vision-language models: PLIP. PLIP leverages contrastive learning on paired pathology images and diagnostic text to learn clinically relevant representations, while Conch, developed by the Mahmood laboratory, specializes in histopathological image understanding through self-supervised learning on large-scale tissue collections.

We established interpretability baselines by computing patch-level semantic similarity scores via dot product operations between patch embeddings and a curated vocabulary of 500 terms, comprising 10 ovary-specific descriptors, 240 general pathological terms, and 250 common English words. Slide-level similarity scores were derived by aggregating patch-level scores using the 95th percentile to capture the most relevant morphological patterns.

Three complementary analyses were performed: (1) validation against ground truth annotations from GTEx pathology reports to assess model accuracy, (2) correlation analysis between similarity scores and chronological age for ovary-specific features including follicular structures, stromal fibrosis, and tissue atrophy, and (3) comparative analysis of pathological features during the menopausal transition period. This multimodal approach enabled quantitative interpretation of morphological changes while maintaining clinical relevance and biological plausibility.

### 5.7 Cross-Tissue Aging Correlation Analysis

To compute cross-tissue co-ordination in Structural Aging during aging, we computed correlations between accelerated Structural Aging or delta Structural Aging scores (denoted as ΔSA) using using Pearson’s correlation coefficients among organs implemented in SciPy. For each tissue pair, individuals with missing data in either tissue were excluded from the correlation calculation to maintain paired observations. Pairwise correlations were computed for ΔSA across all possible tissue combinations, generating both correlation coefficients and associated p-values. A minimum threshold of 10 paired observations was required for correlation calculations to ensure meaningful statistical inference. Statistical significance was assessed at three levels: p < 0.05, p < 0.01, and p < 0.001.

### 5.8 Disease and Lifestyle Association with Accelerated Structural Aging

We tested three categories of clinical features for their association with delta Structural Aging scores (ΔSA): (1) pathology terms extracted from histopathological reports, (2) Hardy Scale death classifications, and (3) binary disease history indicators from phenotypic metadata. For pathology terms, we applied text mining approaches to identify the presence or absence of specific pathological conditions within clinical notes and structured pathology categories for each sample.

Statistical associations were evaluated using multiple linear regression to control for ischemic time as a potential confounding variable. The regression model was specified as: ΔSA ∼ clinical feature + ischemic time + intercept, where clinical feature represents the binary indicator (presence/absence) of each pathological condition, Hardy Scale comparison, or disease history. Missing ischemic time values were imputed as zero to maximize sample retention while maintaining analytical rigor. For each clinical feature, we extracted regression coefficients, p-values, and R² values to quantify the magnitude and significance of associations after controlling for ischemic time effects.

Hardy Scale comparisons were performed systematically across six predefined death circumstance pairs (e.g., violent vs. slow death, intermediate vs. slow death), while disease features compared individuals with and without specific medical histories from the top 20 most prevalent conditions. All pathology terms identified through text mining were tested comprehensively rather than limiting to high-frequency terms. To address multiple testing, we applied Bonferroni correction both within tissues and globally across all comparisons. Only associations with adequate sample sizes (≥10 samples per group) were included in the analysis to ensure statistical power for regression modeling.

### 5.9 Germline Association with Delta SA

We processed the multi-sample GTEx whole-exome sequencing (WES) variant call file (VCF), which contains 979 individuals with variant genotypes and functional annotations. From this file we extracted three layers of information: (i) the core variant descriptors (chromosome, position, reference and alternate alleles, quality, and filter status); (ii) annotation fields from the INFO column, including allele frequency, allele count, quality metrics, and VEP-derived functional annotations such as predicted consequence, impact, gene symbol, and clinical significance; and (iii) individual-level genotypes for each sample, standardized into categorical calls (homozygous reference, heterozygous, homozygous alternate, or missing). This parsing produced structured variant-level tables in which each row corresponded to a single variant, annotated with both functional attributes and per-sample genotypes.

For each tissue, all variants were grouped by gene symbol, and a gene burden score was calculated for every individual by summing the genotypes across all variants mapping to that gene. We also recorded gene-level metadata including the chromosome assignment and the number of variants contributing to each gene’s burden. Gene burden scores were then associated with patient-specific delta SA scores using linear regression of the form:

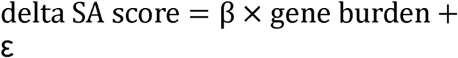

From each model we extracted the slope (β), standard error, p-value, coefficient of determination (R²), and correlation coefficient. The resulting tissue-specific association tables contained, for every gene, the number of contributing variants, number of samples analyzed, and full regression statistics. Multiple-testing correction was applied using the Benjamini–Hochberg false discovery rate procedure, and both complete and significant results were retained.

### 5.10 Molecular Clock

#### 5.10.1 Transcriptomic molecular clock

We trained tissue-specific transcriptomic clocks to predict chronological age from bulk RNA-seq data in GTEx. For each tissue, expression matrices (log-transformed TPMs) were processed after collapsing technical replicates to a single donor-level profile, retaining samples with ≥100 individuals. Within each training fold, genes were pre-filtered by correlation with age, and the top 3,000 most age-associated features (by absolute Pearson correlation) were retained. Features were standardized (z-scored) within the training set, and age prediction models were trained using an elastic net regression framework. Regularization hyperparameters (α across 18 log-spaced values and l1_ratio ∈ {0.1, 0.5, 0.9}) were tuned internally by five-fold cross-validation. To avoid information leakage, feature selection and scaling were repeated independently within each fold, and folds were defined by GroupKFold stratification over donor IDs to ensure that multiple samples from the same individual did not cross training–validation boundaries.

Performance was assessed in each fold using mean absolute error (MAE, in years) and coefficient of determination (R²).

#### 5.10.2 Methylation molecular clock

We trained tissue-specific clocks to predict chronological age from DNA methylation profiles. For each tissue with ≥100 samples, methylation matrices and aligned metadata (age, donor ID) were loaded, and feature selection was performed within each cross-validation fold to avoid leakage. In each fold, CpGs were ranked by their absolute Pearson correlation with age, and the top 2,000 sites were retained. Selected features were standardized using training statistics, and age was predicted using elastic net regression with hyperparameters tuned by internal cross-validation (α across 12 log- spaced values; l1_ratio {0.1, 0.9}). Donor IDs were used as grouping factors in five-fold cross-validation, ensuring samples from the same individual did not cross training–validation boundaries.

Performance was assessed in each fold using mean absolute error (MAE, in years) and coefficient of determination (R²).

## 6. Data Availability Statement

The data used in this study were obtained from the Genotype-Tissue Expression (GTEx) Project (accession: phs000424.v10.p2, Project titled #39103). The H&E histological images and transcriptomic data are publicly available and were accessed through the GTEx Portal v10 release (https://GTExportal.org/home/). The germline variant data, phenotype and metadata, e.g. age, lifestyle factors etc. were extracted after appropriate institutional approval and data access agreement (General Research Use: phs000424.v10.p2.c1). The mean-pooled features generated from GTEx histology images using the UNI foundational model have been deposited on Zenodo(<u>link</u>).

## 7. Code Availability

PathStAR is available on GitHub as a software package for academic use. The complete source code and documentation can be accessed at: https://github.com/Sinha-CompBio-Lab/PathStAR.git

## 8. Conflict Of Interest

The authors declare no conflict of interests.

## 9. Authors Contribution

S.S. conceived and designed the study to quantify morphological changes in tissue using hematoxylin and eosin (H&E) histological images. A.Y. developed the computational framework and implemented the PathStAR pipeline for automated quantification of tissue morphological changes from H&E-stained slides. A.Y. performed validation analyses and computational benchmarking of the PathStAR methodology. K.A. prepared visual presentations of the data and performed some of the analysis. A.Y. drafted the initial manuscript. S.S. provided critical manuscript revisions and supervised the research. K.Y and C.K helped in biological interpretation of structural aging rates and trajectories.

## Supporting information

Supplementary Raw Trajectory

## 10 Acknowledgements

This research was supported by the Sanford Burnham Prebys NCI-designated Cancer Center Start-up Fund for Sinha Lab. We thank Peter Adams for feedback on the manuscript.

## Notes

### Competing Interest Statement

AY. And S.S. have filed a provisional patent related quantifying structural aging rate from H&E images.

### Summary of Updates

This version has been revised to update : 1. New Author has been added 2. The abstract has been revised based on the new results generated 3. The figure 2 and 3 has been changed to include results from new analysis 4. Sec 3, 3.1 to 3.6 has been revised to include the new results as well as provide more clarity 5. Method has been updated

