## Supplementary Figures for "Mapping Structural Aging of Human Tissue reveals tissue-specific trajectories and coordinated deterioration"

### Supplementary Notes and Figures

#### Supplementary Notes

**Notes S1: Tissue specific quality control test**: We tested whether GTEx histology collection is appropriate for studying tissue aging by testing whether a tissue captured canonical aging-related pathology (**Figure S2**). Thus, in GTEx biopsies, based on pathology tissues, with age: Artery showed increase in arthrosis and atherosclerosis, & calcification. Colon and esophagus showed a decrease in mucosal layer thickness. Ovary showed a decrease in ova and follicles and an increase in corpora luteum till menopause age and throughout decrease of atrophy. Prostate showed increased stroma, where testis showed a sharp decrease in spermatogenesis. Uterus shows a decrease in both endometrium and myometrium, and an increase in atrophy, where vagina shows an increase in atrophy and epithelium layer.

**Notes S2: Model Selection and Testing across Multiple Digital Pathology Foundation Models:**

A. Why we chose UNI as our feature extractor? We initially selected UNI as the foundational model for PathStAR due to its large, diverse training corpus that also includes normal tissues (>100,000 H&E slides spanning ~20 tissue types from MGH, BWH, and GTEx), which provides broad and unbiased morphological representations. UNI is built on the DINOv2 framework and is optimized for histopathology-scale feature extraction.
B. Model Dependence: To evaluate model dependence, we repeated the full raw structural trajectory analysis (**Figures R6–R8**) using UNI v2**,** CONCH v1.5**,** and Virchow 2 embeddings. Across tissues, UNI v2 and CONCH produced highly similar trajectory shapes to UNI v1, preserving major inflection points and accelerated structural aging phases. Virchow 2 also reproduced the primary trajectory patterns in most tissues, despite being trained predominantly on cancer-focused cohorts.

These results demonstrate that the non-linear structural aging trajectory identified by PathStAR are not model-specific artifacts but reflect stable morphological signals recoverable across independently trained pathology foundation models.

**Notes S3: Modeling Ischemic time**: Considering ischemic time, total time of tissue processing without oxygen, considerably impacts tissue damage, we reason that its correction is critical during PathStAR modeling. We thus tested whether the natural segregation of old vs. young samples in PCA based on morphology is due to ischemic time. To this end, we modelled age as a function of ischemic time, and then add PC1-2 to the model and test whether this addition explains significantly more variance. This was compared using a LRT test. Specifically, in case of PC1-2 addition, 36/40 tissues, addition of PC1-2 addition adds significantly more variance. Specifically, the largest improvements observed in reproductive tissues (uterus $\Delta R^2 = 0.40$, ovary $\Delta R^2 = 0.34$) and neural tissues (tibial nerve $\Delta R^2 = 0.16$). Four tissues (kidney medulla, cervix ectocervix, bladder, brain cerebellum, thyroid, and breast mammary tissue) showed no significant improvement. For our trajectory analysis, we focused on tissues where PC1-2 adds significantly more variance, aside of three criteria in Notes S4.

**Notes S4: Modules of Morphological Changes for ovary:** We identified age-informative feature modules in ovary tissue using a two-stage approach: first selecting the top 200 features most associated with chronological age via mutual information analysis, then clustering these features using K-means (K= 4-20) to identify co-regulated modules with similar age-related expression trajectories. This analysis revealed three distinct temporal patterns: features that monotonically increase with age, features that monotonically decrease with age, and features that remain relatively stable across the aging process. Higher K values refined these modules by partitioning them based on magnitude of change while preserving the fundamental tri-modal pattern. (**Figure S4**)

**Notes S5: Testing PLIP robustness to capture ovary-pathology**: We evaluated PLIP's robustness to predict ovary-specific pathology using the following framework. We generated PLIP's embedding on 500 diverse terms which consisted of 3 categories: ovary-specific pathology terms, general pathological vocabulary, and common English words unrelated to ovary (**Figure S5, Method 5.6**). The ovary-specific pathology terms are: (Ovum, Ova, Fibrosis, Corpora Albicans, Corpora Albicantia, Corpus Luteum, Calcification, Menopause, and Multi Nucleated Giant Cells). While ovary-specific pathological terms exhibited higher mean similarity scores, the substantial overlap to terms unrelated to ovary suggests limited model specificity. Developing a specific model is out of scope of this study and instead we focused on pathology-terms PLIP can successfully capture, validated by pathologist reports. We identified three ovary-related pathologies that PLIP could reliably detect: Fibrosis, Ovum, and Atrophy (**Figure S6-A**). We next observed that atrophy and fibrosis scores demonstrated a positive correlation with age ($r = 0.38$ & $0.28$, respectively, **Figure S6 B-C**) and ovum scores exhibited a negative correlation with age ($r = -0.341$, **Figure S6-D**). Comparing these three terms across our SSA period, we find out that fibrosis and atrophy show an increase in both SSA periods, whereas ovum shows an overall decline, especially in SSA 2 (**Figure S6 E-G**).

**Notes S6: Choosing tissue for remodeling rate computation:** We applied the following filtering criteria to choose tissue for remodeling trajectory: First, we retained only tissue types with sufficient sample size, requiring a minimum of 200 available samples per tissue. Second, we selected tissues that demonstrated age-related morphological patterns by assessing correlation between UNI-extracted morphological features and chronological age, along with clear age-dependent separation in principal component analysis of the morphological feature space. Third, we applied a statistical robustness filter, retaining only tissues where ≥80% of the remodeling rates were statistically significant. Statistical significance was defined as having >5% of morphological features demonstrate significant effect sizes during remodeling rate calculations. This systematic filtering approach ensured adequate statistical power, demonstrable age-related morphological changes, and robust remodeling signals across all selected tissues for downstream analysis.

**Note S7: Male vs Female:** As shown in **Figure S1, Table S1**, the distribution of male and female samples in our cohort is imbalanced, with males comprising approximately 70% of the dataset. This imbalance, combined with the multiple filtering steps in the PathStAR pipeline, poses challenges for modeling male and female trajectories separately. As a result, the tissue remodeling trajectories presented in the main analysis were jointly modeled across sexes, a limitation of this study. For reference, however, **Figures S9** display the raw remodeling trajectories stratified by sex.

We observed marked differences in structural aging trajectories between males and females. Among the 15 tissues analyzed, five were sex-specific (prostate, testis, uterus, vagina, and ovary), leaving 10 shared tissues for direct comparison. Within the gastrointestinal tract, male samples exhibited a pronounced biphasic aging pattern with two distinct peaks in the colon (transverse and sigmoid), esophagus (muscularis and gastroesophageal junction), and stomach. By contrast, female samples displayed more gradual and moderate changes, without evidence of discrete biphasic peaks.

Distinct vascular patterns were also evident. In the aorta, males showed multiple peaks of structural change, whereas females displayed a sustained elevated rate between ages 30–50 followed by a gradual decline. In the coronary artery, both sexes exhibited an overall decline with age. However, in the tibial artery, males showed a stronger biphasic pattern, with structural aging decreasing until the early fifties and increasing thereafter, while females followed a comparatively attenuated trajectory.

In the tibial nerve, we observed a striking sex divergence: structural aging declined progressively in males, whereas females exhibited the opposite trajectory, with structural change accelerating beyond age 50. Importantly, these findings indicate that male and female tissues not only follow distinct patterns but also operate on different scales of structural aging, underscoring fundamental sex-specific differences in the temporal dynamics of tissue remodeling.

**Note S8: Smoothing strategy for age trajectories:** To model age-related structural trajectories, we evaluated four smoothing approaches: spline regression, Gaussian process regression, LOWESS, and Savitzky–Golay filtering. All methods captured the overall non-linear trends (Figure **S25**); however, spline regression consistently provided the most stable fit across tissues, balancing smoothness with preservation of local transitions and avoiding boundary instability or overfitting. Based on these comparisons, spline smoothing was selected for all downstream analyses.

**Note S9: Correlation between gene expression and structural aging rate**

To test whether molecular trajectories directly mirror structural aging, we correlated age-resolved mean gene expression with the Structural Aging Rate across the lifespan for each tissue. Genes were ranked by Pearson correlation coefficient and subjected to pathway enrichment analysis (Hallmark and KEGG; **Figure S26**). This lifespan-wide correlation approach did not yield biologically coherent or tissue-relevant pathway patterns, likely because opposing molecular programs across distinct structural phases are averaged into a single summary statistic. These results motivated the phase-specific ASA-guided molecular analyses presented in the main text.

### Supplementary Figures

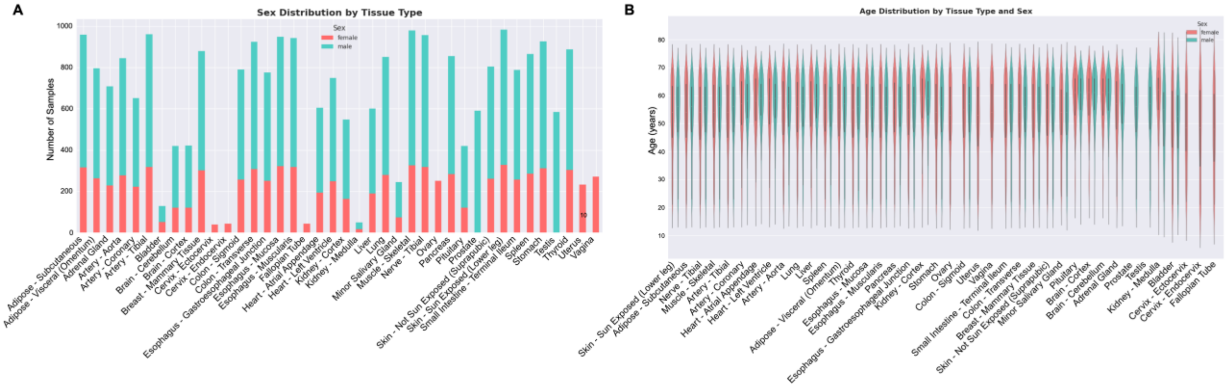


**Figure S1: Sex and Age Visualization of GTEx cohort. (A)** The distribution of data for male vs female. **(B)** Age distribution for both male and female for each tissue. The male samples comprised of total of 66.4% (16808 samples) and female 34.6% (8498). The majority of data comes from older patients with age above 45 years.

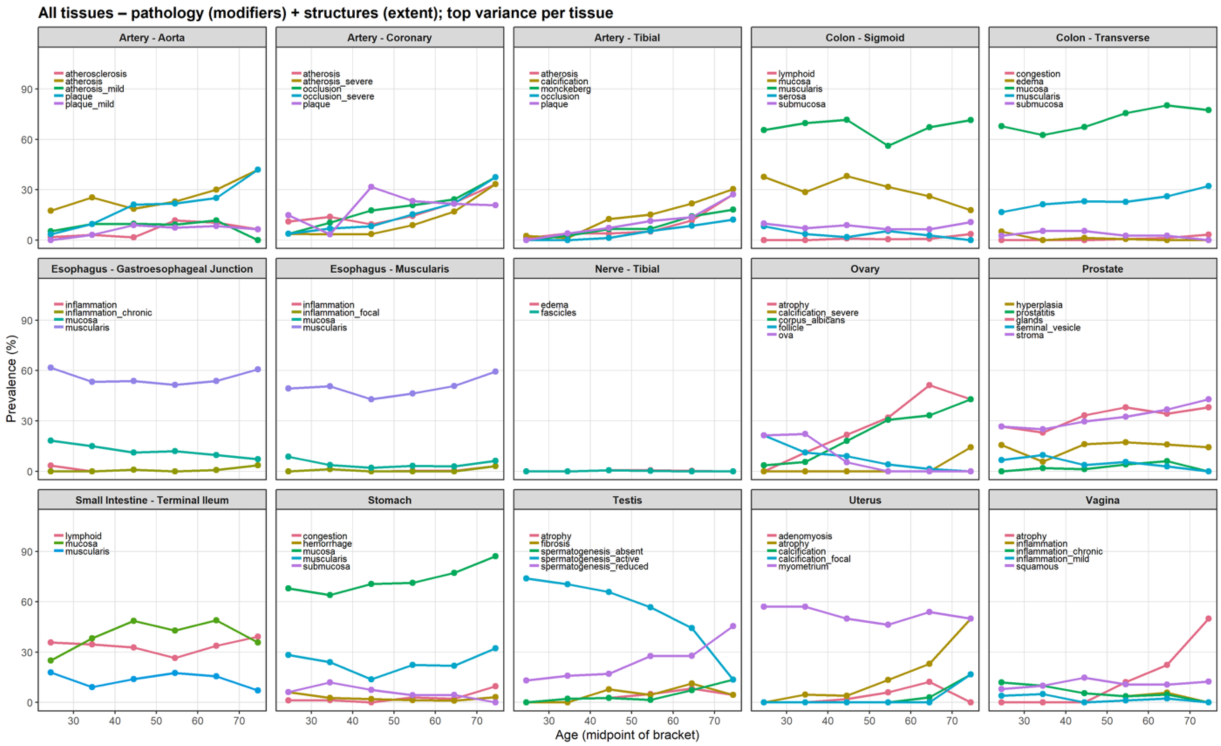


**Figure S2: Age-related changes in the frequency of pathologies across 15 different tissue types.** Each panel shows the prevalence of cases with specific pathological conditions plotted against age groups from 21 years to 70+ years.

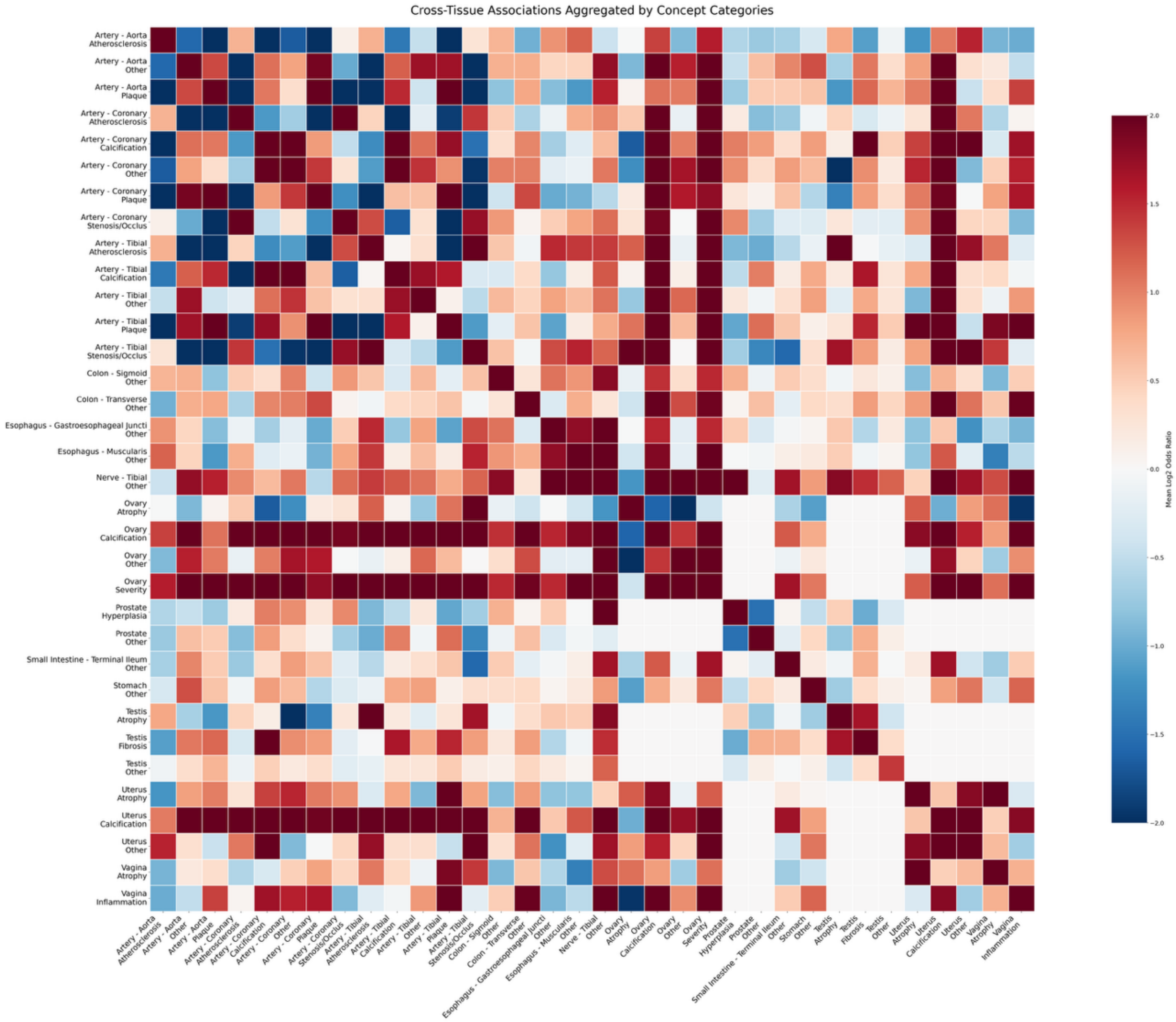


**Figure S2B**: **Cross Tissue Pathology Term Co-occurrence Heatmap.** This heatmap illustrates the strength of association between pairs of pathological findings from the Genotype-Tissue Expression (GTEx) project. To simplify the analysis, numerous specific pathology terms were grouped into the concept categories shown on the x and y-axes. Each cell's color represents the mean Log2 odds ratio, with dark red indicating a strong positive association (i.e., the two conditions tend to occur together) and dark blue indicating a strong negative association.

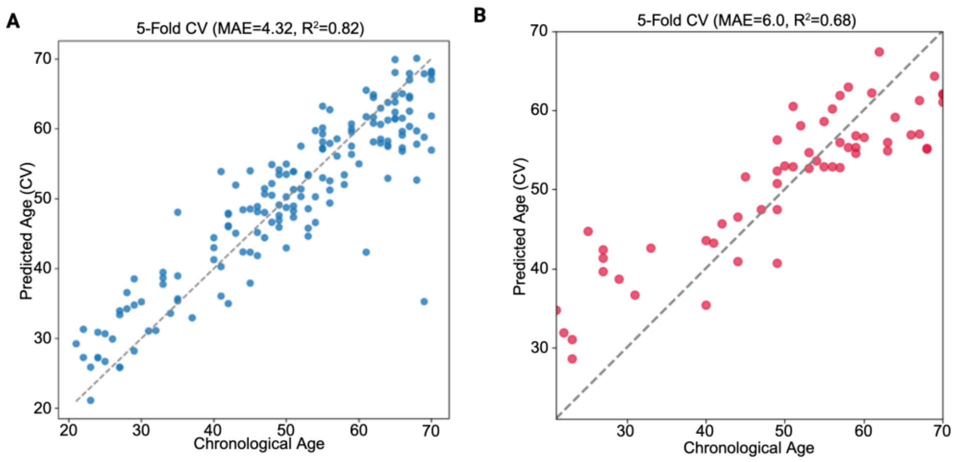


**Figure S3: Methylation and Gene Expression-Based Clocks Ovary:** Performance comparison of age prediction models using 5-fold cross-validation, where x-axis represents chronological age and y-axis shows predicted age. **(A)** Methylation-based model shows high accuracy (MAE = 4.32 years, $R^2$ = 0.82). **(B)** Elastic net gene expression model demonstrates moderate accuracy (MAE = 6.0 years, $R^2$ = 0.68). Dashed diagonal lines indicate perfect prediction.

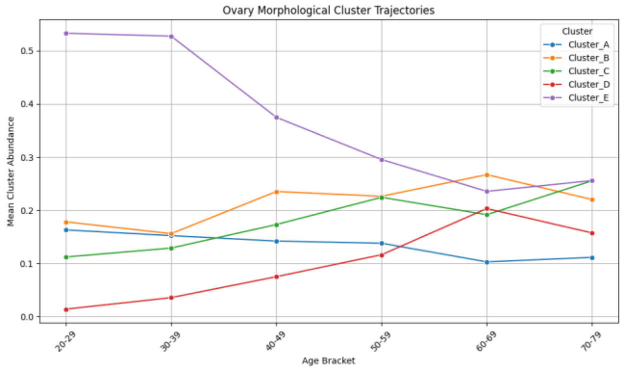


**Figure S4: Ovary morphological organization trend across age**: Patch embeddings were globally clustered with k-means (k=5) to assign fixed morphologic labels A–E. For each slide we computed cluster abundance (fraction of patches per cluster) and averaged abundances within 10-year age brackets. We find that cluster **E** dominates in the 20s (~0.53) but steadily declines to ~0.24–0.26 by late life. Cluster **D** rises from near zero in the 20s to ~0.20 in the 60s, with a slight taper in the 70s. Clusters **B** and **C** increase through mid-life (B peaks ~0.27 at 60–69, then dips; C continues to ~0.26 by 70–79), while **A** gradually decreases (~0.16 → ~0.10–0.11). These abundance shifts are consistent with co-occurrence results showing loss of E self-coherence and growth of D-centered mixing.

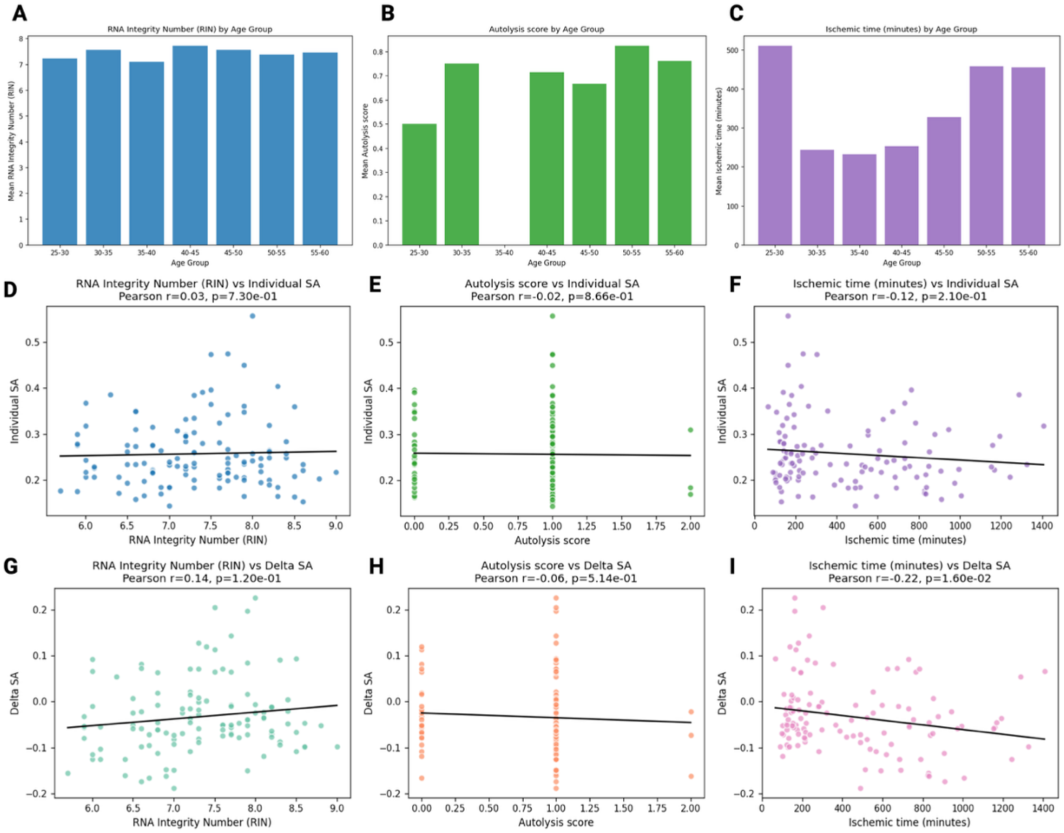


**Figure S5: Quality metrics and covariates across GTEx samples in Ovary. (A–C)** RNA integrity (RIN), autolysis score, and ischemic time stratified by age group. **(D–F)** Associations of these variables with individual structural aging (SA) scores. **(G–I)** Associations with delta SA scores. Only ischemic time showed a modest negative correlation with delta SA

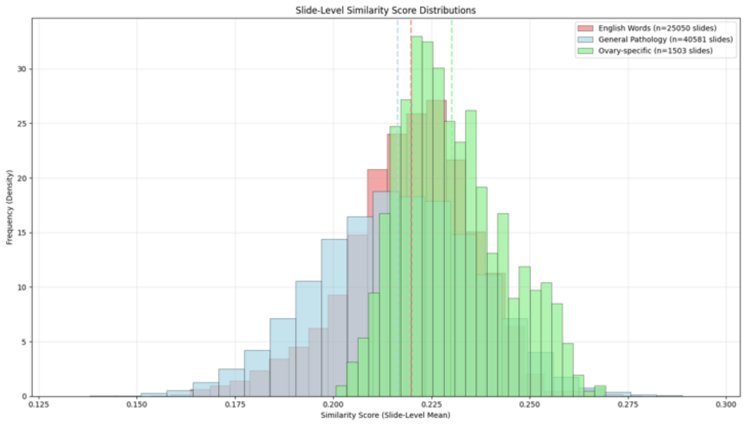


**Figure S6:** Distribution of slide-level (mean across patch) similarity scores between histopathology images and three categories of text terms: ovary-specific terms (green), general pathology terms (blue), and English words (red).

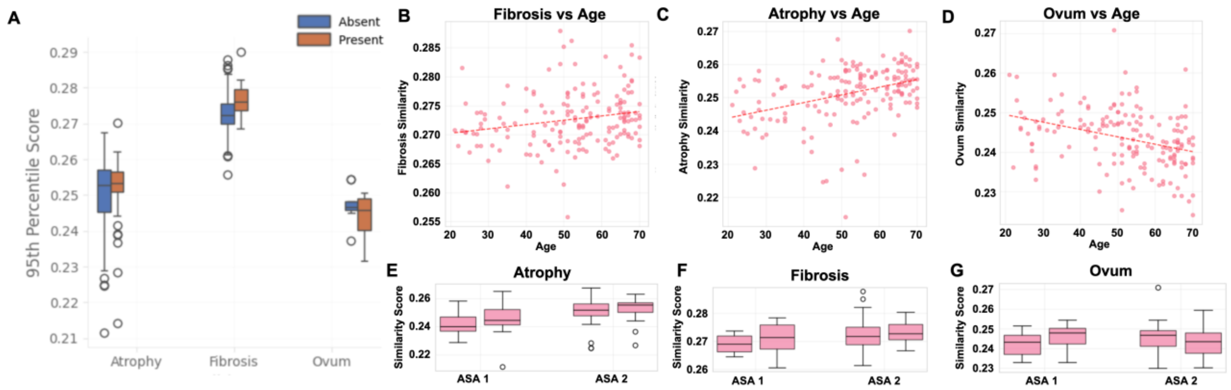


**Figure S7: Interpretation of two ovarian remodeling periods.** ***(A)*** *Box plots comparing 95th percentile similarity scores between pathology-present and pathology-absent cases for fibrosis (n=160 absent, n=7 present), atrophy (n=105 absent, n=62 present), and ovum detection (n=9 each group).* ***(B-D)*** *Scatter plots with regression lines showing age-dependent trends in similarity scores for each pathological feature (atrophy: positive correlation; fibrosis: positive correlation; ovum: negative correlation).* ***(E-G)*** *Box plots stratifying similarity scores by reproductive age groups 25-30 years SSA 1(35-40 years), 45-50 years and SSA 2(55-60 years) for each pathological feature, demonstrating distinct age-related patterns.*

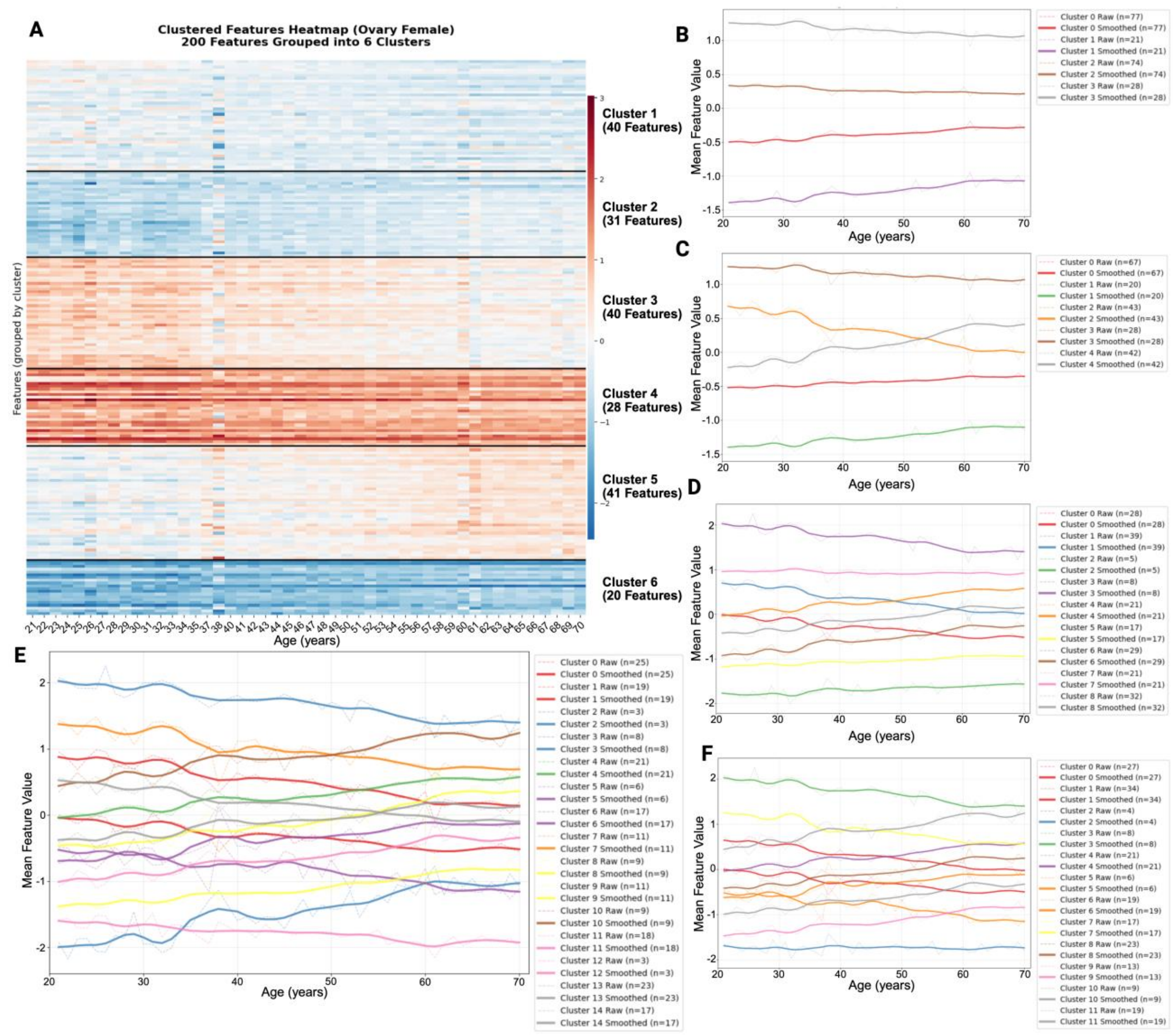


**Figure S8: Identification of Distinct Ovarian Structural Remodeling Modules During Aging.** PathStAR features were ranked by Normalized Mutual Information (NMI) with age, and the top 200 age-associated features were clustered to identify coordinated remodeling patterns. Clustering was performed with k = 4, 5, 6, 9, 12, 15, and 20 to determine optimal feature groupings. **(A)** Heatmap of the top 200 age-associated structural features in ovarian tissue, organized into 6 distinct clusters based on their temporal expression patterns. Features are arranged by cluster membership (indicated by color bars) and show clear age-dependent trajectories from ages 20-70 years. Red indicates higher feature values, blue indicates lower values. **(B-F)** Cluster trajectory plots showing feature group dynamics across different clustering solutions (k = 4, 5, 9, 12, and 15). Each line represents the mean trajectory of features within a cluster, revealing distinct remodeling modules with varying temporal patterns: early declining features, late-declining features, progressively increasing features and stepwise changes. Higher k values provide increased granularity in identifying coordinated structural changes that correspond to specific phases of ovarian aging, demonstrating that tissue remodeling occurs through orchestrated modules.

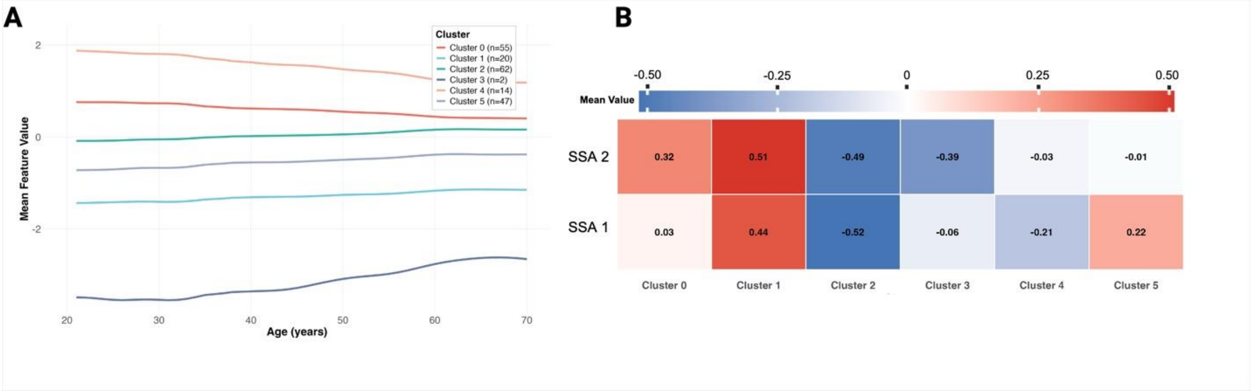


**Figure S9** : **A**) Six different morphological-features modules identified by clustering the top 200 UNI extracted features into 6 clusters, where cluster 0 and 1 showed highest decline with age opposed to cluster 3 which shows increase with age. **(B)** Heatmap showing cluster enrichment in ASA 1 and ASA 2.

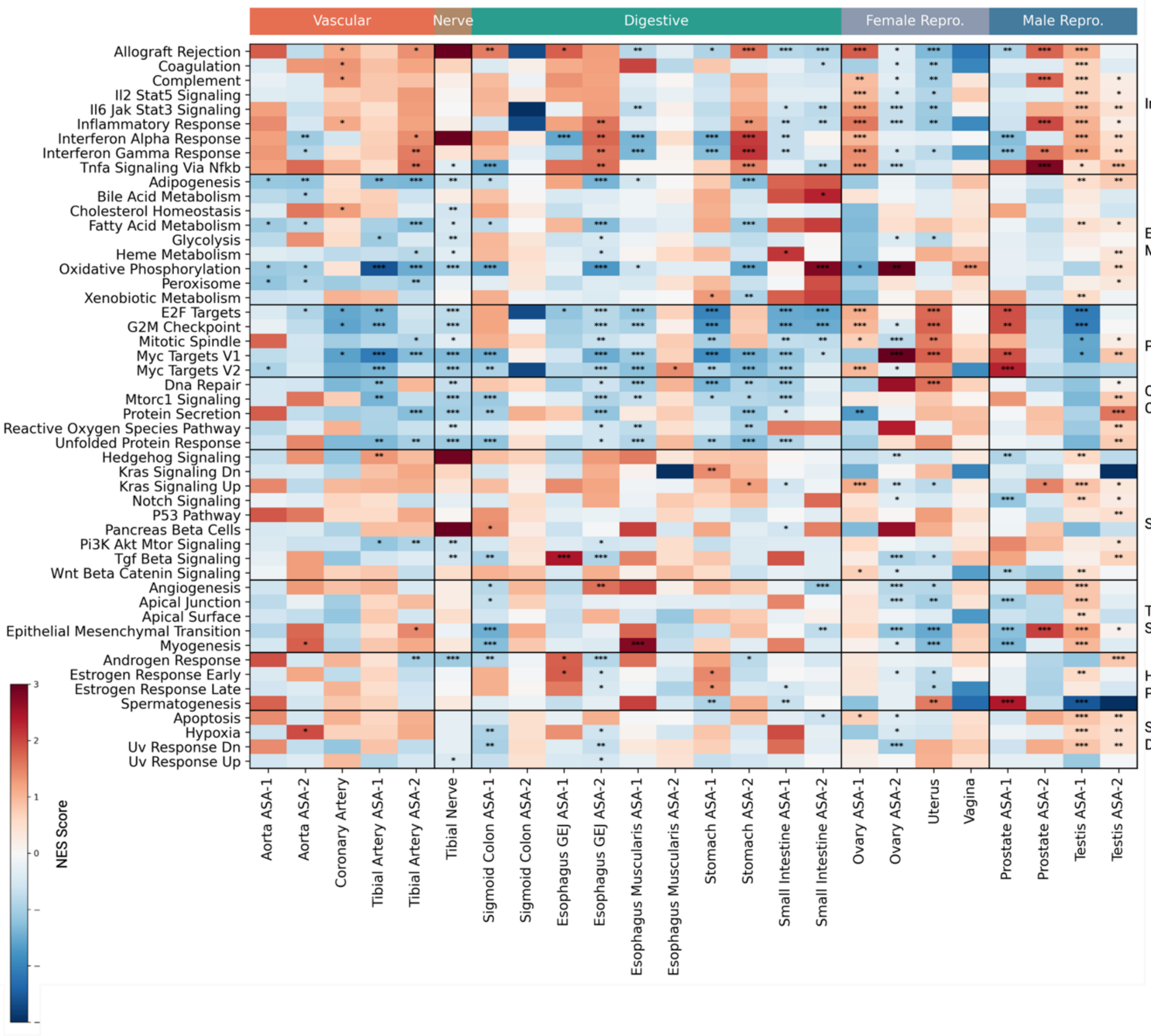


**Figure S10: Hallmark Pathway Enrichment Across Accelerated Structural Aging (ASA) Transitions**. Z-scored Normalized Enrichment Scores (NES) for Hallmark pathways are shown across tissue-specific Accelerated Structural Aging (ASA) transitions. Columns represent tissue–ASA contrasts (ASA-1 and ASA-2 where applicable), grouped by organ system (vascular, nerve, digestive, female reproductive, male reproductive). Rows are organized into eight biological categories: Inflammation, Energy & Metabolism, Proliferation, Quality Control, Signaling, Tissue Structure, Hormone & Reproductive, and Stress & Damage. NES values were z-scored within each tissue–ASA contrast to emphasize relative pathway shifts. Asterisks denote statistically significant enrichment.

Distinct and tissue-specific molecular programs accompany structural aging phases. Vascular tissues show coordinated inflammatory and metabolic remodeling, digestive tissues exhibit shifts in stress-response and quality-control pathways, and reproductive tissues demonstrate pronounced hormone and proliferation-related transitions.

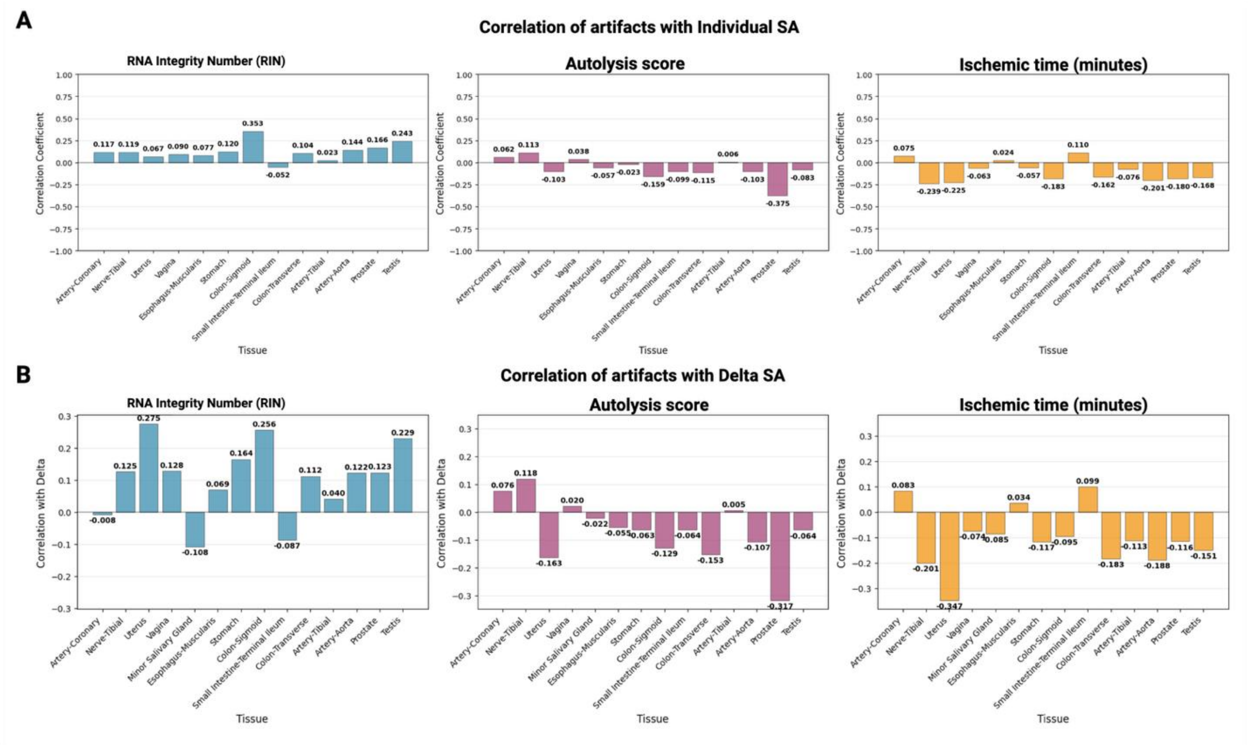


**Figure S11: Correlation of sample-level artifacts with structural age**. **(A)** Correlations between RNA integrity number (RIN), autolysis score, and ischemic time with individual SA across tissues. **(B)** Corresponding correlations with ΔSA. While modest associations are observed in certain tissues (e.g., RIN in colon and testis, autolysis in prostate, ischemic time in nerve and stomach), most correlations are weak or near zero. Together, these results indicate that technical artifacts explain little of the variance in SA or ΔSA, underscoring the robustness of the observed remodeling patterns.

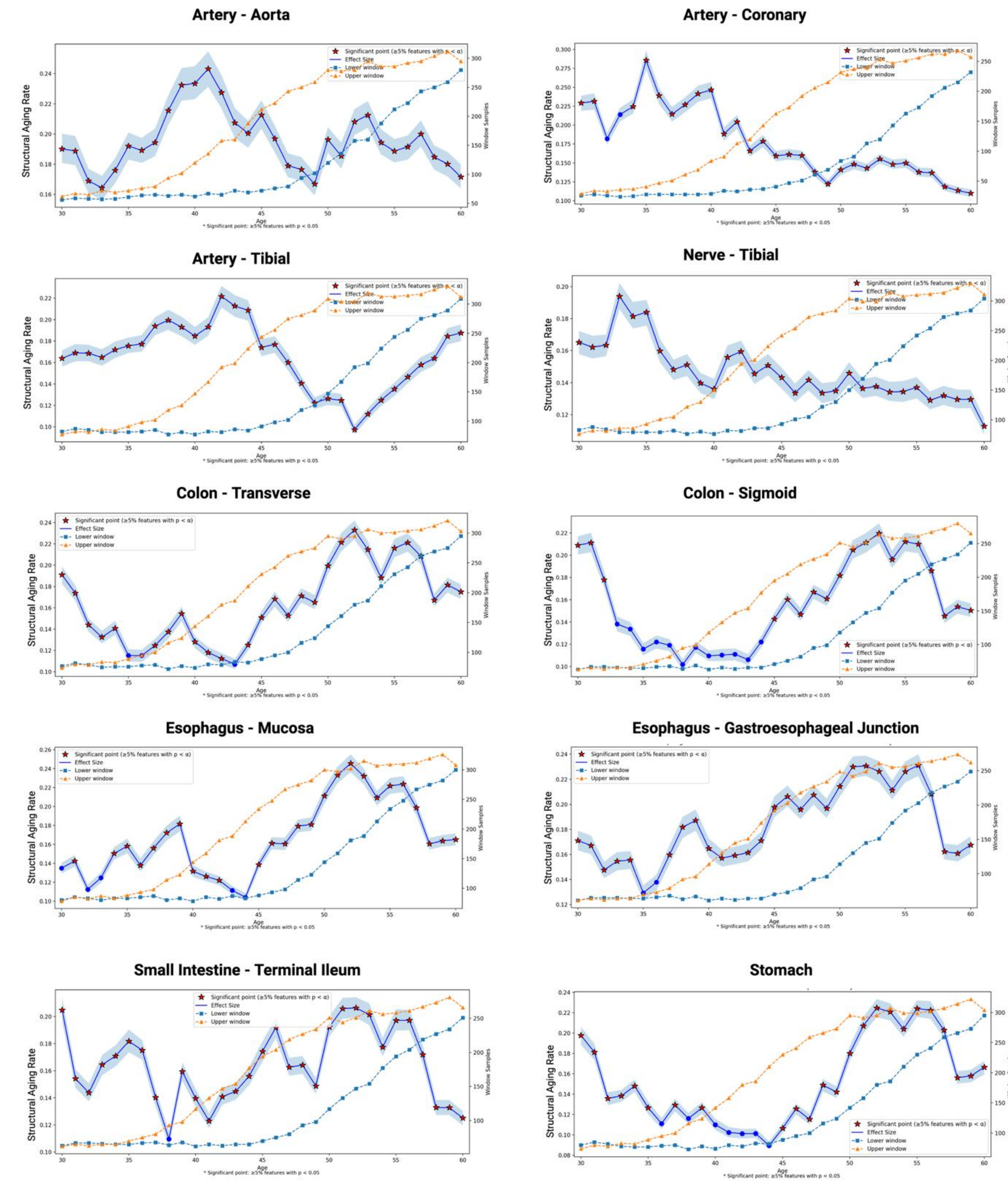
Artery - Aorta
Artery - Coronary
Artery - Tibial
Nerve - Tibial
Colon - Transverse
Colon - Sigmoid
Esophagus - Mucosa
Esophagus - Gastroesophageal Junction
Small Intestine - Terminal Ileum
Stomach
Significant point (≥5% features with p < α)
Effect Size
Lower window
Upper window
Structural Aging Rate
Window Samples
Age
* Significant point: ≥5% features with p < 0.05

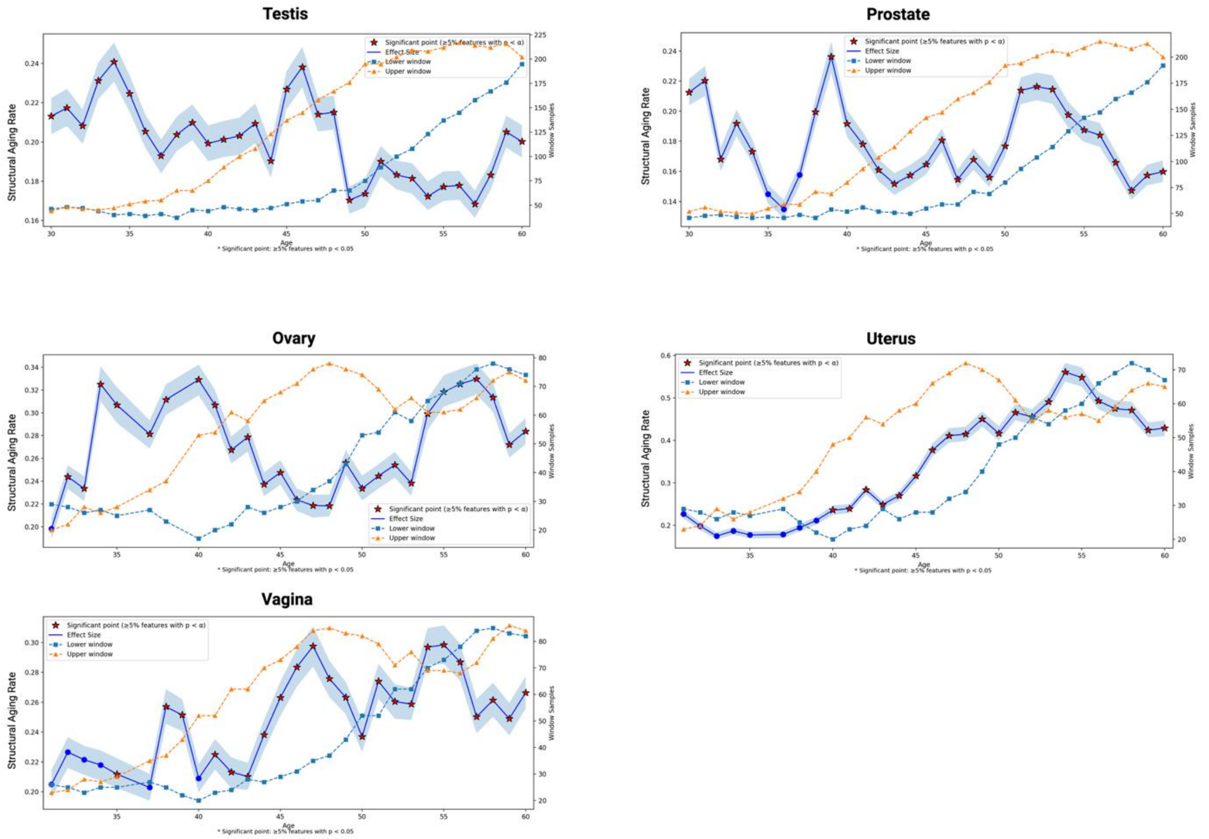


**Figure S12:** Raw Structural Aging Trajectory(blue lines) are shown for the 15 tissues: *Artery (Aorta, Coronary, Tibial), Colon (Sigmoid, Transverse), Esophagus (Gastroesophageal Junction, Muscularis), Nerve (Tibial), Ovary, Prostate, Small Intestine (Terminal Ileum), Stomach, Testis, Uterus,* and *Vagina*, with shaded ribbons indicating 95% bootstrap confidence intervals. Each trajectory represents the magnitude of morphological change between adjacent age windows. Red stars denote age transitions where ≥5% of features exhibit nominal significance ($p < 0.05$). Dashed curves on the secondary axis show the number of samples contributing to the lower (squares) and upper (triangles) age windows at each transition. Together, these trajectories highlight the timing and magnitude of age-related changed across diverse tissue in male and female together.

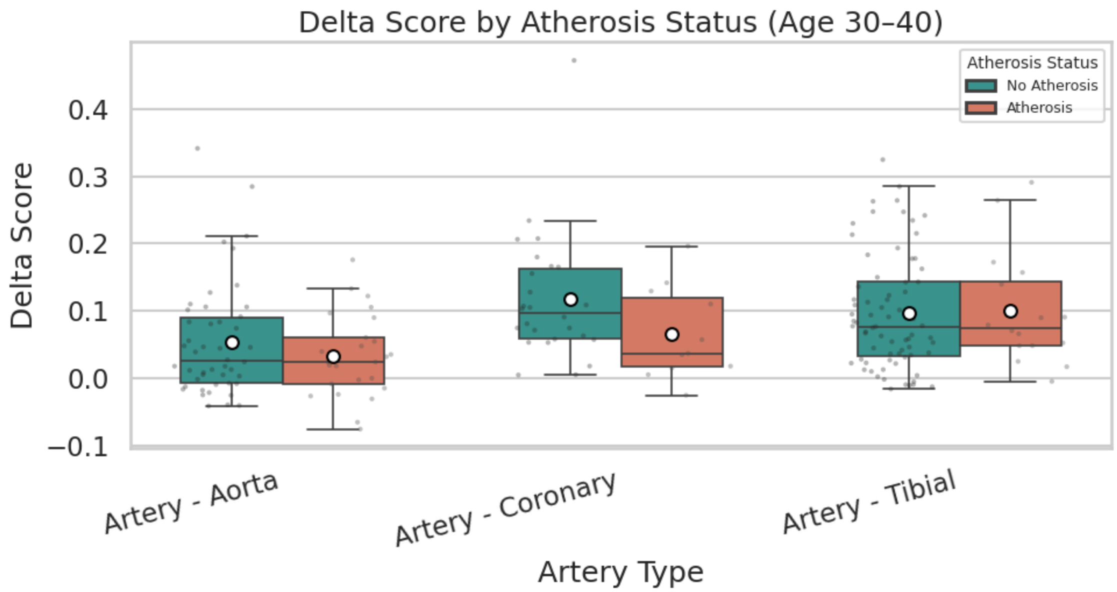


**Figure S13: PathStAR Delta Scores Distinguish Atherosclerosis Presence in Middle-Aged Adults.** Box plots comparing delta SA scores between individuals with atherosclerosis (orange) and without atherosclerosis (teal) across three arterial tissues in the 30-40 age group.

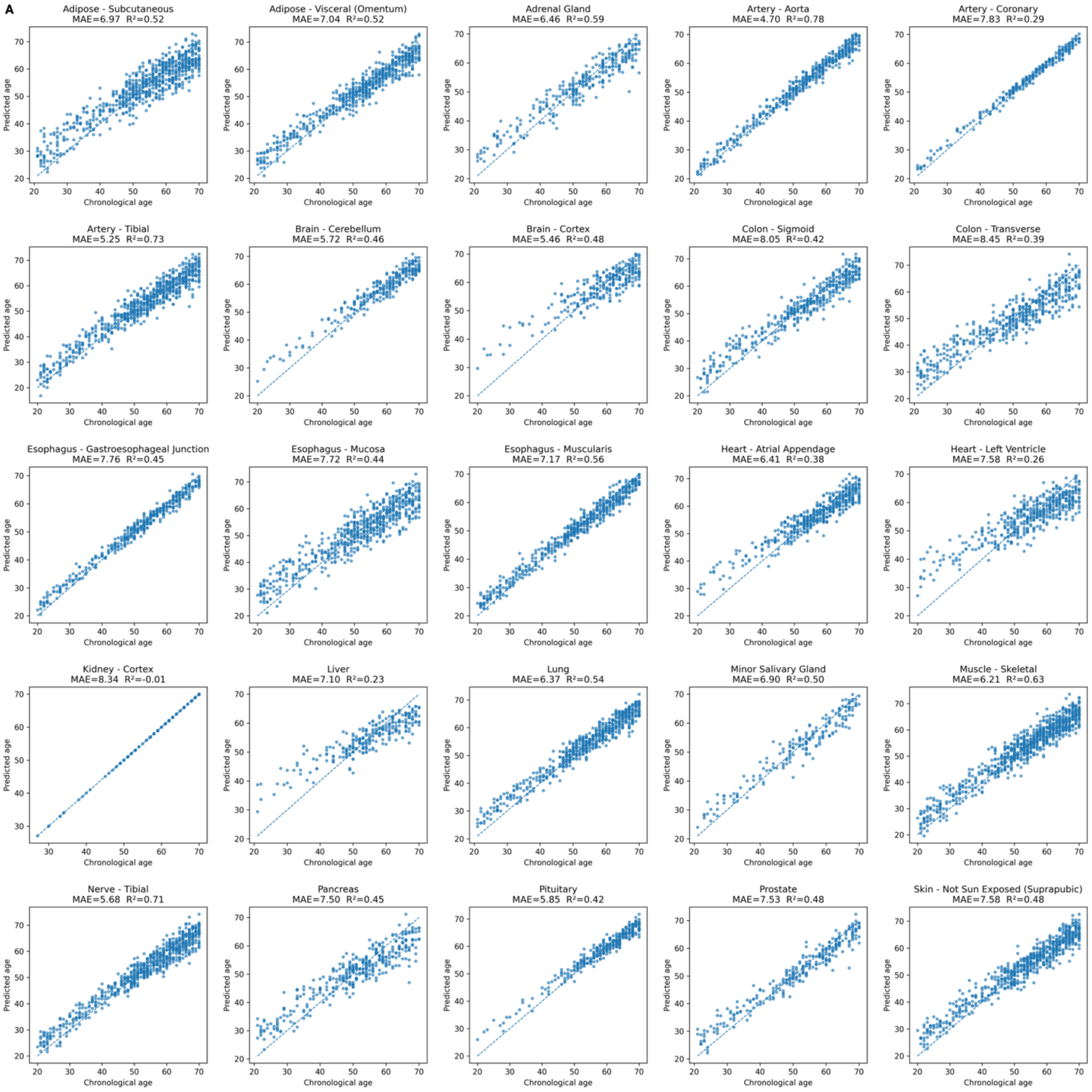
A
Adipose - Subcutaneous
MAE=6.97 R²=0.52
Adipose - Visceral (Omentum)
MAE=7.04 R²=0.52
Adrenal Gland
MAE=6.46 R²=0.59
Artery - Aorta
MAE=4.70 R²=0.78
Artery - Coronary
MAE=7.83 R²=0.29
Artery - Tibial
MAE=5.25 R²=0.73
Brain - Cerebellum
MAE=5.72 R²=0.46
Brain - Cortex
MAE=5.46 R²=0.48
Colon - Sigmoid
MAE=8.05 R²=0.42
Colon - Transverse
MAE=8.45 R²=0.39
Esophagus - Gastroesophageal Junction
MAE=7.76 R²=0.45
Esophagus - Mucosa
MAE=7.72 R²=0.44
Esophagus - Muscularis
MAE=7.17 R²=0.56
Heart - Atrial Appendage
MAE=6.41 R²=0.38
Heart - Left Ventricle
MAE=7.58 R²=0.26
Kidney - Cortex
MAE=8.34 R²=-0.01
Liver
MAE=7.10 R²=0.23
Lung
MAE=6.37 R²=0.54
Minor Salivary Gland
MAE=6.90 R²=0.50
Muscle - Skeletal
MAE=6.21 R²=0.63
Nerve - Tibial
MAE=5.68 R²=0.71
Pancreas
MAE=7.50 R²=0.45
Pituitary
MAE=5.85 R²=0.42
Prostate
MAE=7.53 R²=0.48
Skin - Not Sun Exposed (Suprapubic)
MAE=7.58 R²=0.48
Predicted age
Chronological age

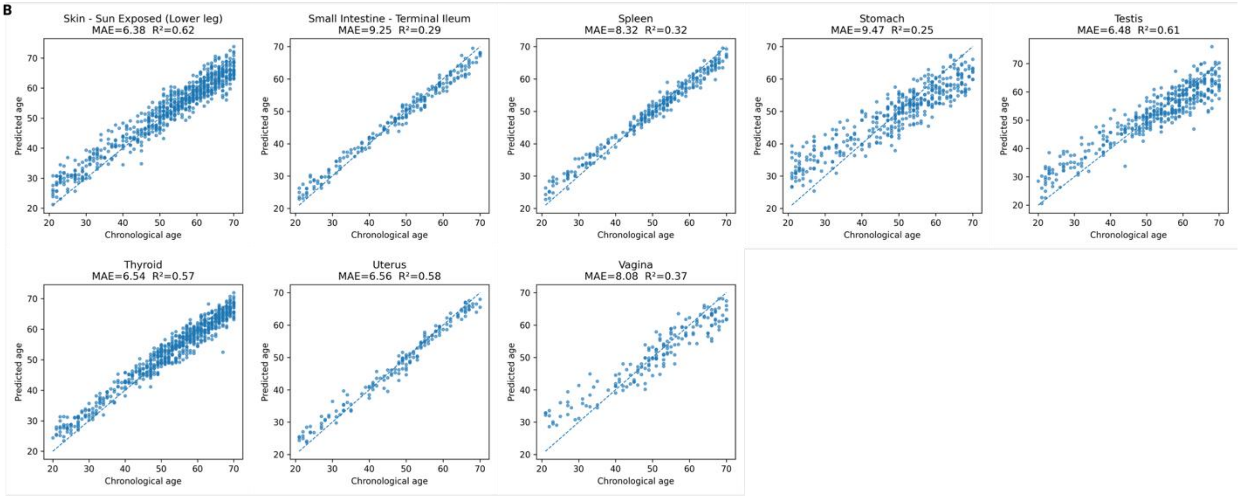


**Figure S14: Transcriptomic Molecular Clocks. (A-B)** Transcription-based molecular clocks trained separately for each tissue type using elastic net regression with 5-fold cross-validation. The clocks demonstrate strong linear correlations with chronological age across tissues (MAE range: 0.57-8.34 years), confirming their ability to capture age-associated molecular changes. However, these linear models fail to detect the non-linear aging trajectories and tissue-specific inflection points revealed by PathStAR's structural approach.

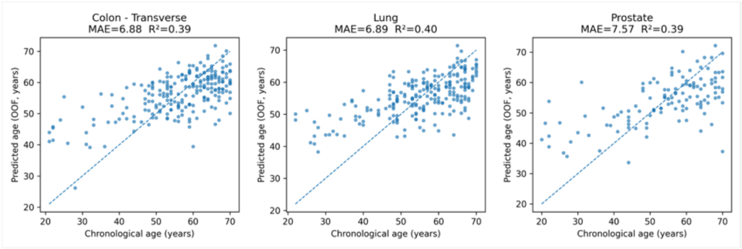


**Figure S15: Methylation Molecular Clocks:** Methylation-based molecular clocks trained for three tissue types with sufficient sample availability, using identical elastic net methodology as transcriptomic clocks. The clocks demonstrate strong linear correlations with chronological age across tissues (MAE range:1.57-8.88), confirming their ability to capture age-associated molecular changes.

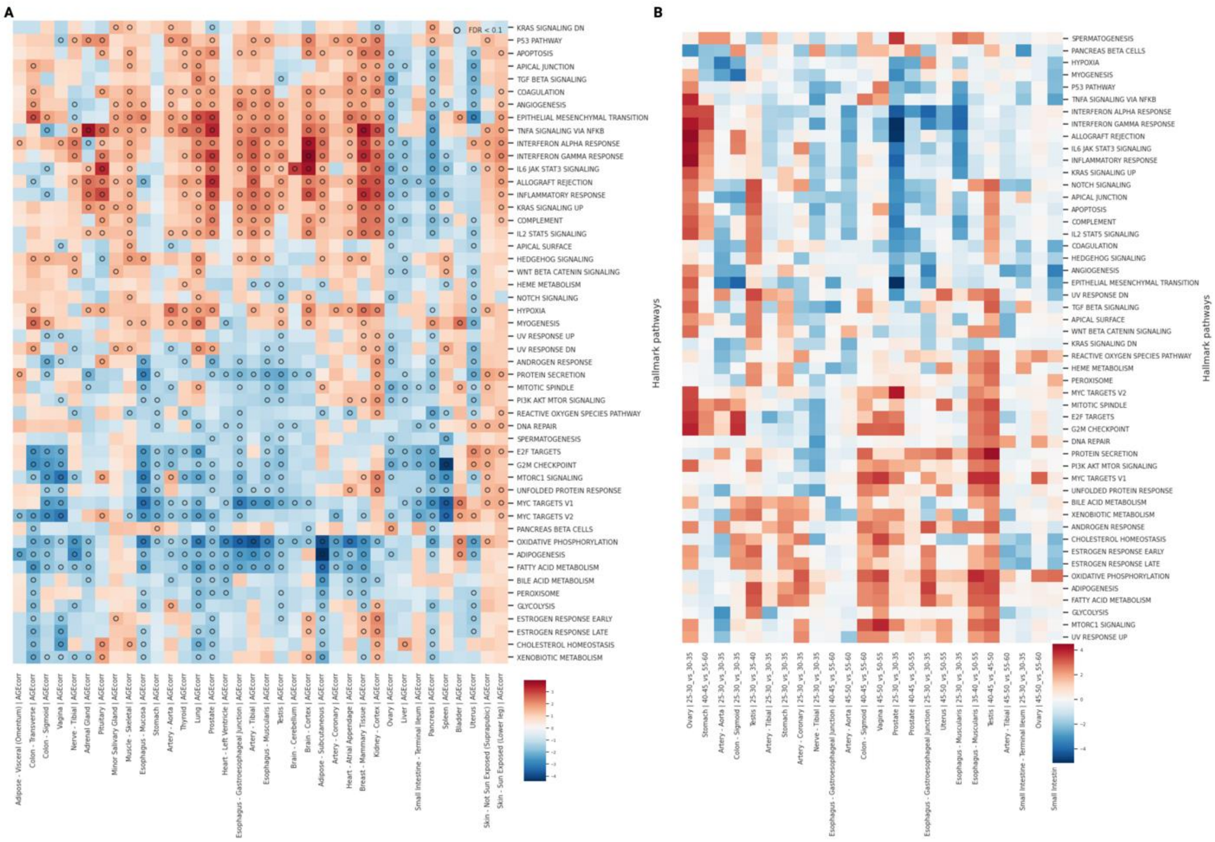


**Figure S16: Age-Related Hallmark Pathway Analysis Reveals Tissue-Specific Aging Signatures. (A)** Gene Set Enrichment Analysis (GSEA) using Hallmark pathways performed on genes ranked by correlation with chronological age across tissues. Normalized Enrichment Scores (NES) reveal distinct tissue-specific patterns of pathway activation (red) and suppression (blue) during aging. Notable patterns include widespread upregulation of inflammatory pathways (TNF-α signaling, inflammatory response, IL6-JAK-STAT3 signaling) across multiple tissues, consistent with the inflammaging hypothesis. Metabolic pathways show tissue-specific divergence, with oxidative phosphorylation predominantly downregulated in metabolically active tissues (muscle, heart, liver) while showing mixed patterns in others. **(B)** Comparative analysis showing the difference between NES scores from the Accelerated structural aging (ASA) period and the overall age-correlation analysis from panel A. This comparison highlights pathways that are specifically dysregulated during periods of accelerated structural aging versus those that change linearly with age.

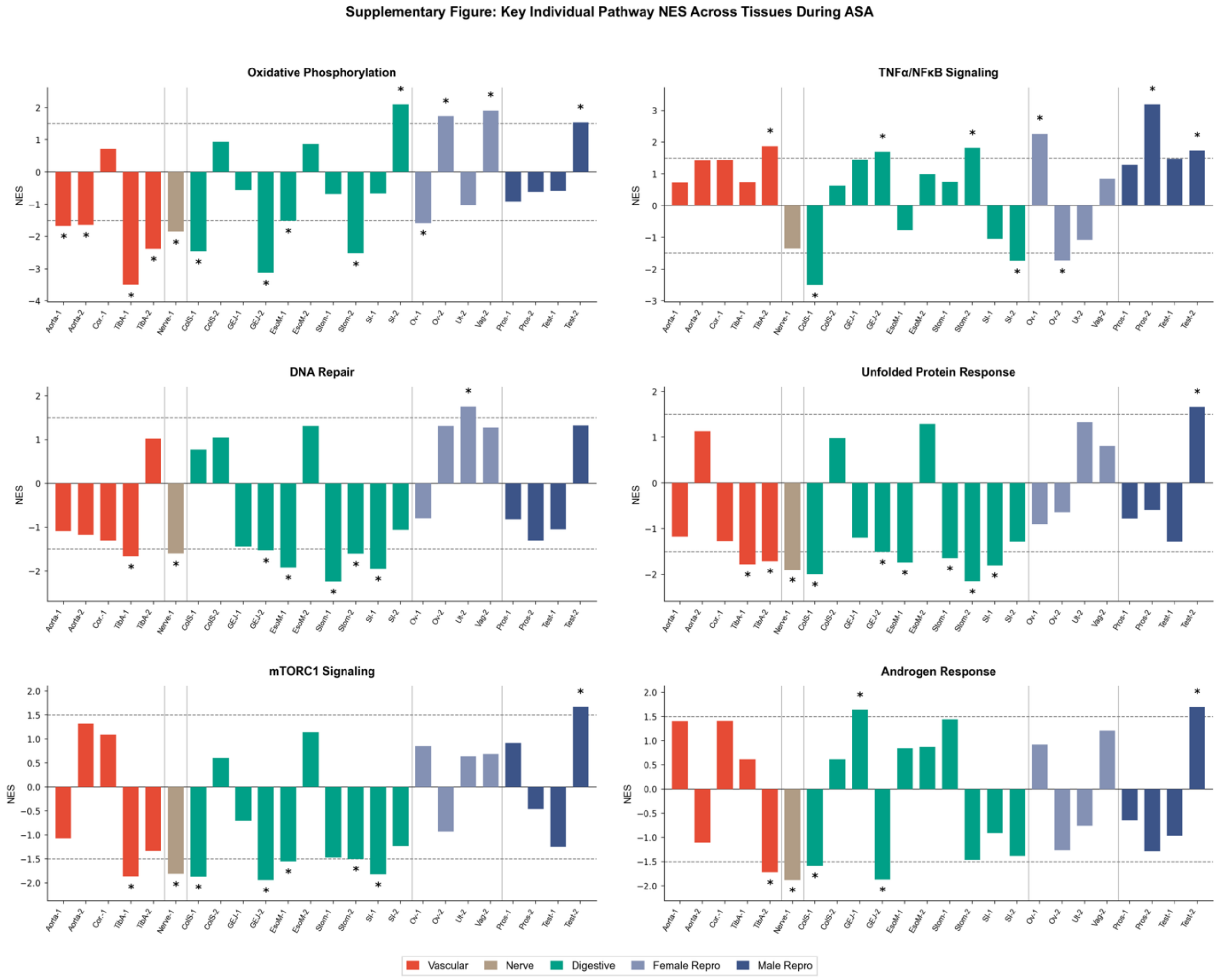


**Figure S17: Key Pathway Changes Across Tissues During Accelerated Structural Aging (ASA).** Normalized Enrichment Scores (NES) for representative Hallmark pathways are shown across tissues during ASA transitions. Pathways include Oxidative Phosphorylation, TNFα/NFκB Signaling, DNA Repair, Unfolded Protein Response, mTORC1 Signaling, and Androgen Response. Bars are grouped by organ system; asterisks indicate significant enrichment.

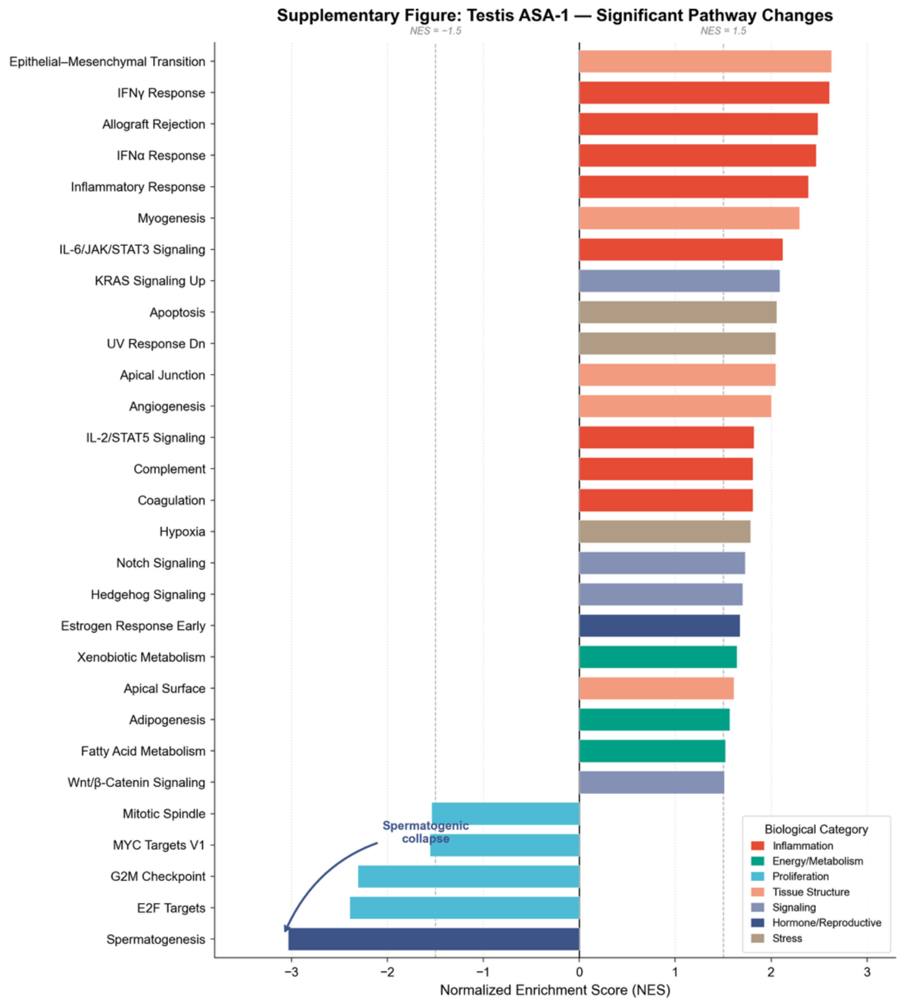


**Figure S18: Testis ASA-1 - Significant Pathway Changes.** Significantly enriched Hallmark pathways during ASA-1 in Testis are ranked by NES. Positive values indicate activation; negative values indicate suppression. ASA-1 is characterized by strong activation of inflammatory and stress pathways alongside marked suppression of spermatogenesis and cell-cycle programs (MYC, E2F, G2M). This pattern indicates a major early remodeling event in male reproductive tissue.

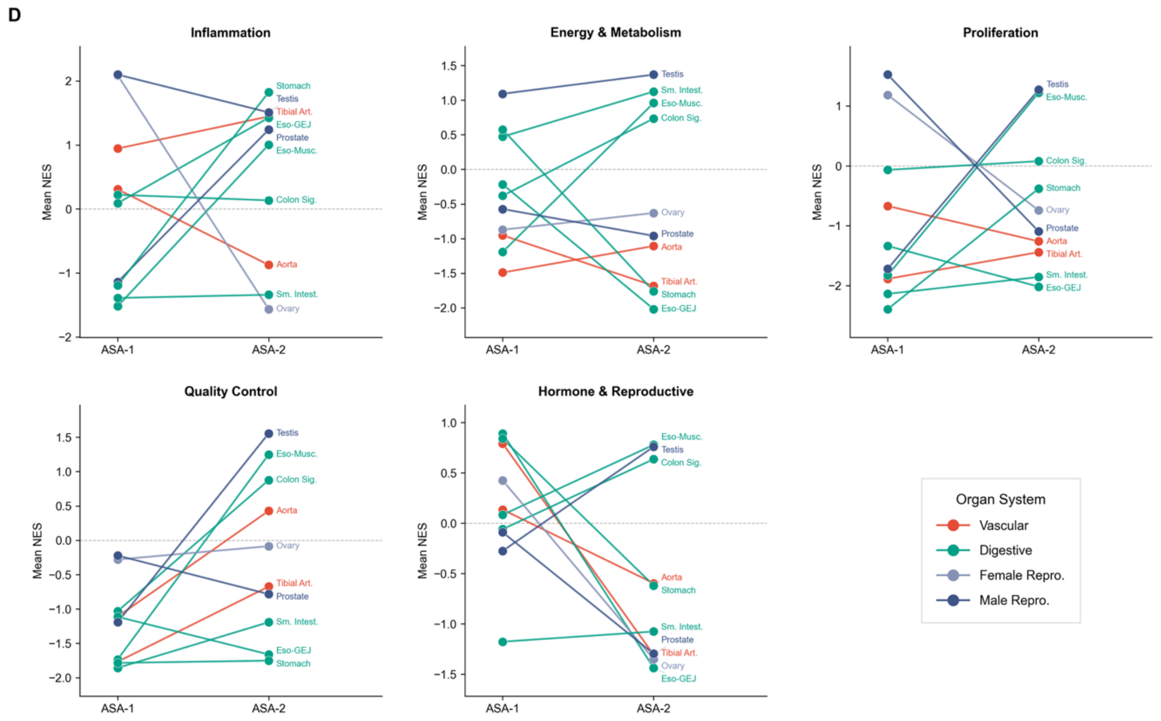


**Figure S19: System-Level Pathway Shifts Between ASA-1 and ASA-2**

Mean NES values for major biological categories (Inflammation, Energy & Metabolism, Proliferation, Quality Control, Hormone & Reproductive) are shown across two accelerated aging phases (ASA-1 and ASA-2). Lines connect the same tissue across phases; colors denote organ systems.

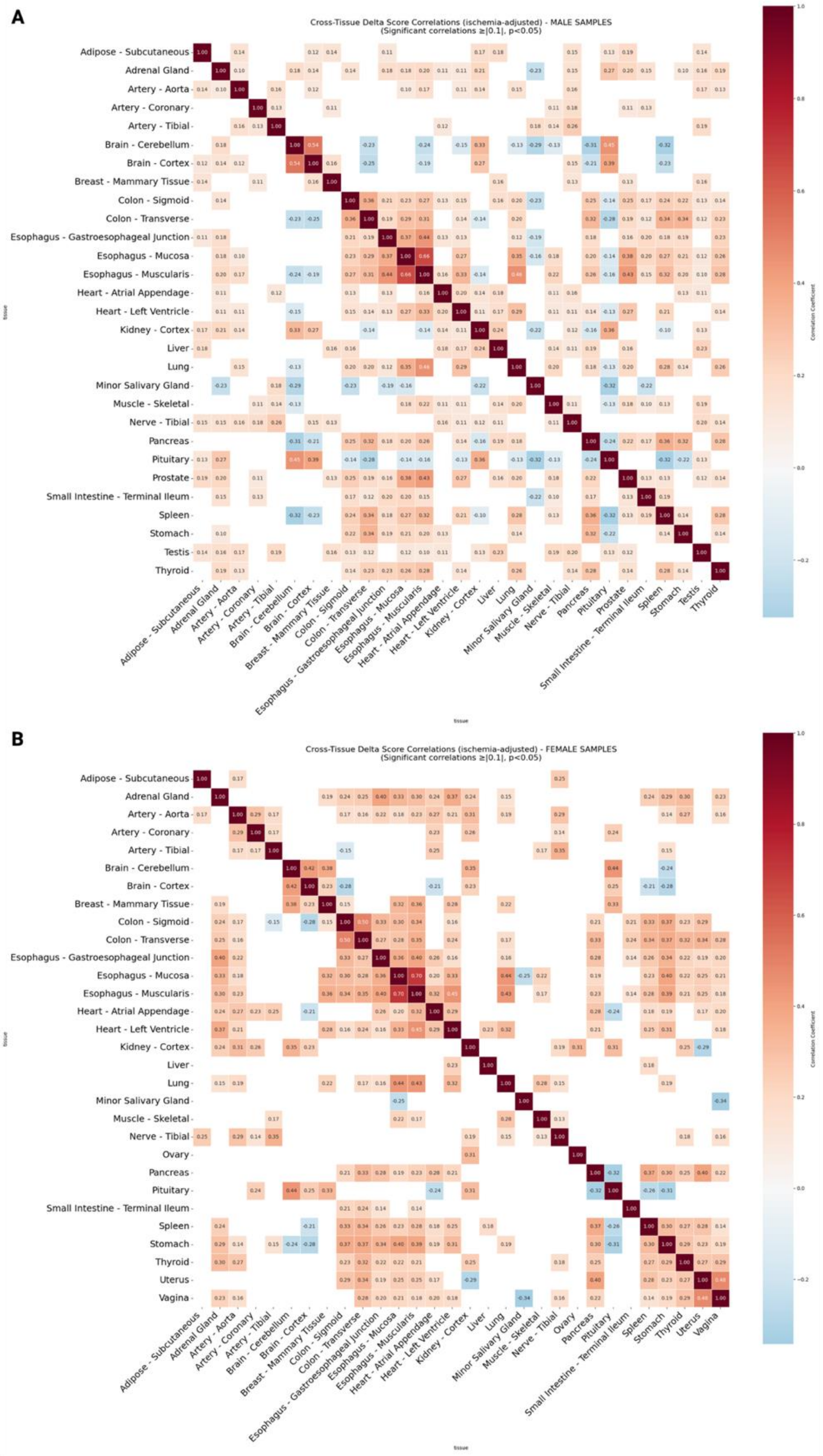


**Figure S20:** Cross organ **(A-B) Heatmaps showing Pearson correlation coefficients of with delta SA (ΔSA) between all tissue pairs across 40 tissues from the same individuals for both male and female.** Color intensity represents correlation strength, with darker red indicating stronger positive correlations and blue indicating negative correlations. Only correlations with ≥10 paired observations are displayed. Statistical significance levels are indicated by asterisks ($p < 0.05$, $p < 0.01$, and $p < 0.001$). The analysis reveals distinct patterns of coordinated aging across tissues, with particularly strong positive correlations observed within canonical organ systems. Notable findings include high correlations within the gastrointestinal system (colon-sigmoid, colon-transverse, esophagus-gastroesophageal junction, esophagus-muscularis, and stomach), reproductive tissues (uterus-vagina correlation of 0.47) in female, and vascular-neural coordination (nerve tibial-artery tibial correlation of 0.28). The stomach demonstrates broad correlations with multiple digestive system organs too. These correlation patterns provide evidence for synchronized deterioration across functionally related tissues during aging. Missing correlations indicate insufficient paired observations (<10 individuals) for reliable statistical inference.

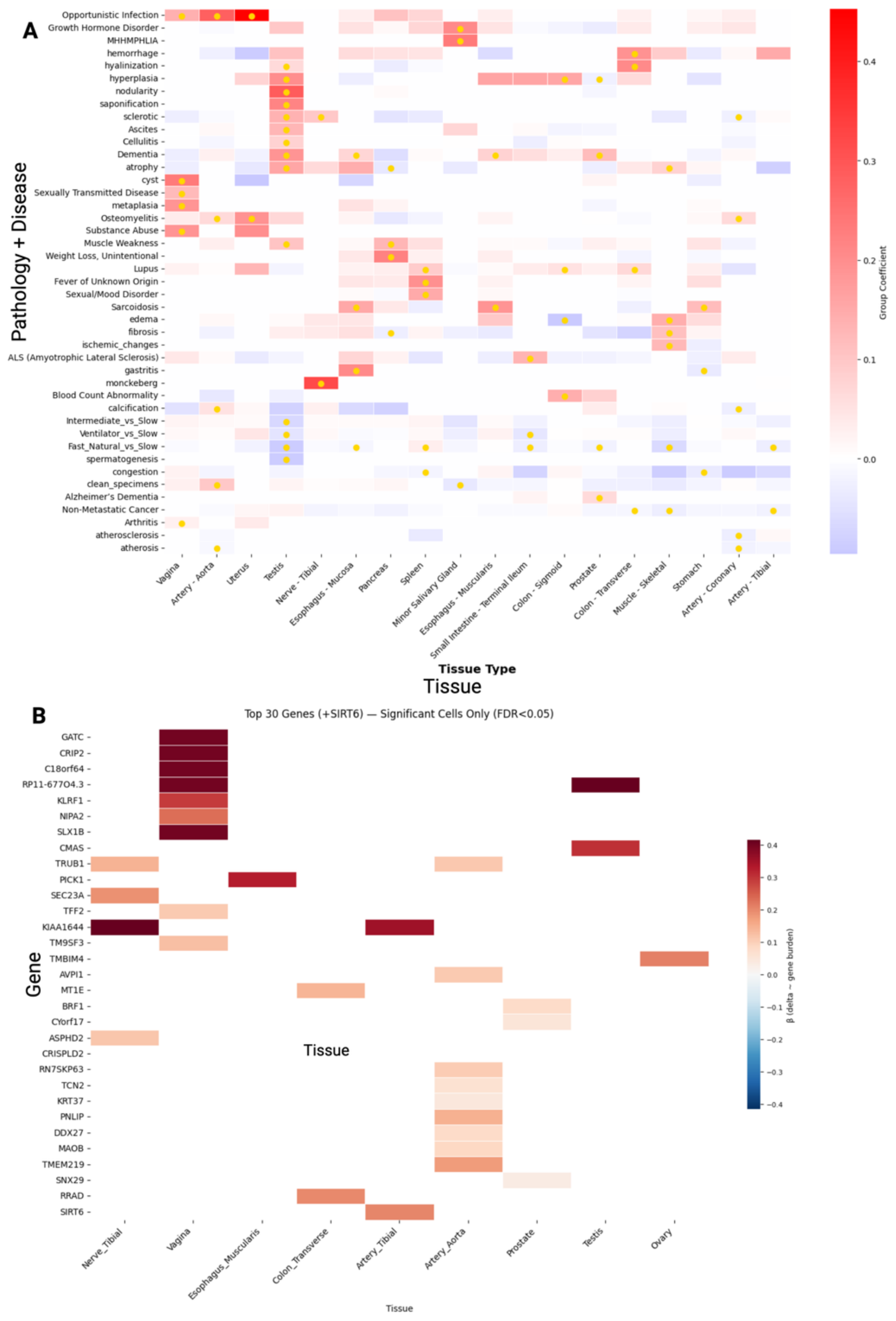


**Figure S21: Heatmaps showing associations between delta-SA scores (ΔSA) and health/genetic factors across tissue types. (A)** Multivariate regression coefficients for pathological conditions and post-mortem factors, with red indicating positive associations with accelerated remodeling and blue indicating negative associations. **(B)** Gene-level associations from germline variant analysis (n=970 individuals), showing significant associations (FDR P<0.1) between functional variants and tissue-specific remodeling rates. Color intensity represents effect size magnitude, with 17 genes are plotted here out of 226 meeting significance thresholds across tissues.

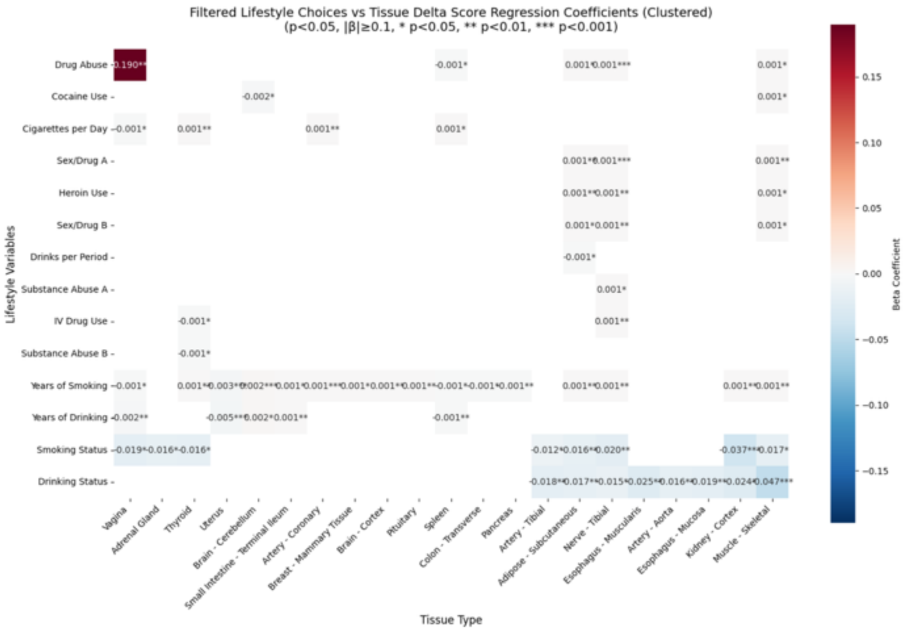


**Figure S22: Association between lifestyle factors and tissue-specific accelerated remodeling (ΔR) scores across GTEx samples.** Heatmap showing Mann-Whitney U test results comparing accelerated remodeling (ΔR) scores between individuals with and without various lifestyle characteristics across different tissue types. Lifestyle factors were extracted from GTEx phenotypic metadata and include smoking history, alcohol consumption, drug use, and physical activity patterns. For each lifestyle factor, samples were stratified into binary groups (presence vs. absence of the characteristic), and statistical associations were evaluated using the Mann-Whitney U test, a non-parametric method suitable for comparing distributions without assuming normality.

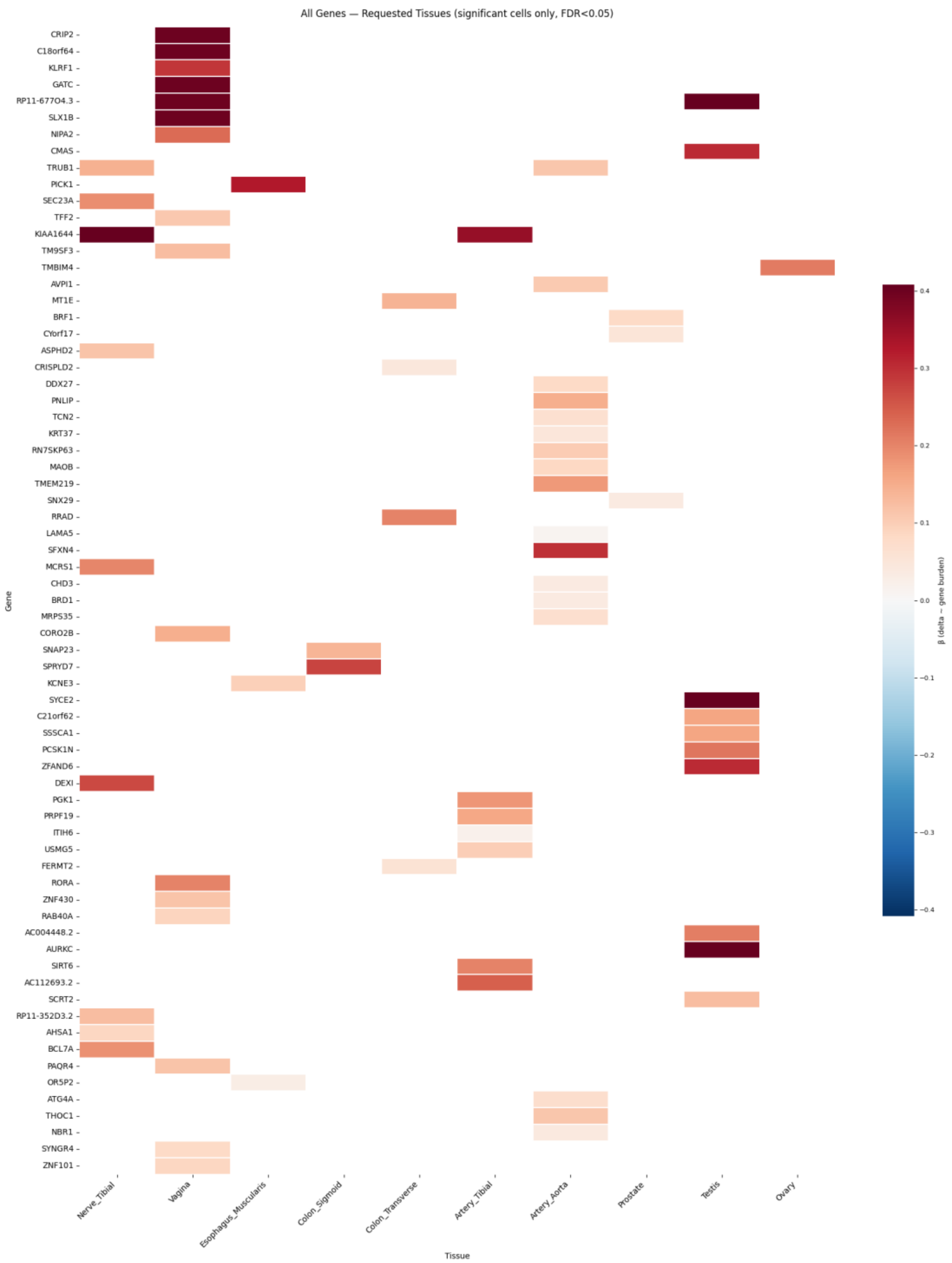


**Figure S23: Association between germline variants and remodeling score.** Focused on genes with statistically significant hit in at least one tissue, this figure provides association strength between germline variants and accelerated remodeling score, where red represents positive association and blue represents negative.

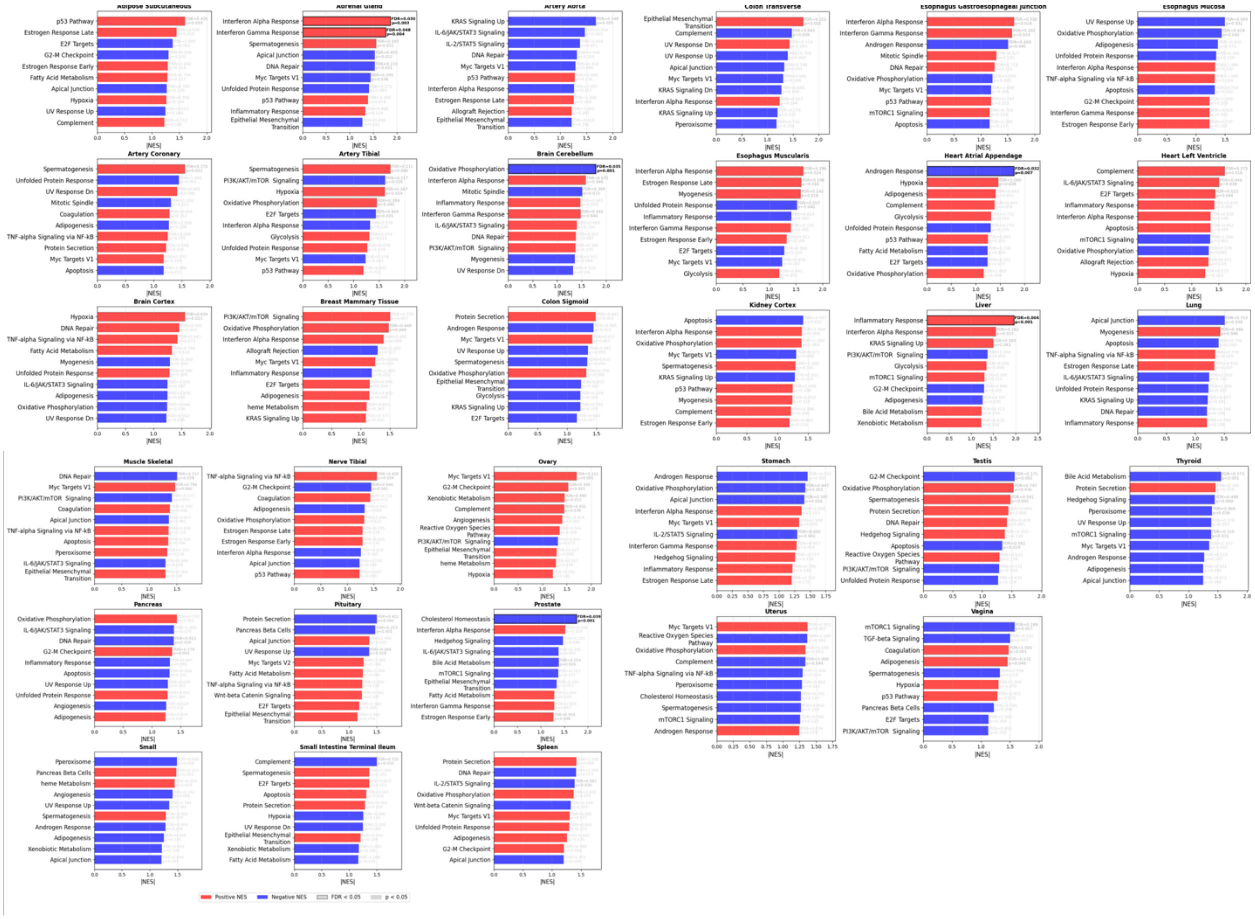


**Figure S24**: Tissue-specific Hallmark Pathways enriched in genes whose germline variants were associated with delta-SA score.

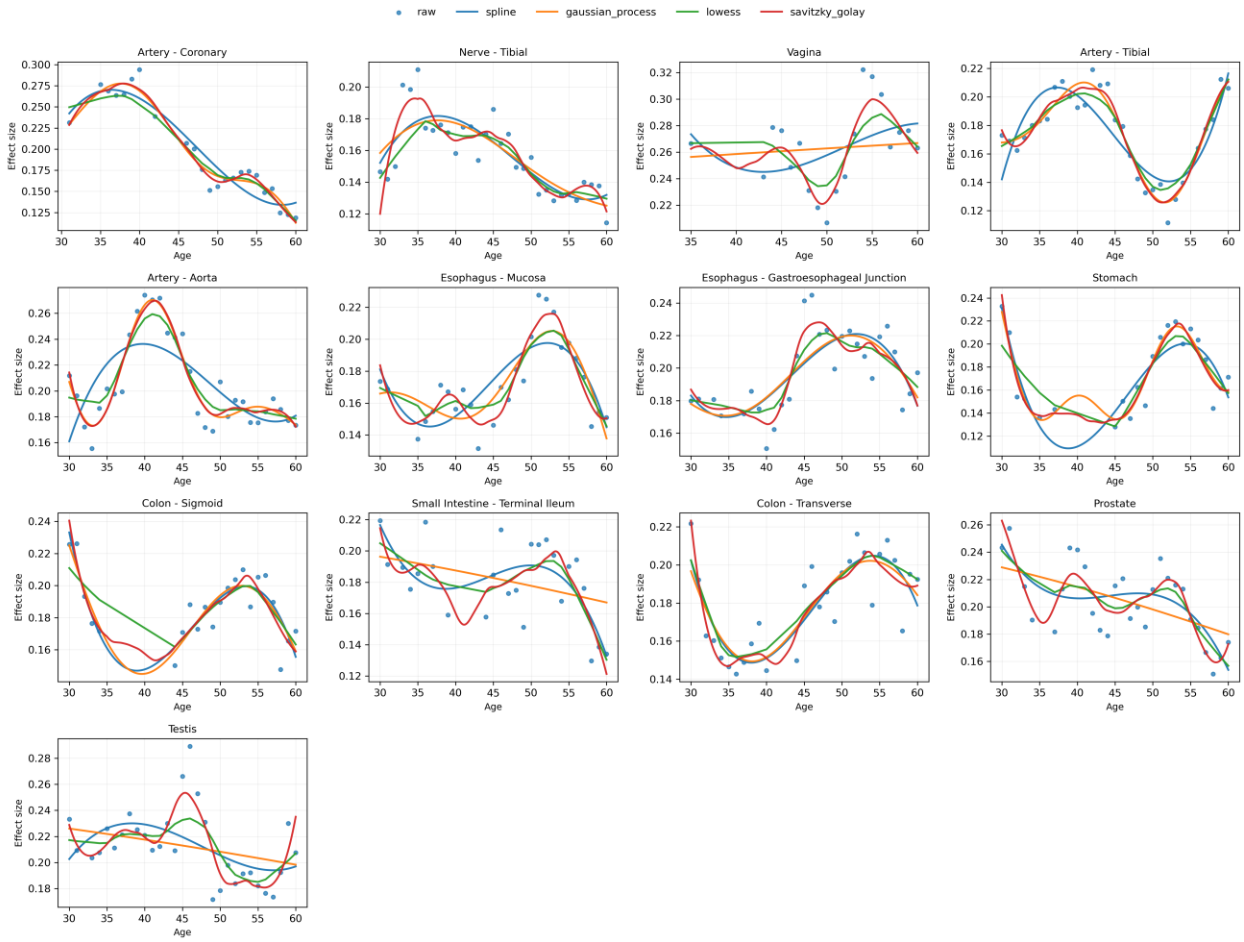


**Figure S25: Effect of Smoothing Methods on Raw Structural Aging Trajectories.** Raw Structural Aging Trajectories derived from UNI v1 embeddings are shown for 15 tissues: Artery (Aorta, Coronary, Tibial), Colon (Sigmoid, Transverse), Esophagus (Gastroesophageal Junction, Muscularis), Nerve (Tibial), Ovary, Prostate, Small Intestine (Terminal Ileum), Stomach, Testis, Uterus, and Vagina. Blue dots represent the raw effect size at each adjacent age transition, reflecting the magnitude of morphological change between consecutive age windows. To evaluate the influence of smoothing strategy on trajectory shape, four commonly used approaches were applied: cubic spline interpolation (blue), Gaussian process regression (orange), LOWESS (green), and Savitzky–Golay filtering (red). While all methods recover similar global trends, methods such as LOWESS and Savitzky–Golay introduce local fluctuations in some tissues, and Gaussian process smoothing can produce broader curvature depending on kernel assumptions. Cubic spline smoothing captures the main inflection points and periods of accelerated remodeling while avoiding unnecessary fluctuations. It provides a smooth and stable fit without overfitting the data. Based on this balance, spline fitting was used for primary PathStAR analyses.

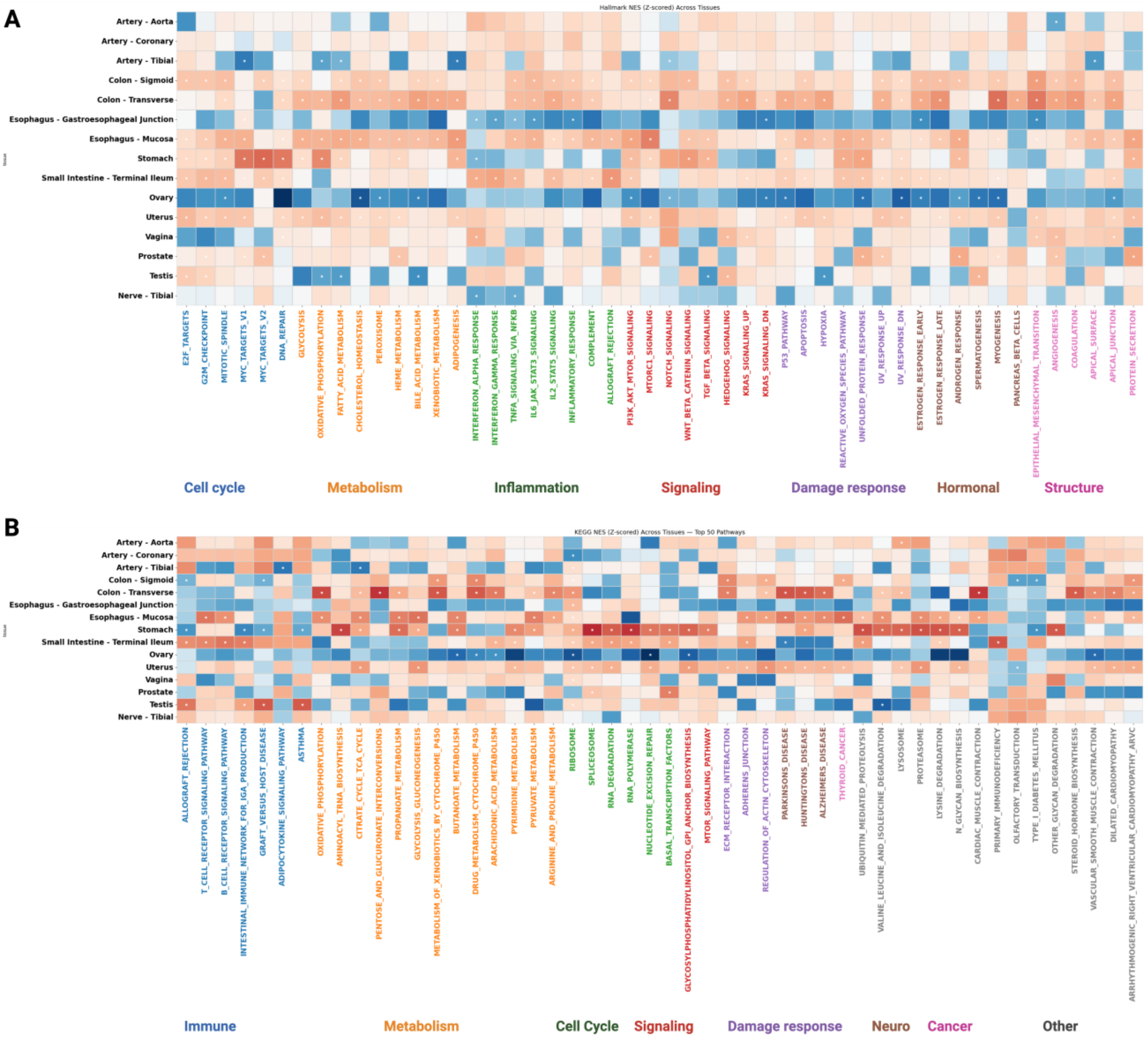


**Figure S26: Pathways enrichment of genes that follow structural aging trajectory.** **(A)** based on hallmarks pathways and **(B)** based on KEGG pathways. Red pathways are positively correlated with structural aging rate and vice versa for blue.
