## Supplementary Raw Trajectory for "Mapping Structural Aging of Human Tissue reveals tissue-specific trajectories and coordinated deterioration"

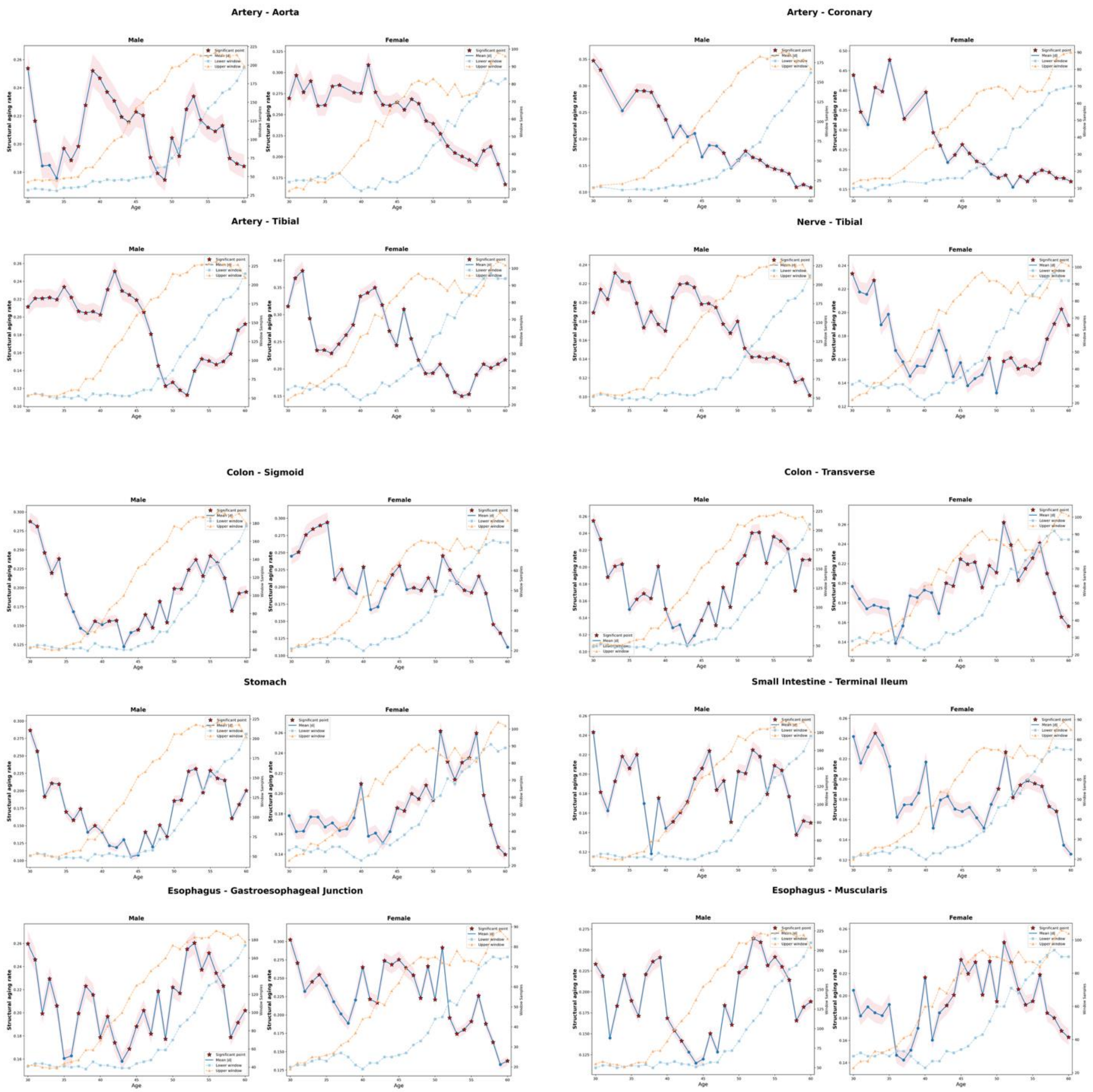


**Figure R1: Sex-stratified trajectories of structural aging across human tissues.** Line plots depict age-related structural aging rates, estimated as the mean absolute effect size between adjacent 10-year age windows, separately for male (left) and female (right) donors. Blue lines indicate effect sizes, with shaded regions showing the 95% confidence intervals obtained by bootstrap resampling. Red stars mark significant age transitions (≥5% of features with Welch's *t*-test $p < 0.05$). Orange and light-blue dashed lines indicate the sample counts contributing to the upper and lower age windows, respectively. Panels summarize 10 of the 15 tissues studied here; the remaining five tissues (Prostate, Testis, Ovary, Uterus, and Vagina) are sex-specific and are shown in a supplementary figure.

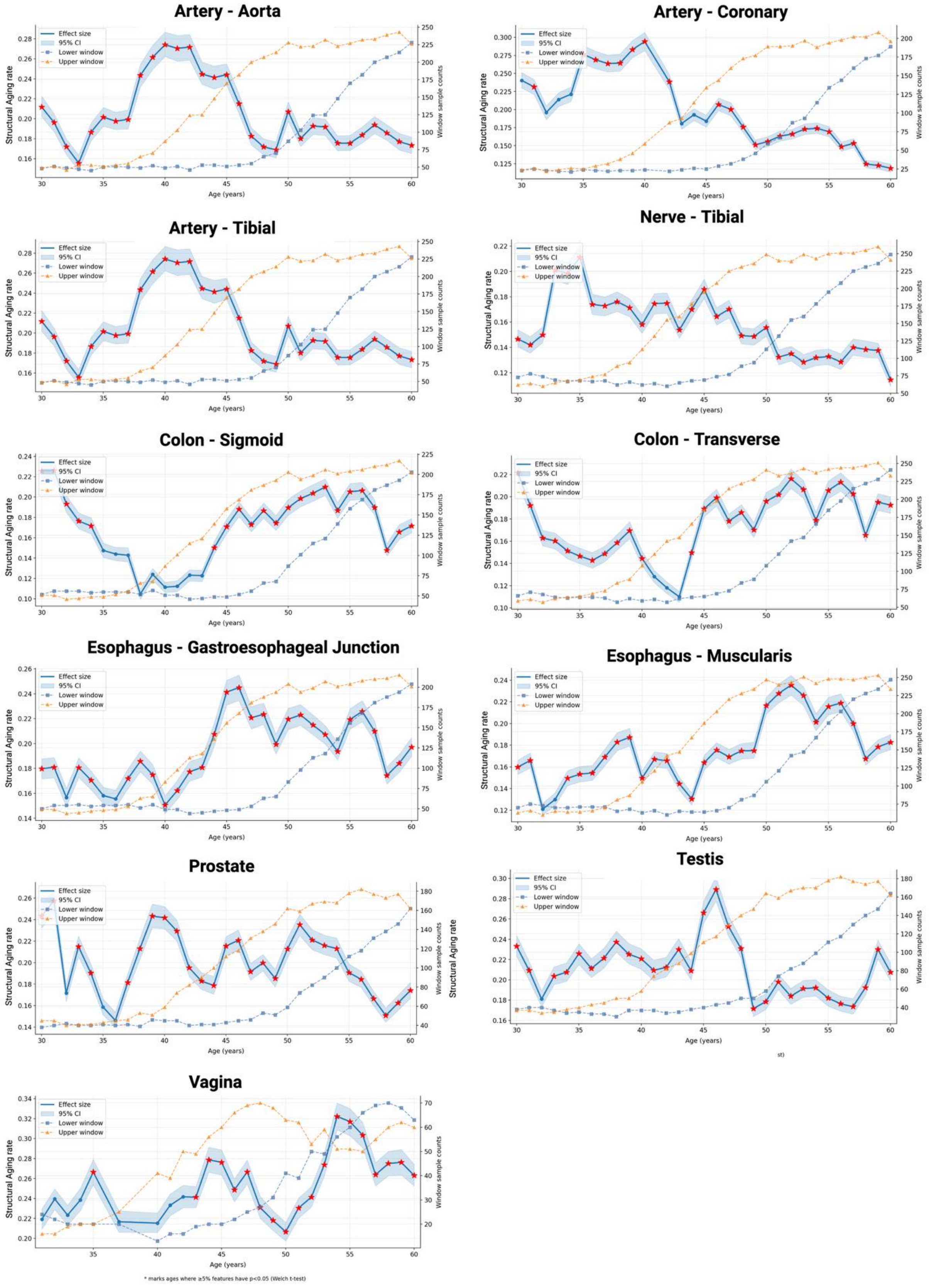


* marks ages where ≥5% features have p<0.05 (Welch t-test)

**Figure R2**: **No disease trajectory:** The plots show age-related structural aging rates quantified as the mean absolute effect size between adjacent 10-year age windows (blue line, left *y*-axis). Shaded regions represent the 95% confidence interval of the effect size, estimated by bootstrap resampling. Red stars indicate significant age transitions (≥5% of features with Welch's *t*-test $p < 0.05$). Sample counts contributing to each transition are shown for the lower (blue dashed line, right *y*-axis) and upper (orange dashed line, right *y*-axis) age windows. Each panel depicts trajectories for a distinct tissue, restricted to donors without major medical conditions or cancers, thereby capturing baseline, non-diseased structural aging dynamics.

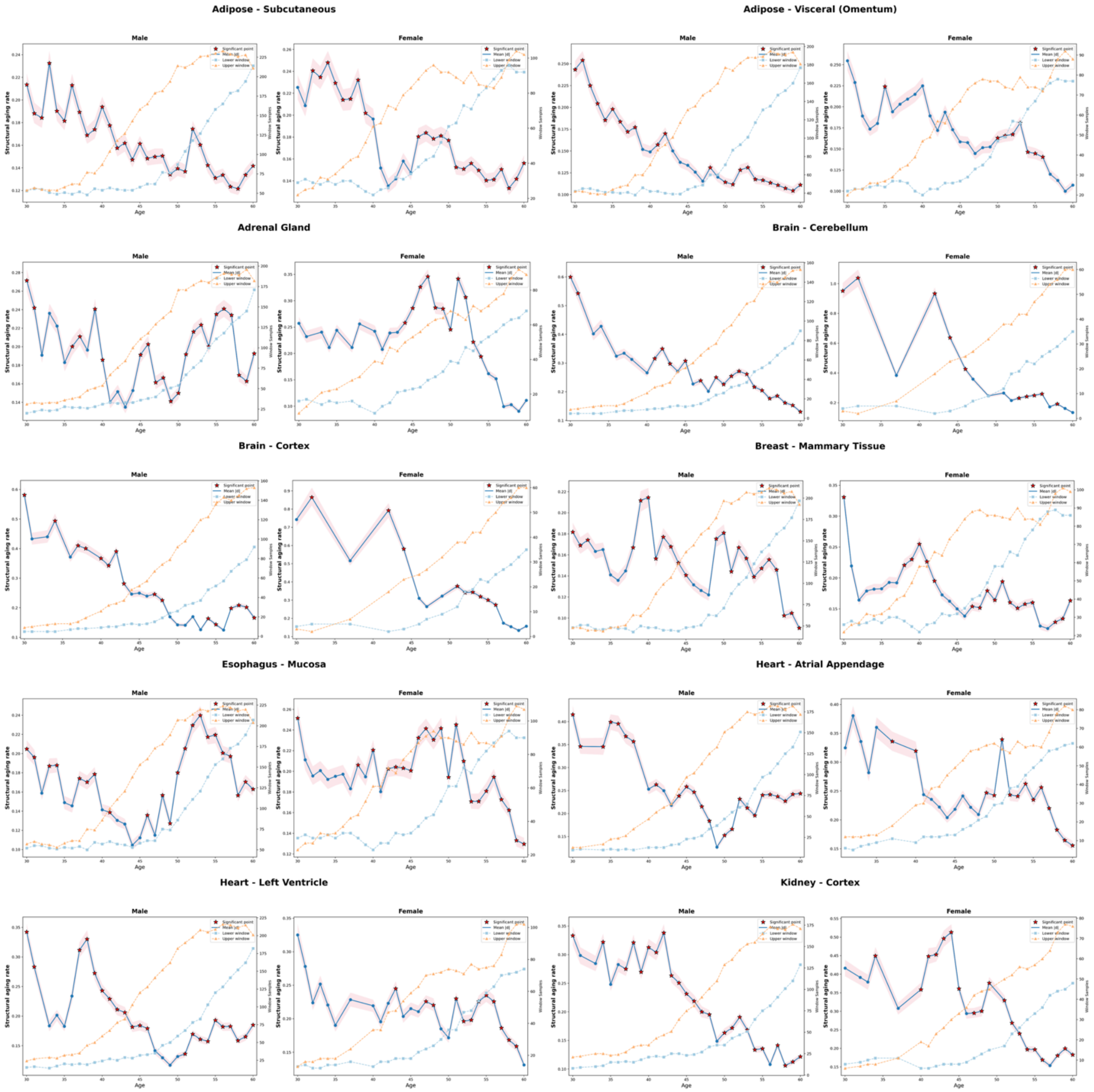
Adipose - Subcutaneous
Adipose - Visceral (Omentum)
Adrenal Gland
Brain - Cerebellum
Brain - Cortex
Breast - Mammary Tissue
Esophagus - Mucosa
Heart - Atrial Appendage
Heart - Left Ventricle
Kidney - Cortex
Male
Female
Structural aging rate
Age
Window Samples
Significant point
Mean |d|
Lower window
Upper window

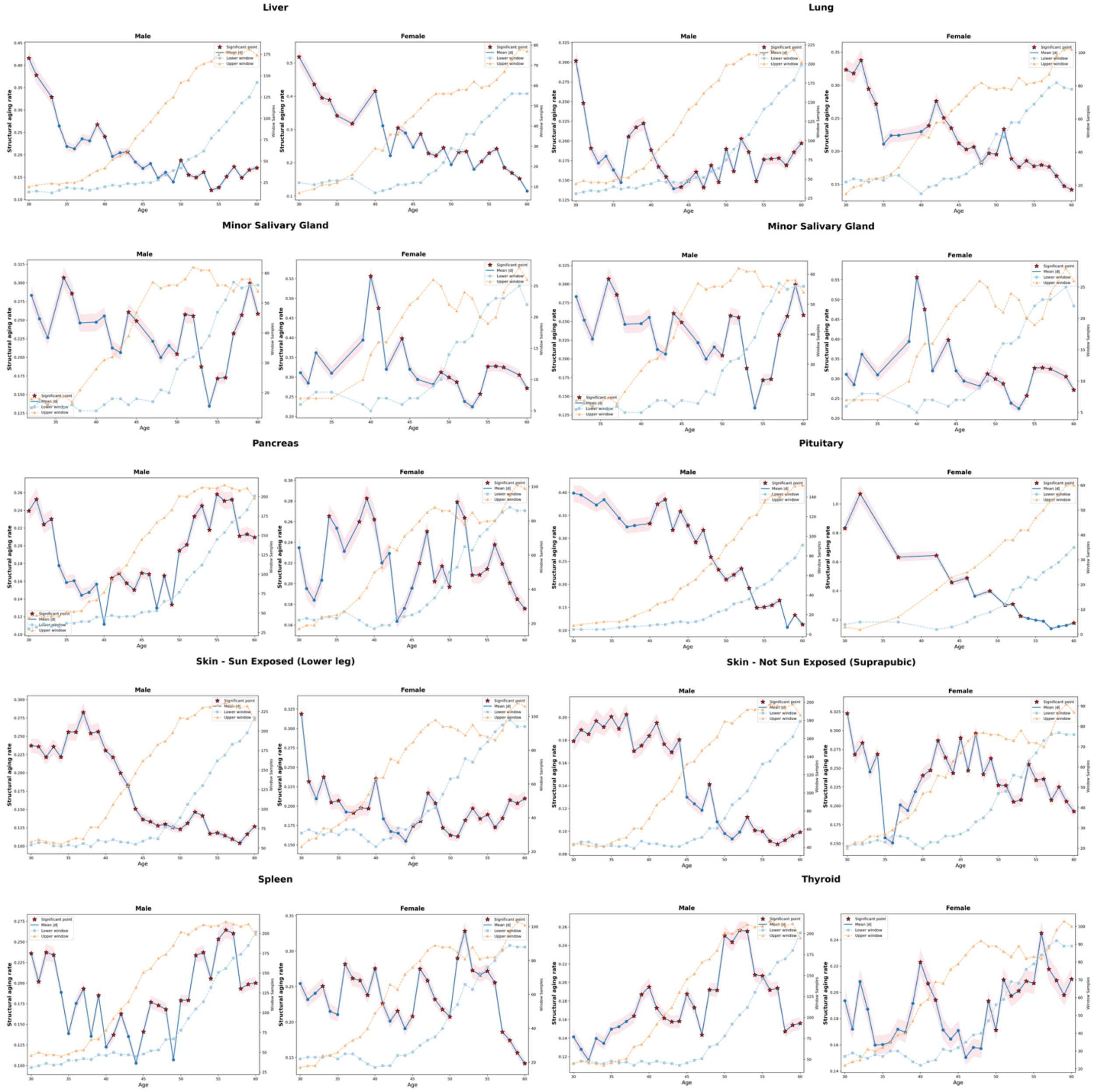


**Figure R3: Raw Structural Aging Analysis Across Additional Human Tissues with Sufficient Sample Size (n≥200) Stratified by Biological Sex**: This figure presents the raw, unsmoothed Structural Aging patterns across 20 additional human tissue types that met the minimum sample size requirement (n≥200) for PathStAR analysis, analyzed separately for male and female samples using the methodology described in Method 5.4. The tissues examined represent diverse organ systems including adipose tissue (Adipose - Subcutaneous, Adipose - Visceral [Omentum]), endocrine organs (Adrenal Gland, Pancreas, Pituitary, Thyroid), vascular structures (Artery - Aorta), central nervous system (Brain - Cerebellum, Brain - Cortex), mammary tissue (Breast - Mammary Tissue), gastrointestinal components (Esophagus - Mucosa, Liver), cardiovascular system (Heart

- Atrial Appendage, Heart - Left Ventricle), renal tissue (Kidney - Cortex), respiratory system (Lung), musculoskeletal system (Muscle - Skeletal), integumentary system (Skin - Not Sun Exposed [Suprapubic], Skin - Sun Exposed [Lower leg]), and lymphoid tissue (Spleen).(See Method 5.3)

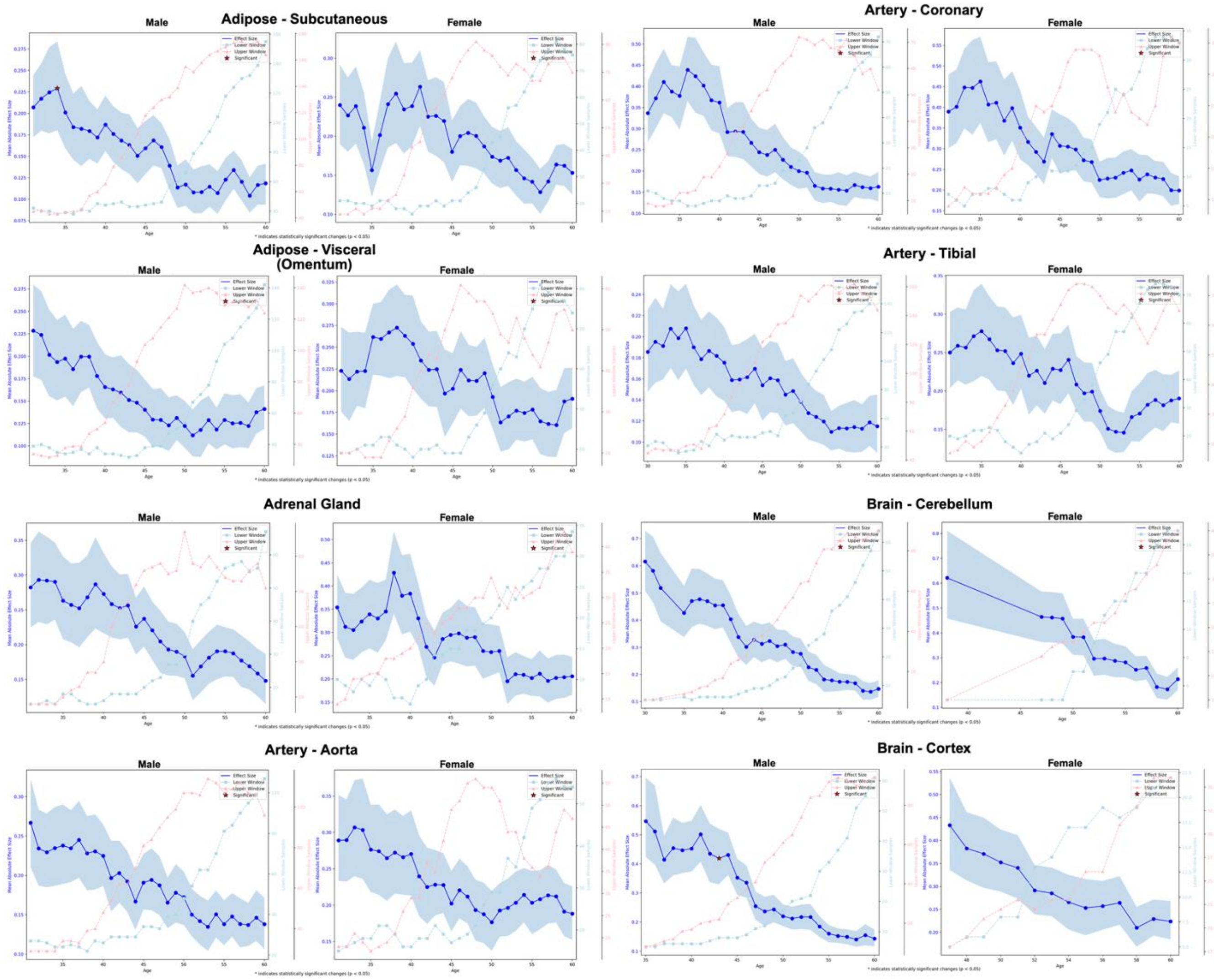
Adipose - Subcutaneous
Male
Female
Artery - Coronary
Male
Female
Adipose - Visceral (Omentum)
Male
Female
Artery - Tibial
Male
Female
Adrenal Gland
Male
Female
Brain - Cerebellum
Male
Female
Artery - Aorta
Male
Female
Brain - Cortex
Male
Female
Effect Size
Lower Window
Upper Window
Significant
Mean Absolute Effect Size
Age
* indicates statistically significant changes (p < 0.05)

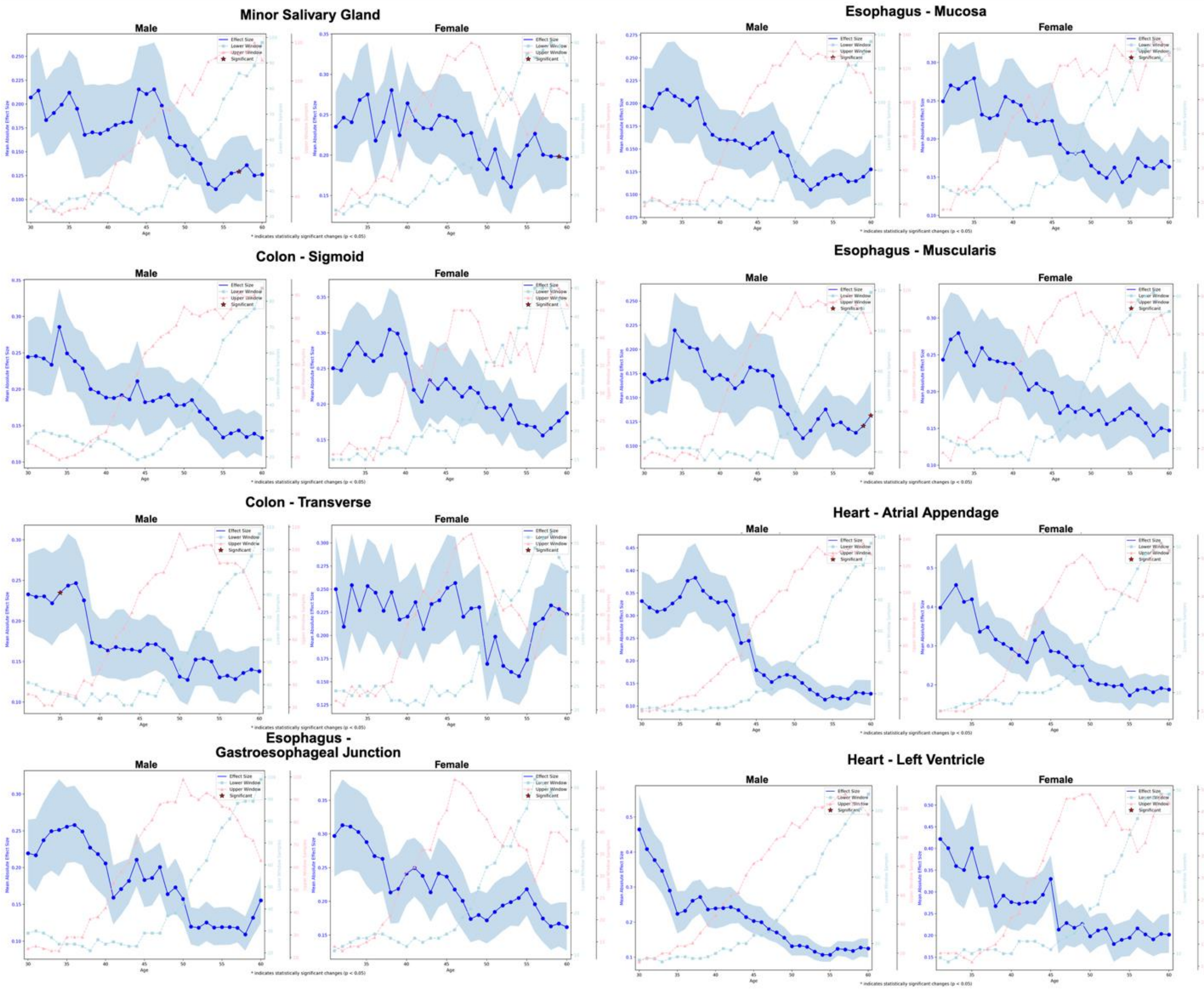

Minor Salivary Gland
Esophagus - Mucosa
Colon - Sigmoid
Esophagus - Muscularis
Colon - Transverse
Heart - Atrial Appendage
Esophagus -
Gastroesophageal Junction
Heart - Left Ventricle
Male
Female
Effect Size
Lower Window
Upper Window
Significant
Mean Absolute Effect Size
Age
* indicates statistically significant changes (p < 0.05)

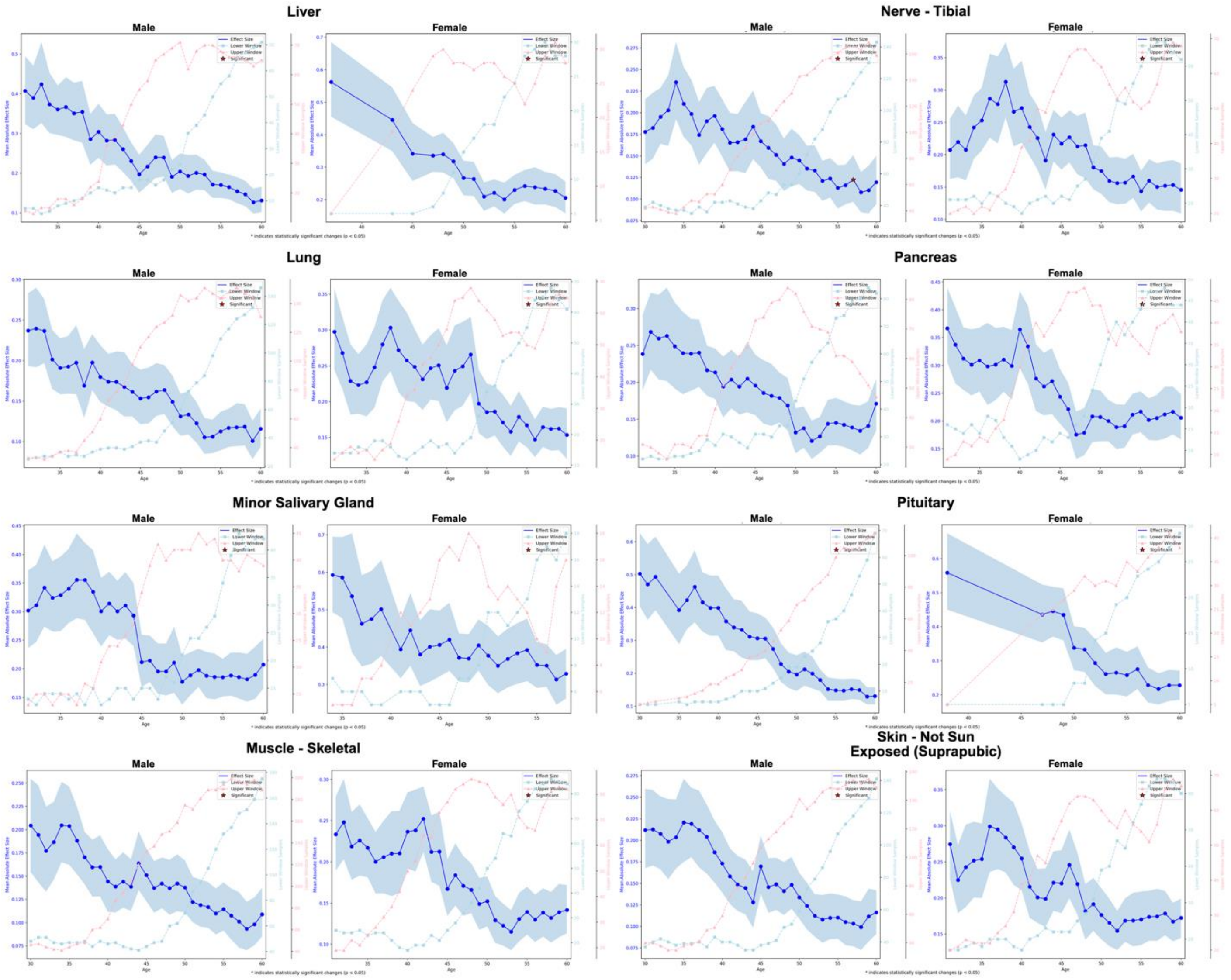
Liver
Nerve - Tibial
Lung
Pancreas
Minor Salivary Gland
Pituitary
Muscle - Skeletal
Skin - Not Sun
Exposed (Suprapubic)
Male
Female
Effect Size
Lower Window
Upper Window
Significant
Mean Absolute Effect Size
Age
* indicates statistically significant changes (p < 0.05)

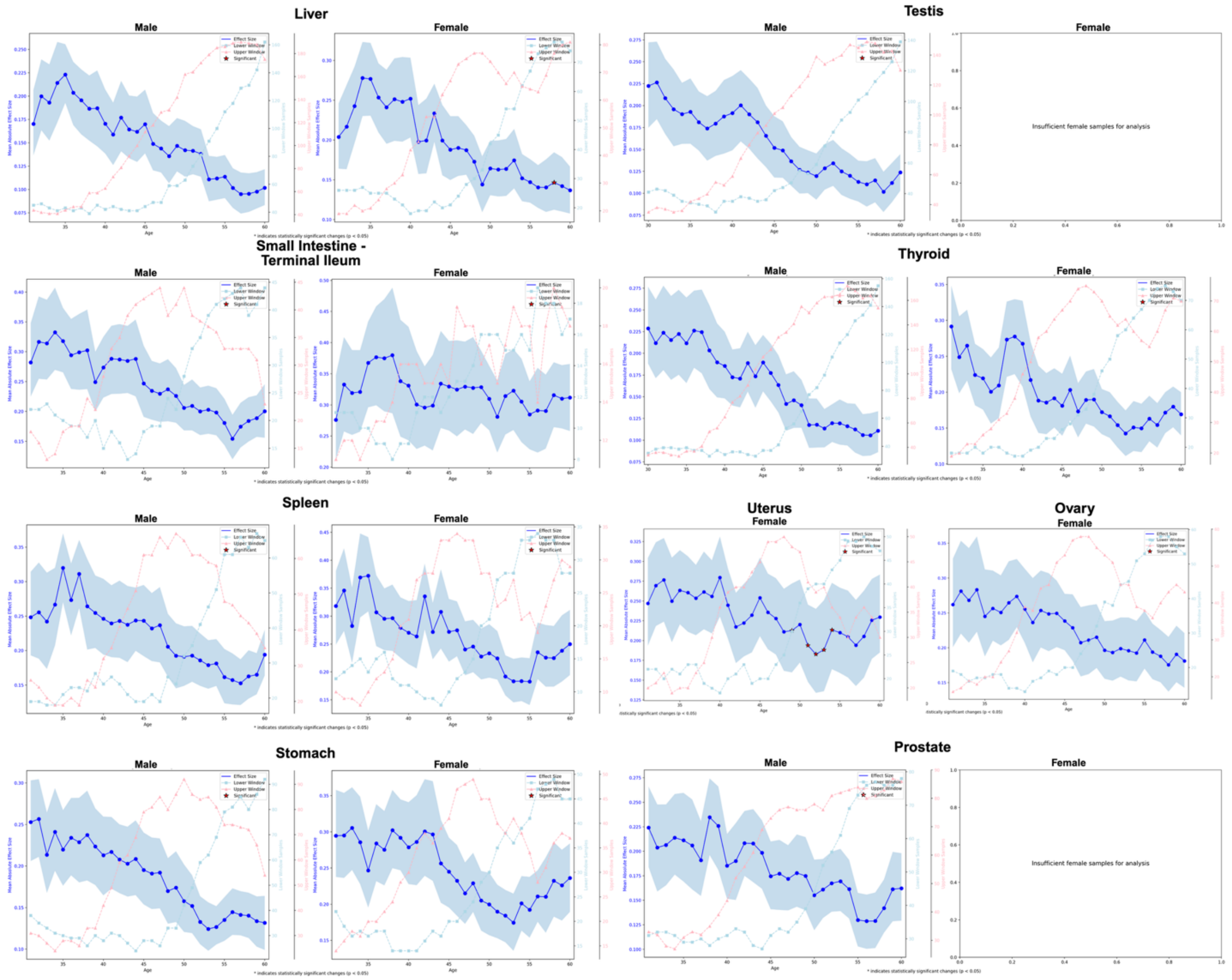


**Figure R4: Transcriptomic Aging Rate Trajectories Across Human Tissues Stratified by Biological Sex.** Sliding window analysis applied to transcriptomic principal component features across 58 tissue-sex combinations (29 tissues with sufficient samples for both sexes, plus 4 sex-specific tissues) to quantify the rate of molecular change during aging. Gene expression data (TPMs) were log1p-transformed and subjected to tissue-specific PCA using the top 5,000 most variable genes reduced to 50 principal components. Using 10-year sliding windows across ages 30-79, the magnitude of change in transcriptomic PC space between consecutive age periods was computed to derive tissue-specific and sex-specific aging rate trajectories.

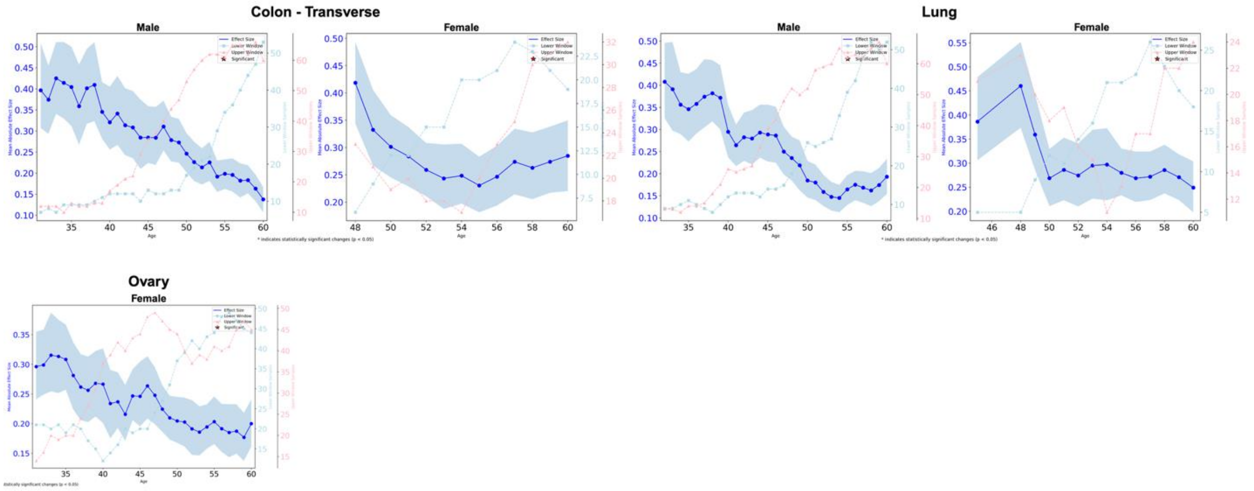


**Figure R5: DNA Methylation Aging Rate Trajectories Across Human Tissues Stratified by Biological Sex.** Sliding window analysis applied to DNA methylation principal component features across 5 tissue-sex combinations (Colon-Transverse and Lung for both sexes, Ovary for females only) to quantify the rate of epigenetic change during aging. Methylation beta values were normalized, converted to M-values, filtered for <5% missing data per CpG, imputed with per-CpG medians, and reduced to 50 principal components using the top 10,000 most variable CpG sites per tissue. Using 10-year sliding windows across ages 30-79, the magnitude of change in methylation PC space between consecutive age periods was computed to derive tissue-specific and sex-specific aging rate trajectories.

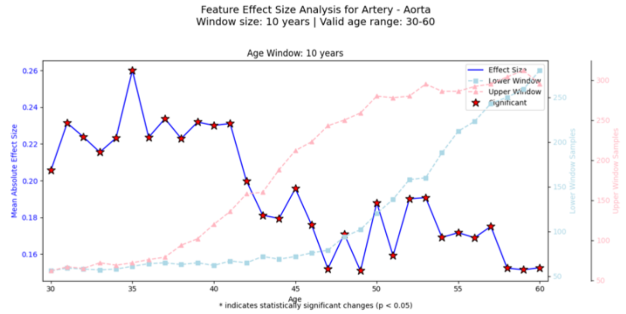
Feature Effect Size Analysis for Artery - Aorta
Window size: 10 years | Valid age range: 30-60
Age Window: 10 years
Effect Size
Lower Window
Upper Window
Significant
Mean Absolute Effect Size
Lower Window Samples
Upper Window Samples
Age
* indicates statistically significant changes (p < 0.05)

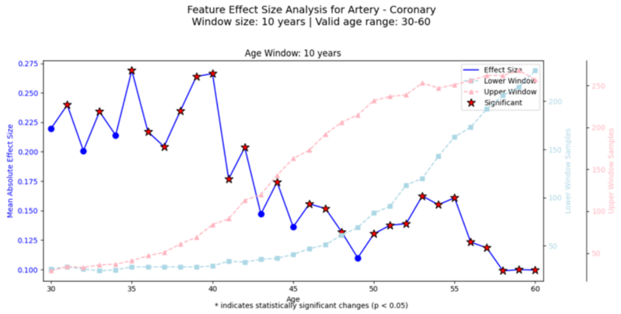
Feature Effect Size Analysis for Artery - Coronary
Window size: 10 years | Valid age range: 30-60
Age Window: 10 years
Effect Size
Lower Window
Upper Window
Significant
Mean Absolute Effect Size
Lower Window Samples
Upper Window Samples
Age
* indicates statistically significant changes (p < 0.05)

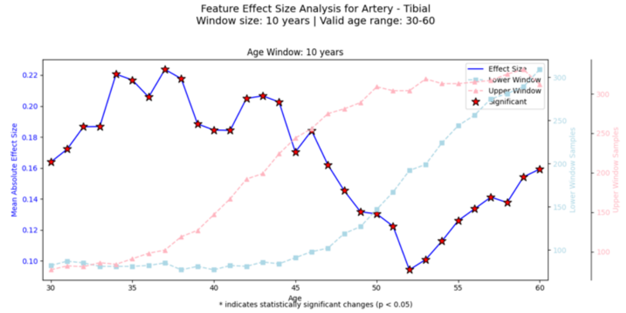
Feature Effect Size Analysis for Artery - Tibial
Window size: 10 years | Valid age range: 30-60
Age Window: 10 years
Effect Size
Lower Window
Upper Window
Significant
Mean Absolute Effect Size
Lower Window Samples
Upper Window Samples
Age
* indicates statistically significant changes (p < 0.05)

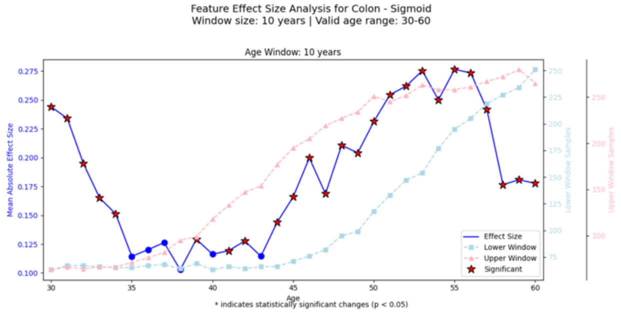
Feature Effect Size Analysis for Colon - Sigmoid
Window size: 10 years | Valid age range: 30-60
Age Window: 10 years
Effect Size
Lower Window
Upper Window
Significant
Mean Absolute Effect Size
Lower Window Samples
Upper Window Samples
Age
* indicates statistically significant changes (p < 0.05)

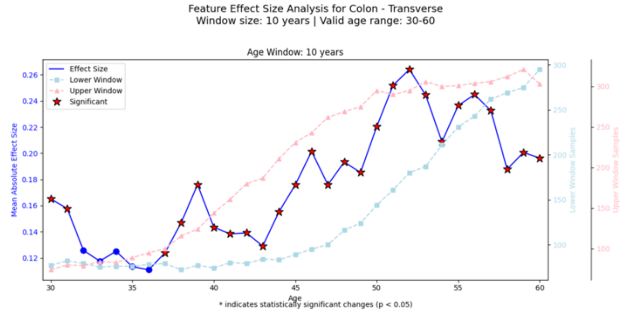
Feature Effect Size Analysis for Colon - Transverse
Window size: 10 years | Valid age range: 30-60
Age Window: 10 years
Effect Size
Lower Window
Upper Window
Significant
Mean Absolute Effect Size
Lower Window Samples
Upper Window Samples
Age
* indicates statistically significant changes (p < 0.05)

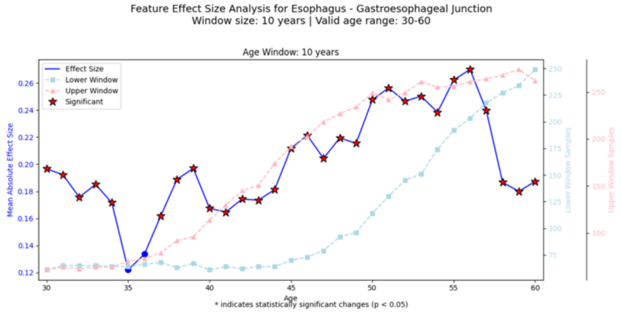
Feature Effect Size Analysis for Esophagus - Gastroesophageal Junction
Window size: 10 years | Valid age range: 30-60
Age Window: 10 years
Effect Size
Lower Window
Upper Window
Significant
Mean Absolute Effect Size
Lower Window Samples
Upper Window Samples
Age
* indicates statistically significant changes (p < 0.05)

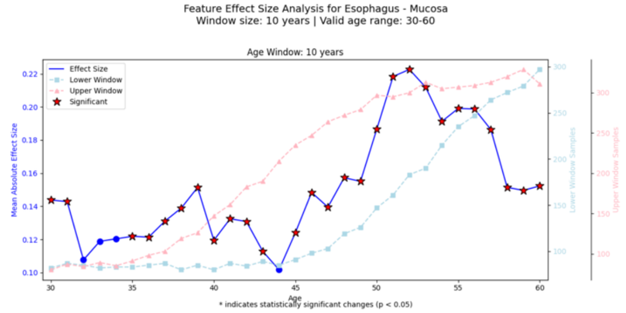
Feature Effect Size Analysis for Esophagus - Mucosa
Window size: 10 years | Valid age range: 30-60
Age Window: 10 years
Effect Size
Lower Window
Upper Window
Significant
Mean Absolute Effect Size
Lower Window Samples
Upper Window Samples
Age
* indicates statistically significant changes (p < 0.05)

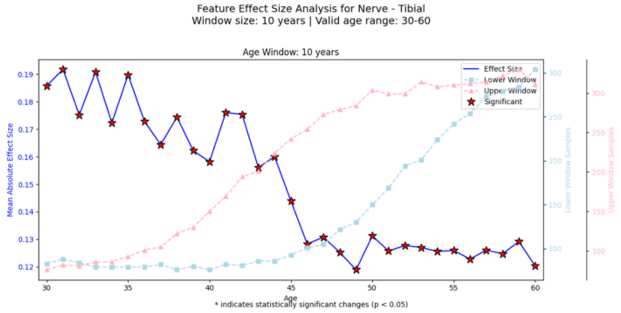
Feature Effect Size Analysis for Nerve - Tibial
Window size: 10 years | Valid age range: 30-60
Age Window: 10 years
Effect Size
Lower Window
Upper Window
Significant
Mean Absolute Effect Size
Lower Window Samples
Upper Window Samples
Age
* indicates statistically significant changes (p < 0.05)

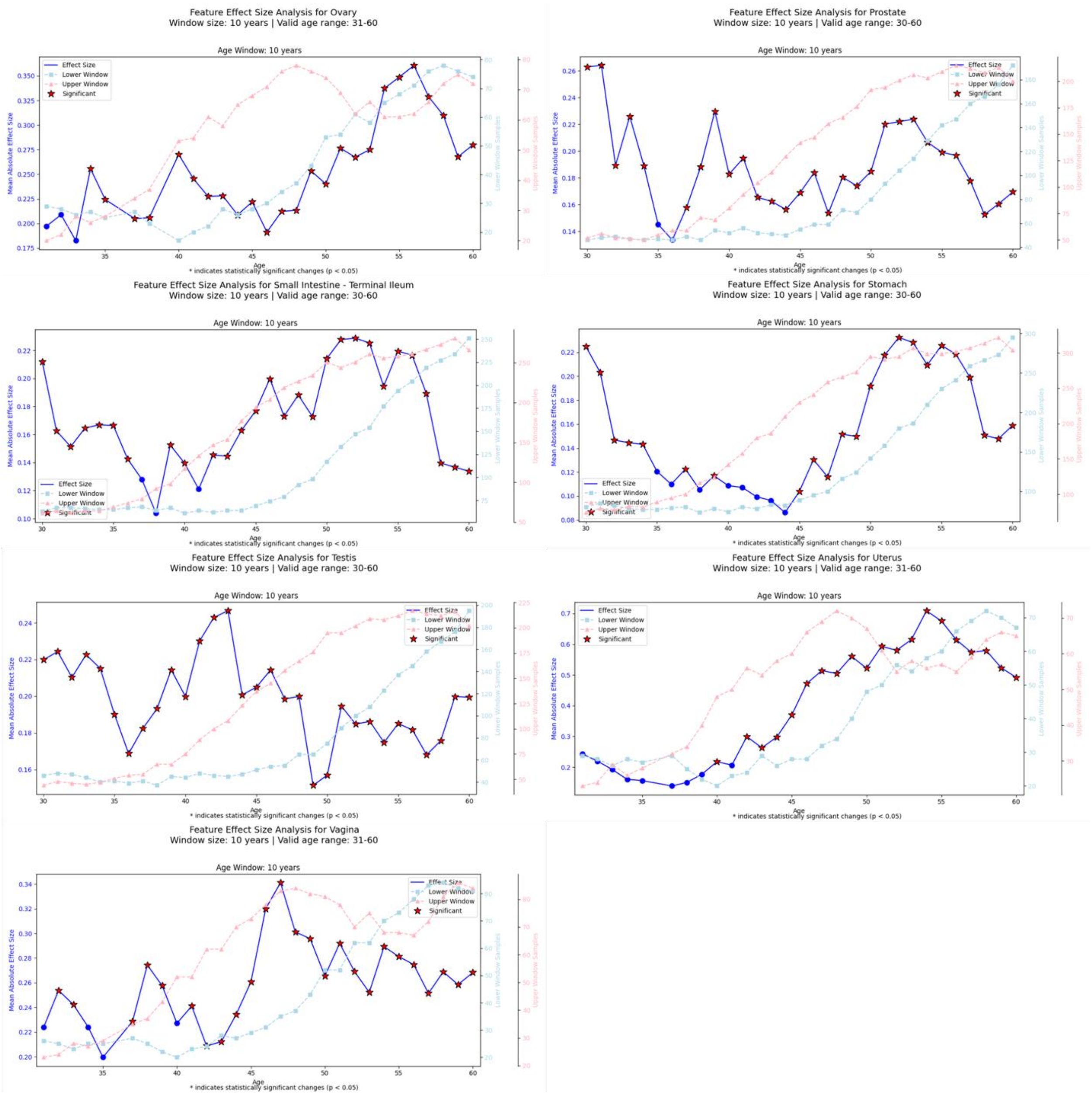


**Figure R6: Raw Structural Aging Trajectories using UNI v2**

Raw Structural Aging Trajectories (blue lines) are shown for the same 15 tissues as in the main analysis. Morphological change between adjacent age windows was computed using slide-level embeddings extracted from **UNI v2**, an updated Vision Transformer–based pathology foundation model trained via large-scale self-supervised learning on diverse whole-slide images. UNI v2 produces high-capacity 1536-dimensional patch embeddings, which were aggregated to construct structural representations for each sample before applying the PathStAR pipeline.

Red stars denote age transitions where ≥5% of features exhibit nominal significance ($p < 0.05$). Dashed curves indicate the number of samples contributing to the lower (squares) and upper (triangles) age windows. The persistence of discrete accelerated structural aging phases across tissues using UNI v2 confirms that PathStAR captures biologically meaningful tissue remodeling patterns independent of embedding dimensionality or specific pretraining strategy.

Feature Effect Size Analysis for Artery - Aorta
Window size: 10 years | Valid age range: 30-60

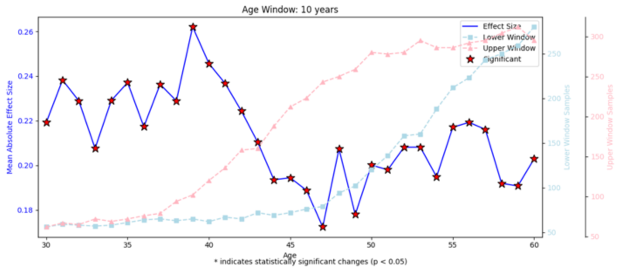

* indicates statistically significant changes (p < 0.05)

Feature Effect Size Analysis for Artery - Coronary
Window size: 10 years | Valid age range: 30-60

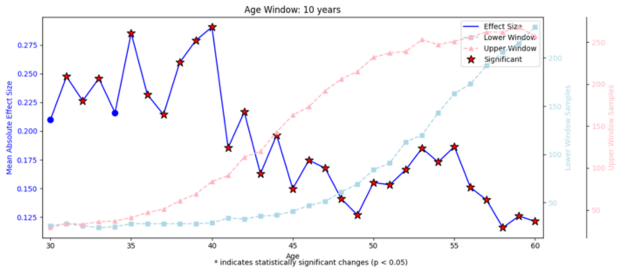

* indicates statistically significant changes (p < 0.05)

Feature Effect Size Analysis for Artery - Tibial
Window size: 10 years | Valid age range: 30-60

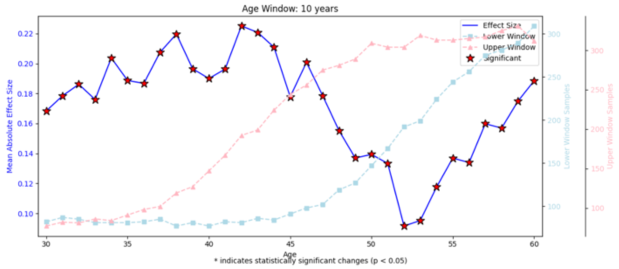

* indicates statistically significant changes (p < 0.05)

Feature Effect Size Analysis for Colon - Sigmoid
Window size: 10 years | Valid age range: 30-60

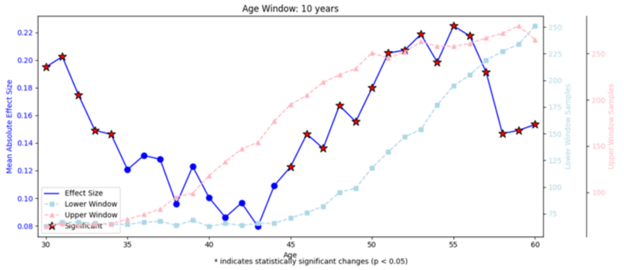

* indicates statistically significant changes (p < 0.05)

Feature Effect Size Analysis for Colon - Transverse
Window size: 10 years | Valid age range: 30-60

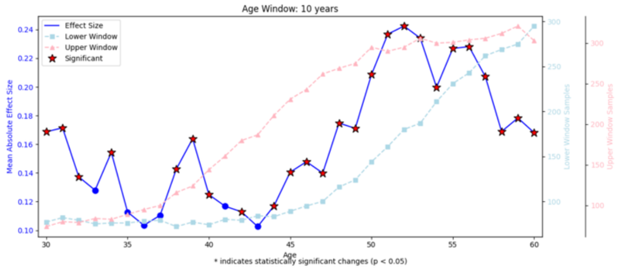

* indicates statistically significant changes (p < 0.05)

Feature Effect Size Analysis for Esophagus - Gastroesophageal Junction
Window size: 10 years | Valid age range: 30-60

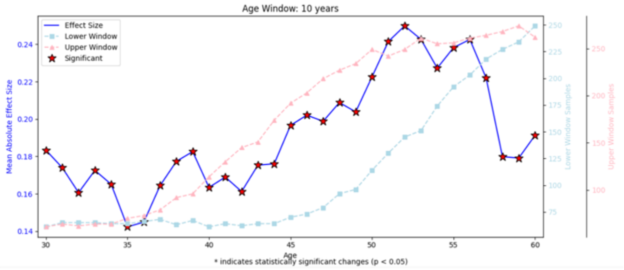

* indicates statistically significant changes (p < 0.05)

Feature Effect Size Analysis for Esophagus - Mucosa
Window size: 10 years | Valid age range: 30-60

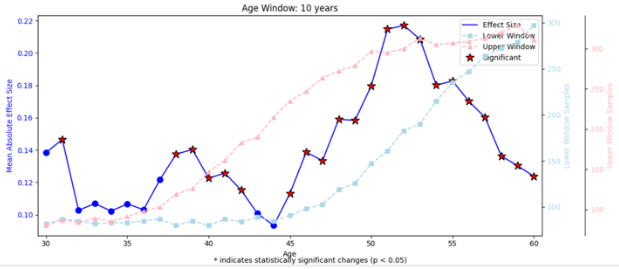

* indicates statistically significant changes (p < 0.05)

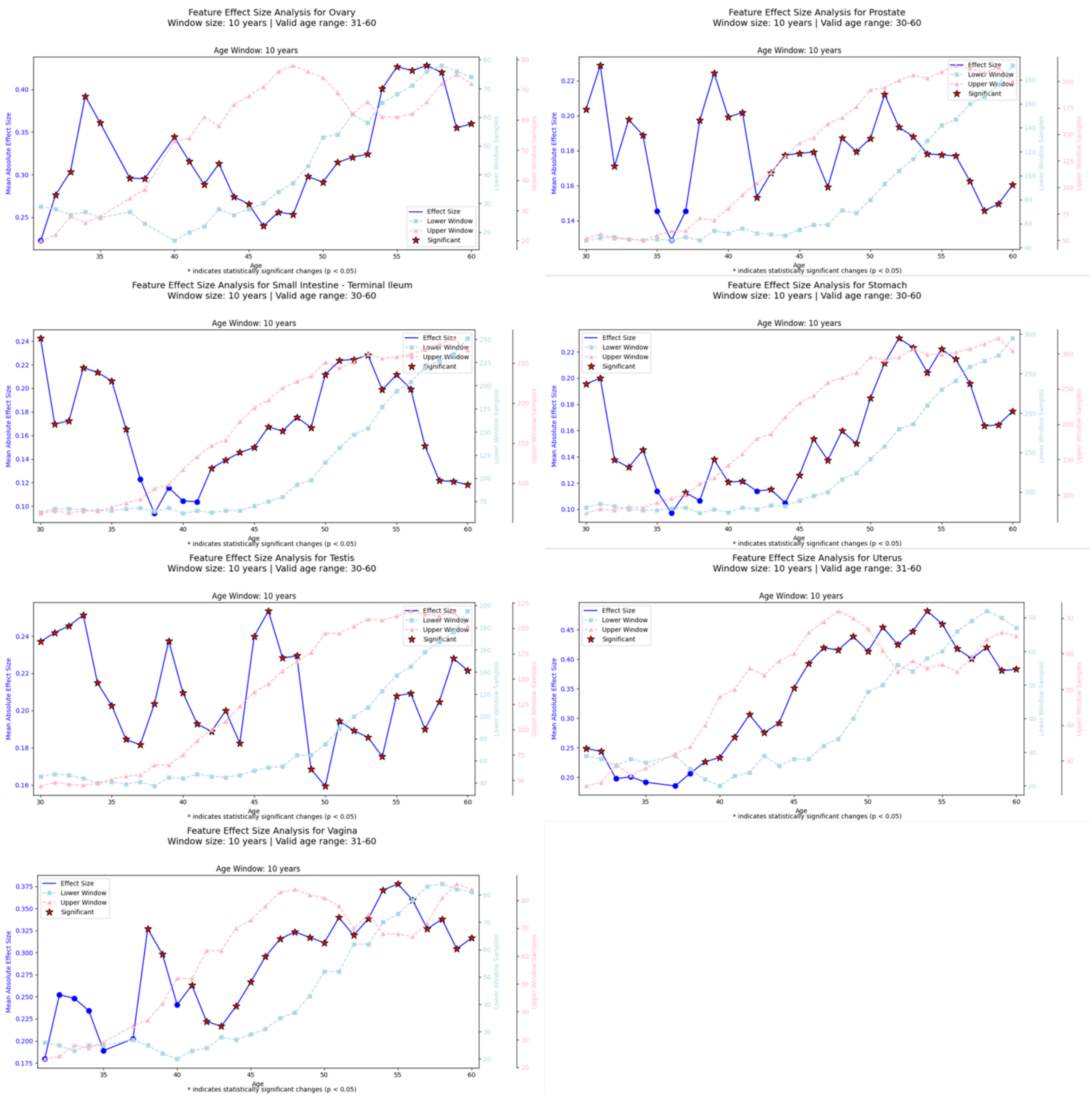


**Fig R7 : Raw Structural Aging Trajectories using CONCH v1.5.** Raw Structural Aging Trajectories (blue lines) are shown for the same 15 tissues: Artery (Aorta, Coronary, Tibial), Colon (Sigmoid, Transverse), Esophagus (Gastroesophageal Junction, Muscularis), Nerve (Tibial), Ovary, Prostate, Small Intestine (Terminal Ileum), Stomach, Testis, Uterus, and Vagina. Each trajectory represents the magnitude of morphological change between adjacent age windows, computed using slide-level representations derived from **CONCH v1.5**, a vision-language pathology foundation model trained on large-scale histopathology datasets with contrastive supervision. Patch embeddings (768-dimensional) were extracted and aggregated to generate tissue-level structural representations prior to PathStAR analysis.

Red stars denote age transitions where ≥5% of features exhibit nominal significance ($p < 0.05$). Dashed curves on the secondary axis show the number of samples contributing to the lower (squares) and upper (triangles) age windows at each transition. The recapitulation of tissue-specific accelerated periods using CONCH demonstrates that structural aging trajectories are robust to the choice of feature extractor and not dependent on a single pretrained model.

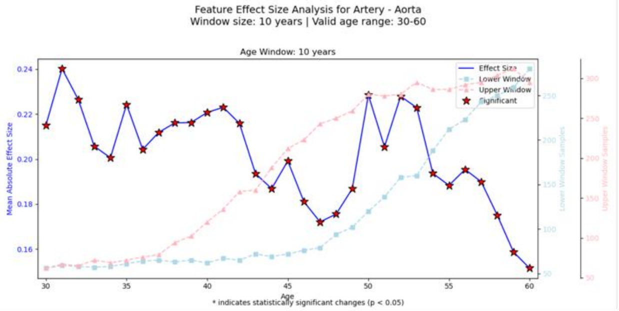
Feature Effect Size Analysis for Artery - Aorta
Window size: 10 years | Valid age range: 30-60
Age Window: 10 years
Effect Size
Lower Window
Upper Window
Significant
Mean Absolute Effect Size
Lower Window Samples
Upper Window Samples
Age
* indicates statistically significant changes (p < 0.05)

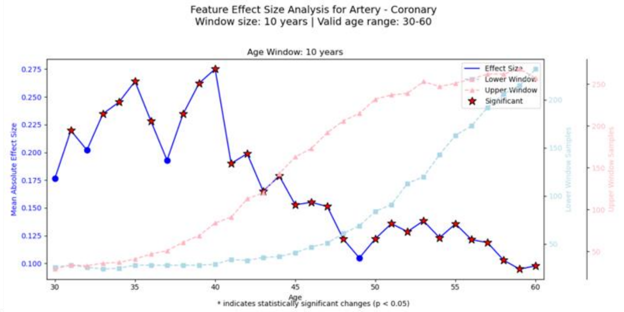
Feature Effect Size Analysis for Artery - Coronary
Window size: 10 years | Valid age range: 30-60
Age Window: 10 years
Effect Size
Lower Window
Upper Window
Significant
Mean Absolute Effect Size
Lower Window Samples
Upper Window Samples
Age
* indicates statistically significant changes (p < 0.05)

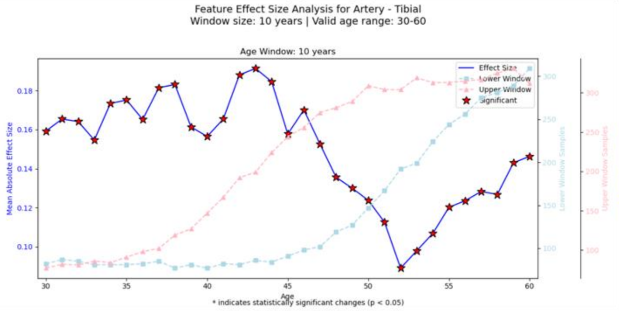
Feature Effect Size Analysis for Artery - Tibial
Window size: 10 years | Valid age range: 30-60
Age Window: 10 years
Effect Size
Lower Window
Upper Window
Significant
Mean Absolute Effect Size
Lower Window Samples
Upper Window Samples
Age
* indicates statistically significant changes (p < 0.05)

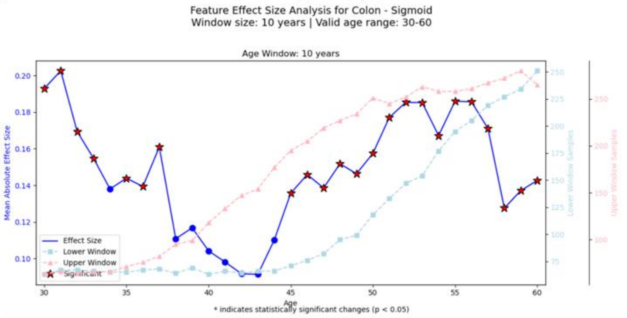
Feature Effect Size Analysis for Colon - Sigmoid
Window size: 10 years | Valid age range: 30-60
Age Window: 10 years
Effect Size
Lower Window
Upper Window
Significant
Mean Absolute Effect Size
Lower Window Samples
Upper Window Samples
Age
* indicates statistically significant changes (p < 0.05)

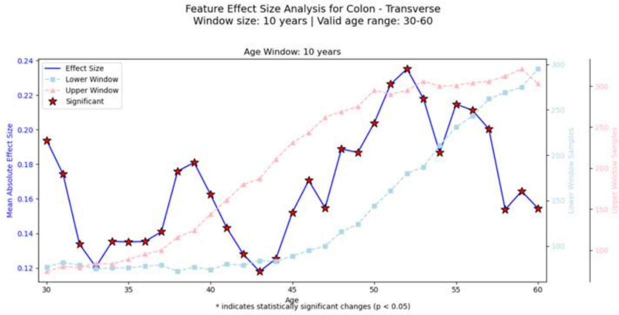
Feature Effect Size Analysis for Colon - Transverse
Window size: 10 years | Valid age range: 30-60
Age Window: 10 years
Effect Size
Lower Window
Upper Window
Significant
Mean Absolute Effect Size
Lower Window Samples
Upper Window Samples
Age
* indicates statistically significant changes (p < 0.05)

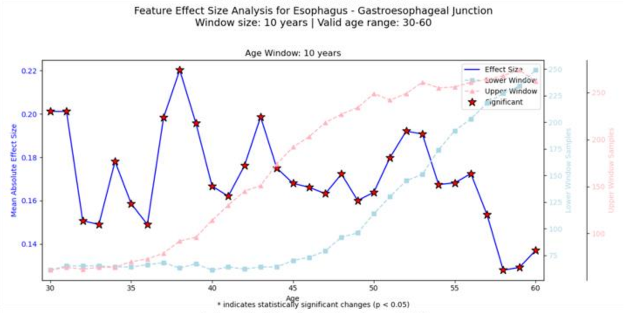
Feature Effect Size Analysis for Esophagus - Gastroesophageal Junction
Window size: 10 years | Valid age range: 30-60
Age Window: 10 years
Effect Size
Lower Window
Upper Window
Significant
Mean Absolute Effect Size
Lower Window Samples
Upper Window Samples
Age
* indicates statistically significant changes (p < 0.05)

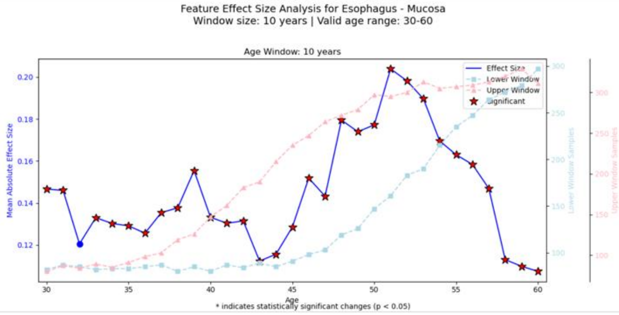
Feature Effect Size Analysis for Esophagus - Mucosa
Window size: 10 years | Valid age range: 30-60
Age Window: 10 years
Effect Size
Lower Window
Upper Window
Significant
Mean Absolute Effect Size
Lower Window Samples
Upper Window Samples
Age
* indicates statistically significant changes (p < 0.05)

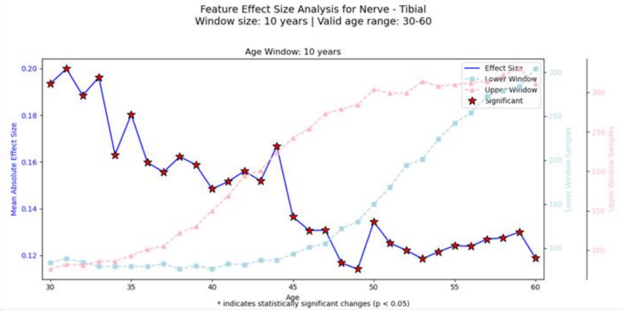
Feature Effect Size Analysis for Nerve - Tibial
Window size: 10 years | Valid age range: 30-60
Age Window: 10 years
Effect Size
Lower Window
Upper Window
Significant
Mean Absolute Effect Size
Lower Window Samples
Upper Window Samples
Age
* indicates statistically significant changes (p < 0.05)

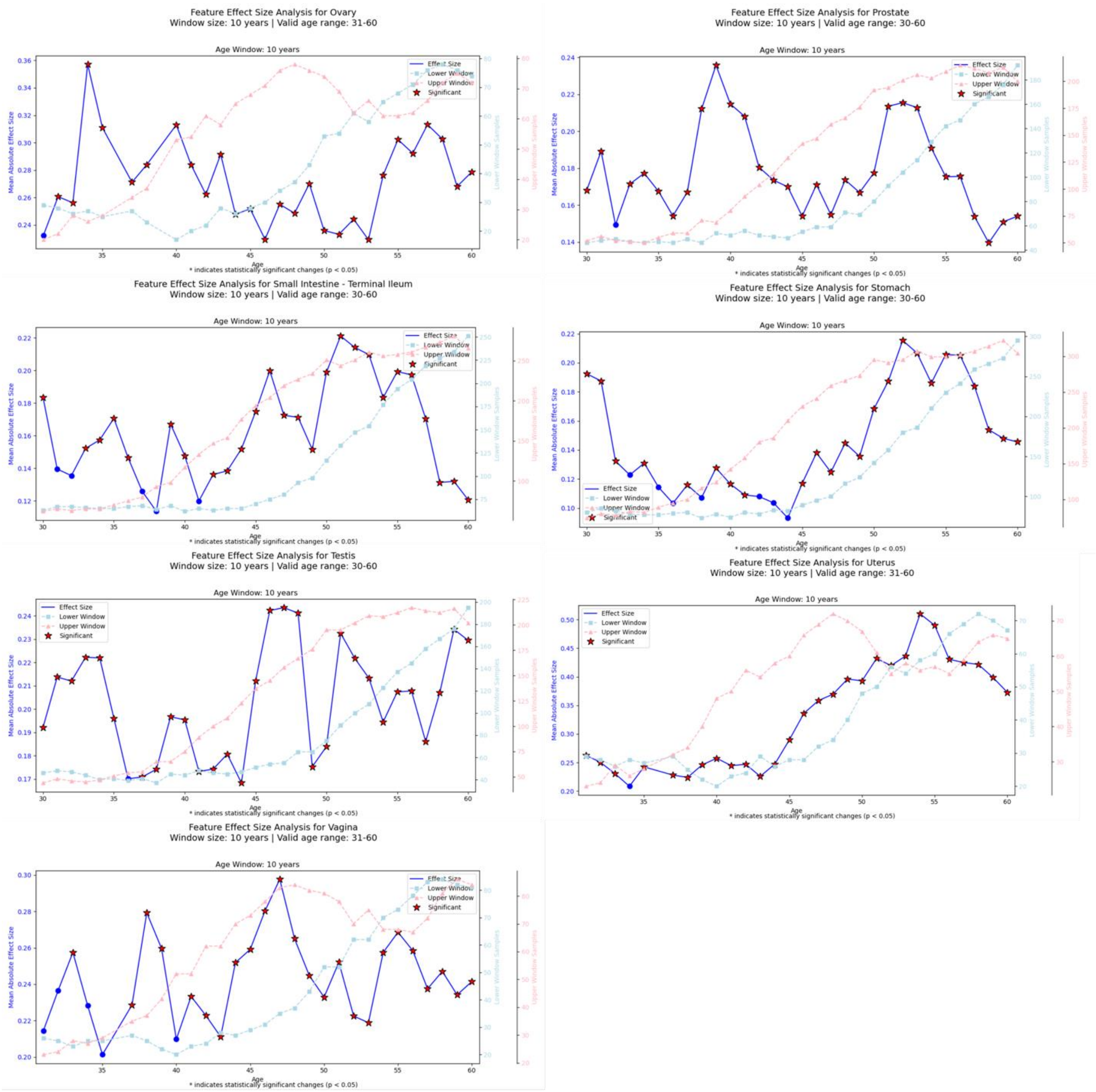


**Figure R8: Raw Structural Aging Trajectories using Virchow 2.** Raw Structural Aging Trajectories (blue lines) are shown for the same 15 tissues. Structural transitions were computed using embeddings extracted from **Virchow 2**, a large-scale histopathology foundation model trained using self-supervised transformer architectures optimized for morphological representation learning. Virchow 2 generates 1280-dimensional patch embeddings, which were aggregated into slide-level representations prior to PathStAR analysis.Red stars denote age transitions where ≥5% of features show nominal significance ($p < 0.05$). Dashed curves display the number of samples contributing to each adjacent age comparison. Replication of tissue-specific structural aging patterns in most tissues using Virchow 2 further demonstrates that the observed non-linear and tissue-specific remodeling trajectories are not artifacts of a particular model architecture but reflect consistent age-dependent structural reorganization detectable across independent foundation models.

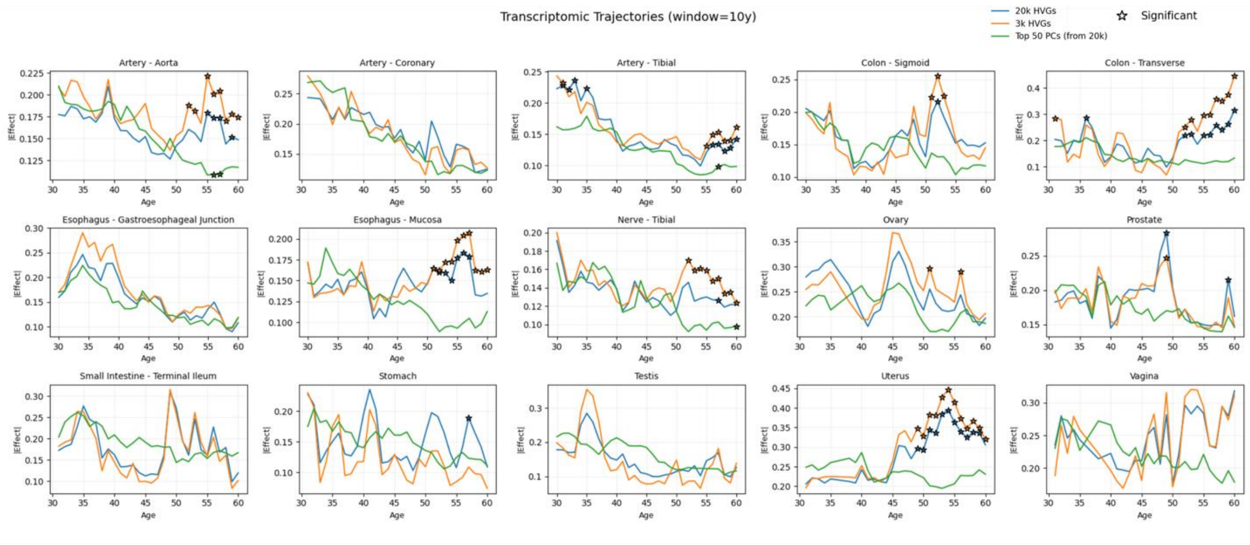


**Figure R9 : Transcriptomic aging trajectories across tissues (window = 10 years).** We computed sliding-window effect sizes across age using three feature spaces per tissue: top 20k highly variable genes (HVGs), top 3k HVGs, and the top 50 principal components (PCs) derived from the 20k HVGs. Across tissues, trajectories derived from PCA (top 50 PCs) show smoother, more consistent age-dependent patterns and clearer significant transitions, indicating that PCA better captures the dominant aging trajectory compared to raw gene-level features.

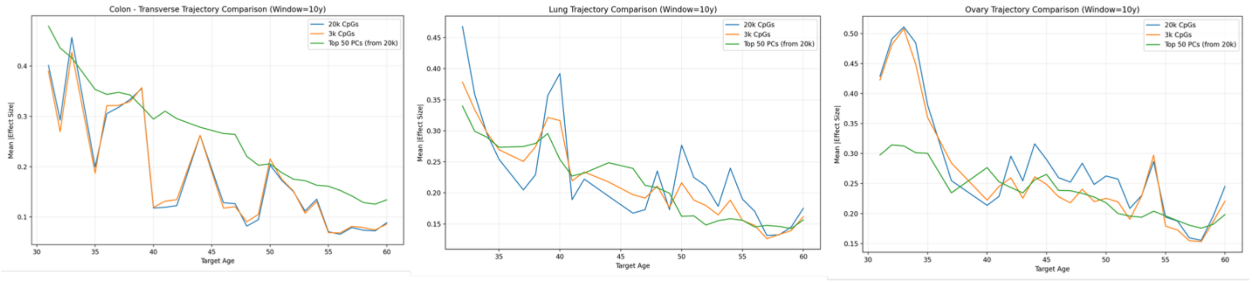


**Figure R10: Methylation aging trajectory comparison (window = 10 years).** Using the same sliding-window framework, we compared trajectories built from 20k CpGs, 3k CpGs, and the top 50 PCs derived from the 20k CpGs. PCA-based trajectories exhibit more stable age trends and reduced noise, suggesting that low-dimensional principal components more effectively capture the underlying methylation aging structure than individual CpG features.
